# Mapping the regulatory effects of common and rare non-coding variants across cellular and developmental contexts in the brain and heart

**DOI:** 10.1101/2025.02.18.638922

**Authors:** Andrew R. Marderstein, Soumya Kundu, Evin M. Padhi, Salil Deshpande, Austin Wang, Esther Robb, Ying Sun, Chang M. Yun, Diego Pomales-Matos, Yilin Xie, Daniel Nachun, Selin Jessa, Anshul Kundaje, Stephen B. Montgomery

## Abstract

Whole genome sequencing has identified over a billion non-coding variants in humans, while GWAS has revealed the non-coding genome as a significant contributor to disease. However, prioritizing causal common and rare non-coding variants in human disease, and understanding how selective pressures have shaped the non-coding genome, remains a significant challenge. Here, we predicted the effects of 15 million variants with deep learning models trained on single-cell ATAC-seq across 132 cellular contexts in adult and fetal brain and heart, producing nearly two billion context-specific predictions. Using these predictions, we distinguish candidate causal variants underlying human traits and diseases and their context-specific effects. While common variant effects are more cell-type-specific, rare variants exert more cell-type-shared regulatory effects, with selective pressures particularly targeting variants affecting fetal brain neurons. To prioritize *de novo* mutations with extreme regulatory effects, we developed FLARE, a context-specific functional genomic model of constraint. FLARE outperformed other methods in prioritizing case mutations from autism-affected families near syndromic autism-associated genes; for example, identifying mutation outliers near *CNTNAP2* that would be missed by alternative approaches. Overall, our findings demonstrate the potential of integrating single-cell maps with population genetics and deep learning-based variant effect prediction to elucidate mechanisms of development and disease–ultimately, supporting the notion that genetic contributions to neurodevelopmental disorders are predominantly rare.

## Introduction

The non-coding regions of the human genome have been recognized as central regulators of gene expression and disease risk, with genome-wide association studies (GWAS) revealing their pivotal role in shaping phenotypes^1–5^. Some regulatory regions are deeply conserved across evolutionary timescales, reflecting stringent selective pressures that preserve their function ^5–10^. Yet, our understanding of how non-coding sequences influence cell type–specific gene regulation across diverse tissues—and how human evolution has shaped genetic variation in these elements—remains incomplete. This gap continues to hinder the translation of genetic associations into evolutionary and mechanistic insights^11–13^.

A major challenge in understanding the non-coding genome is due to context-specificity. During cellular development, regulatory landscapes shift dynamically, with enhancers and promoters becoming active or silenced as cells differentiate, respond to stimuli, or transition between states. Even among seemingly similar cell types—whether in different organs^14,15^, along the same differentiation trajectory^16^, or across the human lifespan^17,18^—unique transcription factor (TF) repertoires and chromatin landscapes dictate how variants exert their regulatory influence. For example, a variant altering transcription factor binding in an adult-specific enhancer might contribute to aging-associated diseases while having minimal impact during earlier stages of development^19^. Single-cell technologies have produced rich data detailing dynamic, context-dependent regulatory programs, but few contexts have scaled to the sample sizes required for powerfully associating genotypes with cellular phenotypes in order to understand disease risk^20,21^.

In parallel, due to their context-specificity, prioritization of non-coding variants relevant to disease remains a significant challenge in human genetics. GWAS have yielded hundreds of thousands of genetic associations, but these signals often include multiple tightly linked common variants. Experimental approaches, such as massively parallel reporter assays^22^ and base editing^23^, offer scalable strategies to assess variant effects but remain constrained by limited throughput and a narrow set of cellular contexts which are experimentally tractable. Computational approaches like statistical fine-mapping or colocalization with QTLs also struggle to resolve single causal variants with high confidence^13,24–26^. Meanwhile, rare non-coding variants, which are abundant in population data, are underpowered to test in GWAS and often ignored^27–31^. Thus, many potentially impactful variants remain unmapped to a specific regulatory function, hindering our ability to translate genetic associations into mechanistic insights and, ultimately, clinical interventions.

Adding to this complexity are evolutionary forces that shape non-coding regions. Selective constraint prevents highly deleterious variants from reaching common frequencies, with GWAS variants enriched in constrained regions and most disease-causing variants expected to be rare. However, the interplay between selective pressures, cell-type-specific regulation, and developmental timing is poorly understood, yet can provide insights into how cell types have evolved or been constrained over human history. For instance, immune-related cell types may be rapidly shaped by positive selection in response to infectious pathogens, while regulatory elements critical to fitness are conserved across evolutionary timescales^32–34^. Understanding how these forces influence genetic variation can provide valuable insights into disease architecture, informing the distribution of allele frequencies affecting disease-relevant cellular contexts.

Recent advances in deep learning models trained on DNA sequences provide a powerful framework for studying context-specific effects of regulatory variants^35–43^. These models learn how relevant motifs and sequence features contribute to functional readouts such as chromatin accessibility, TF binding, transcription initiation, and gene expression. By comparing predictions for DNA sequences differing only at a variant of interest, these models infer a variant’s functional impact without being explicitly trained on specific variants.

Here, we present a novel framework for analyzing regulatory effects of non-coding variants, consisting of ChromBPNet and FLARE. ChromBPNet is a recently developed convolutional neural network model that predicts chromatin accessibility (obtained from ATAC-seq) at base-pair resolution while automatically correcting for enzymatic sequence bias inherent in these experiments^36,43–45^. The model is trained on pseudobulk chromatin accessibility profiles for specific cell types identified from single-cell datasets and predicts the impact of genetic variants by quantifying changes in predicted accessibility between alleles. Furthermore, applying model interpretation tools such as DeepLIFT^46,47^ and TF-MoDISco^48^ allows the identification of critical TF motifs disrupted by a given variant by assessing the contribution of each base in an input sequence to the model’s predicted output. In a cellular context-specific manner, ChromBPNet and its suite of interpretation tools reveals how DNA sequence shapes chromatin accessibility, predicts which variants alter accessibility, and uncovers the TFs affected by these changes, addressing challenges that exist through conventional variant-to-function approaches. Complementing ChromBPNet, we developed FLARE (Functional Lasso Analysis of Regulatory Evolution) to integrate variant predictions with evolutionary pressures to prioritize rare variants with impactful regulatory effects. FLARE captures variants’ regulatory potential across multiple cell types and can be flexibly trained on any cellular context of interest, making it a powerful tool for prioritizing non-coding variants in disease-relevant settings.

In this study, we use ChromBPNet and FLARE to systematically investigate the consequences of both common and rare variants across cell types in the adult and fetal brain and heart. By applying ChromBPNet to GWAS and QTL data, we uncover context-specific regulatory effects underlying genetic associations and integrate these predictions with measures of evolutionary constraint to reveal the critical role of fetal neuron regulation in shaping genomic constraint and allele frequency distributions. FLARE models of the fetal brain further prioritize de novo mutations in autism-affected families, identifying rare, impactful regulatory variants that elude detection by conventional approaches. Together, these tools harness deep learning to predict context-specific variant effects and prioritize variants likely to disrupt regulatory function, addressing major challenges in linking non-coding variation to gene regulation, development, and disease.

## Results

### A resource of predicted regulatory effects across contexts

We created a comprehensive resource of predicted regulatory effects across diverse cellular and developmental contexts by integrating data from the 1000 Genomes Project^49^ (1KG), single-cell ATAC-seq (scATAC-seq) of adult and fetal brain and heart^15,44,45,50^, and cell-type-specific deep learning models of chromatin accessibility using ChromBPNet^43^ (Figure 1A).

**Figure 1:**
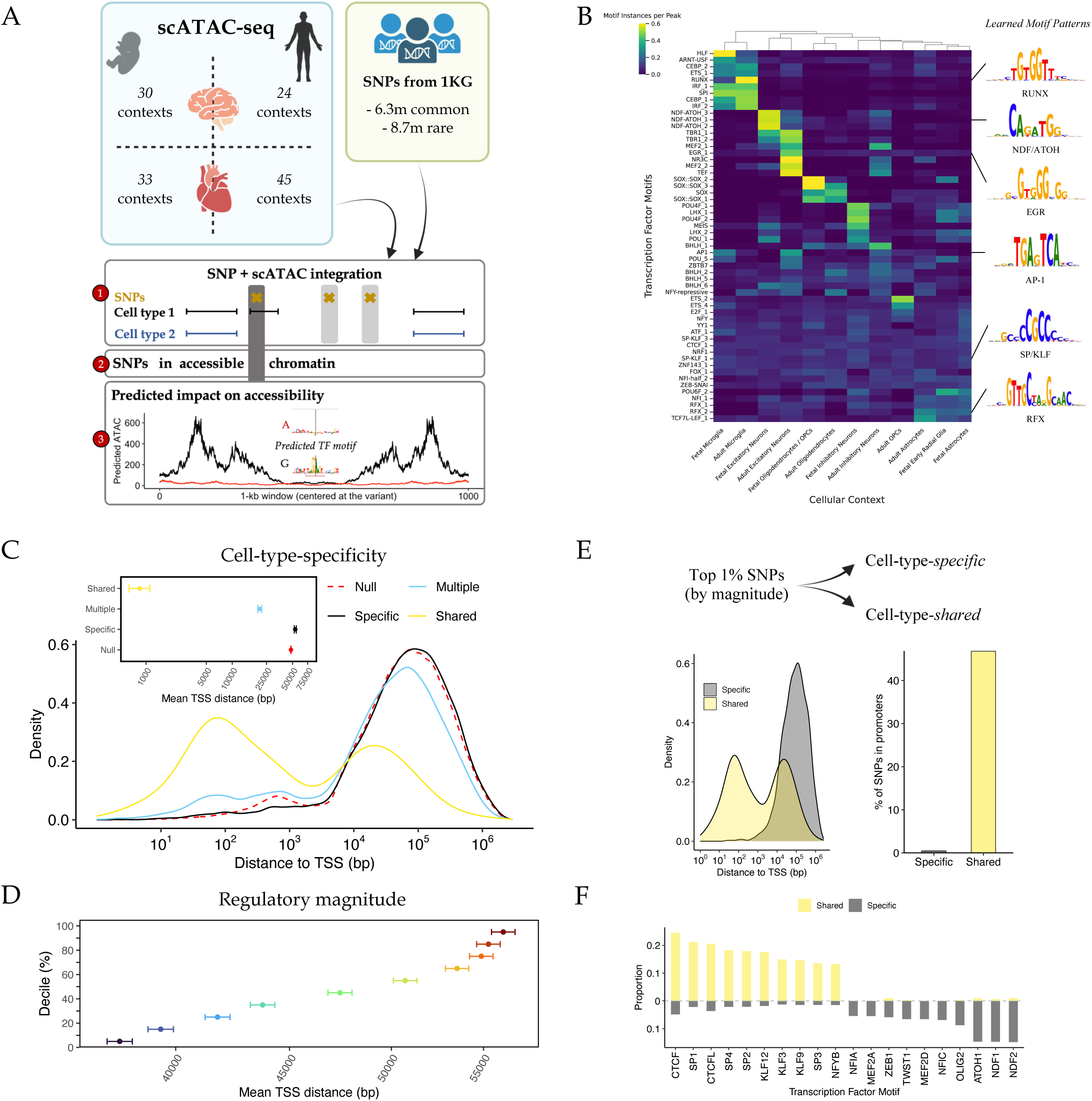
Creation of a resource for context-specific regulatory effect predictions. (A) Overview of the dataset generation process. SNPs were curated from 1KG (6.3 million rare, 8.7 million common), and their effects on chromatin accessibility were predicted using ChromBPNet models trained on scATAC-seq data across 132 contexts. (B) Heatmap of 56 transcription factor (TF) motifs learned across the 54 cell types in the fetal and adult brain. Shown are 1 representative cluster for 12 different cell-type contexts. Values are row-normalized motif instances per peak for each of the 12 representative cell types. The 56 motifs are composed of the union of the top 50 motifs with the greatest variance across the representative cell types and the top 50 motifs with the highest mean value across those cell types. (C) Density plots showing the genomic context of variants (distance to TSS) stratified by their cell-type-specificity. Inset plot shows the mean distance within each group with 95% CI. Variants were categorized as: “null” (impacting 0 cell types), “specific” (impacting only 1 cell type), “shared” (impacting >80% of cell types), and “multiple” (impacting more than 1 but fewer than 80% of cell types). (D) Mean distance to TSS of variants within each decile of regulatory magnitude (the maximum ChromBPNet score across contexts). (E)Large-effect SNPs (top 1% regulatory magnitude) were stratified into cell-type-specific and cell-type-shared sets. Density plot shows distance to TSS of only the large-effect SNPs, and barplot shows the percentage of the large-effect SNPs in promoter regions for specific and shared variant sets. (F) Summary of top motifs disrupted by large-effect cell-type-specific and shared variants.

To generate variant predictions, we first curated a variant dataset from 3,202 whole genomes in 1KG, consisting of 6,349,771 common SNPs (MAF > 5% globally and in the European subset) and 8,757,029 rare SNPs (MAF < 0.1%), totaling 15,106,800 variants. In parallel, we uniformly processed pseudobulk ATAC-seq data from 132 cellular contexts across five studies, including 30 from fetal brain, 24 from adult brain, 33 from fetal heart, and 45 from adult heart, and trained ChromBPNet models for each context (Supplementary Note 1, Supplementary Figure 1, Supplementary Tables 1-2). Model interpretations derived from ChromBPNet using DeepLIFT and TF-MoDISco identified motif instances that influence local chromatin accessibility. For example, in microglia, SPI and IRF motifs were key regulators of accessibility^51^, with differentiation-associated RUNX playing a more prominent role in adult microglia compared to the stem-like HLF, which was more influential in fetal microglia. In neurons, accessibility was governed by motifs such as MEF2^52^, TBR1^53^, NDF^54^/ATOH^55^, and EGR. Notably, EGR, which rapidly increases in expression after birth^56,57^, and AP-1 emerged as specific regulators of chromatin accessibility in adult excitatory neurons, while NDF/ATOH was more specific to fetal excitatory neurons (Figure 1B, Supplementary Table 3). Therefore, ChromBPNet models captured cell-type-specific and developmental-stage-specific motif instances, which we leveraged to predict chromatin accessibility for all 15 million variants across 132 cellular contexts, producing nearly two billion variant-by-context predictions.

We used our dataset to: (1) analyze the distribution of regulatory effects across various cellular contexts, (2) prioritize common variants linked to gene expression and human disease, (3) understand the relationship of cell-type-specific regulatory variation to evolutionary constraint, and (4) develop **FLARE** (Functional Lasso Analysis of Regulatory Evolution) to nominate candidate rare non-coding variants in disease-affected families.

### Variants effects are shaped by genomic context and TF binding

Predicting SNP effects across diverse contexts allows us to investigate the relationship between genomic context (the proximity of a variant to transcribed regions), cell-type specificity (the number of cell types predicted to be affected by a variant), and regulatory magnitude (the maximum predicted effect of a variant on chromatin accessibility across contexts). As variant effects in QTL studies are linked to the distance from the transcription start site^58^ (TSS), we used our ChromBPNet predictions to test whether the distance of variants from the TSS correlates with their cell-type-specificity and regulatory magnitude. We concentrated the analysis on rare variants and fetal tissues to minimize the confounding effects of selective pressures, and calculated cell-type-specificity from ChromBPNet scores using empirical *P*-values derived in each cell type (see Methods).

By comparing with accessible but non-functional variants (*P* > 0.01 in all cell types), we observed that variants with cell-type-specific effects (*P* < 0.01 in only 1 type) tend to be more distal to genes, whereas variants with cell-type-shared effects (*P* < 0.01 in 80% of cell types) are more proximal (Figure 1C, Supplementary Figure 2). Additionally, we observed that cell-type-shared variants exhibited larger magnitude effects than cell-type-specific ones (on average) in multiple sensitivity analyses (Supplementary Figure 2C, Supplementary Note 2). However, variants with stronger effects tend to be more distally-located to the TSS, suggesting a complex interplay between genomic context, cell-type-specificity, and regulatory magnitude (Figure 1D).

To further understand this interplay, we analyzed the top 1% of variants with the strongest predicted effects. We hypothesized that the largest-effect variants are more likely to exhibit effects in multiple cell types due to their roles in promoter regions or disruption of ubiquitous TF motifs shared across cell types. To test this, we focused on understanding the role of promoters and putative transcription factor binding in shaping cell-type-*shared* versus cell-type-*specific* variant effects. Compared to specific variants, shared variants were closer to the TSS with 46.9% located in experimentally-verified promoter regions^59^ versus only 0.5% of specific variants (Figure 1E). This mix of distal enhancer variants and proximal promoter variants underlies the bimodal distribution of TSS distances for shared variants (Figure 1C). Additionally, shared variants disrupted motifs for broadly expressed TFs, such as those from the CTCF or SP/KLF families, while specific variants primarily disrupted lineage-specific TFs, including ATOH1, NDF1, NDF2, and OLIG2, which regulate neuronal and oligodendrocyte differentiation (Figure 1F, Supplementary Table 4). This disruption of ubiquitous TF motifs likely explains why shared variants, independent of their proximity to the TSS, tend to exhibit stronger regulatory effects, highlighting the intricate relationship between cell-type-specificity and regulatory magnitude.

### Context-specific models reveal regulatory effects of fine-mapped eQTLs

We next used ChromBPNet to prioritize fine-mapped variants and their context-specific effects underlying expression quantitative trait loci (eQTLs) from the GTEx Project^3^, which are purely based on statistical modeling of population genetic data and are derived from bulk RNA-seq data in adult tissues. First, we analyzed eQTLs identified in GTEx brain samples. Within fine-mapped regions, we found that high-confidence eQTLs (posterior inclusion probability, PIP > 0.9) exhibited more cell-type-shared regulatory effects across adult brain contexts than did lower-confidence variants in the same regions (PIP < 0.01) (*P* < 10^-15^). A similar trend emerged for fine-mapped heart eQTLs from GTEx using adult heart ChromBPNet models. The number of affected cell types in adult heart and adult brain models had the strongest enrichment for fine-mapped eQTLs in the respective GTEx tissue contexts, outperforming mismatched models such as heart models applied to brain eQTLs, or fetal models applied to adult-derived eQTLs, as expected (Supplementary Figure 3).

We next asked how the predicted regulatory effects of fine-mapped QTLs vary across cellular contexts. We examined individual cell types within the adult brain, evaluating their ChromBPNet scores at eQTL loci. Compared to non-fine-mapped eQTLs, fine-mapped eQTLs showed significantly elevated ChromBPNet scores in accessible regions, with stronger enrichment in adult brain contexts than in fetal brain, and weaker trends observed in heart tissues (see Methods). Applying the same approach to fine-mapped eQTLs from GTEx heart and artery tissues, we found that ChromBPNet models trained on heart cell-types displayed a much stronger enrichment compared to brain models, and that adult heart contexts outperformed fetal heart contexts (Figure 2A, Supplementary Table 5). Thus, context-specific models prioritize context-relevant QTLs.

**Figure 2:**
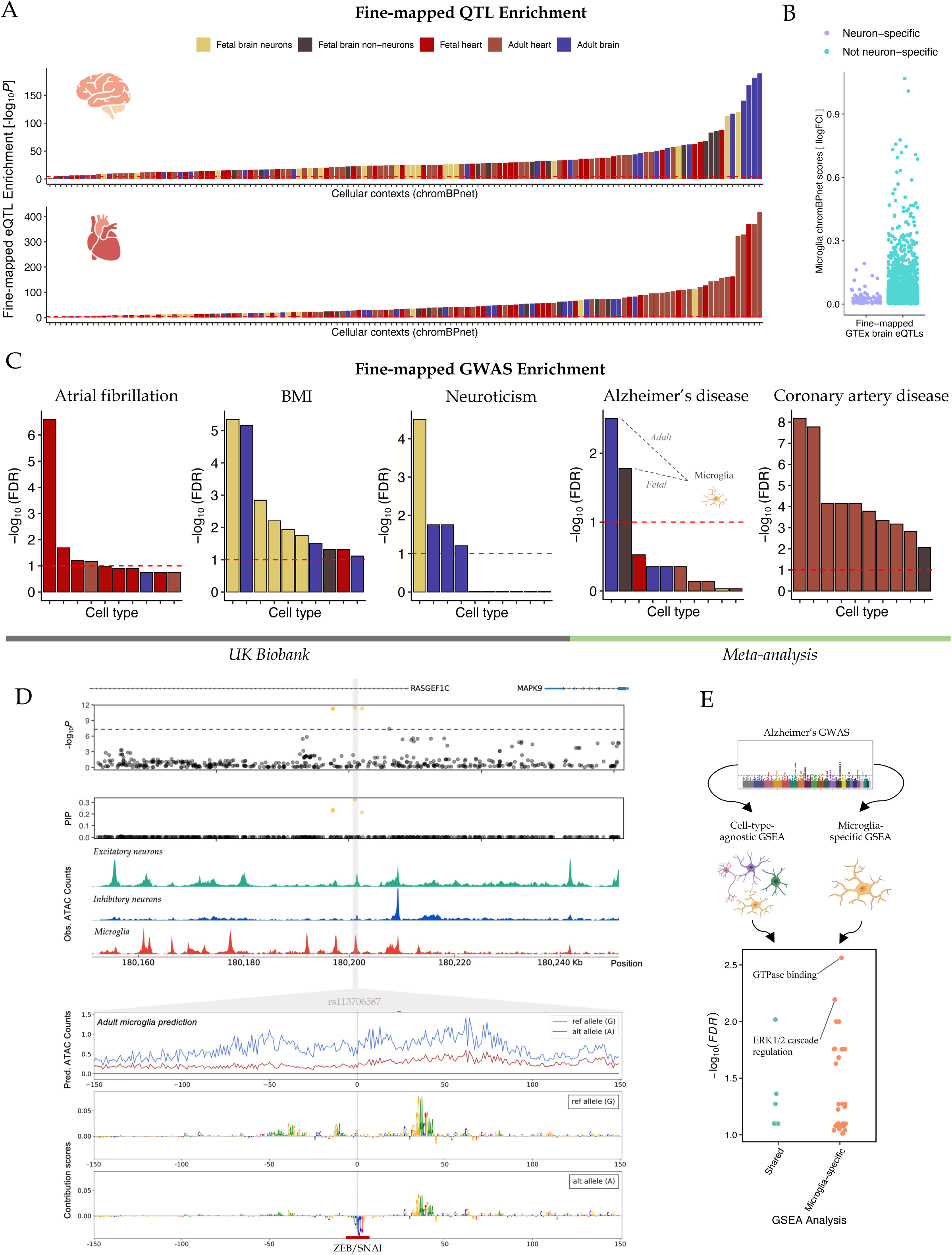
ChromBPNet prioritizes variants influencing gene expression and disease risk. (A) Enrichment of fine-mapped eQTLs (relative to non-fine-mapped eQTLs) using ChromBPNet in different cell types. Top panel shows all GTEx brain eQTLs, and bottom panel shows all GTEx heart and artery eQTLs. Only accessible variants were tested. The red line indicates the Bonferroni-corrected significance threshold. Bars colored by developmental and organ context. “Adult brain” includes neuronal, glial, and other cell types; “Fetal brain non-neurons” include glial and other cell types (e.g. pericytes, vascular endothelial cells). (B) Distribution of microglia ChromBPNet scores (in |log fold-change|) for fine-mapped eQTL variants, stratified by neuron-specific and non-neuron-specific eQTLs. (C) Similar to (A), enrichment of fine-mapped GWAS loci across 5 brain and heart-associated phenotypes. The red line indicates *FDR* = 0.1. Grey bar under the plots indicate fine-mapped variant sets derived from GWAS in UK Biobank, while the green bar under the plots indicate those derived from GWAS meta-analyses. (D) A 100 kb region surrounding an Alzheimer’s disease GWAS locus in a *RASGEF1C* intron. GWAS -log_10_ *P*-values are shown along the *y*-axis in the top panel for all tested variants. Variants in credible set are shown with orange points, with their fine-mapped posterior inclusion probability (PIP) values displayed in the panel below. Normalized scATAC–seq-derived pseudobulk tracks for excitatory neurons, inhibitory neurons, and microglia are shown next. Finally, ChromBPNet predictions in microglia are shown for rs113706587 in a 300 bp window surrounding the variant, with predicted profiles for the reference and alternate alleles shown in blue and red respectively. DeepLIFT contribution scores for each base in the two input sequences (containing either the reference or the alternate alleles) are shown in the bottom two panels. The putative causal motif is highlighted. (E) Gene set enrichment analysis (GSEA) was performed for both GWAS variants with cell-type-specific effects in microglia and GWAS variants genome-wide. The FDR significance displayed for each gene set in the microglia-specific analysis (shown along the *y*-axis) are stratified by whether the gene sets also appear in a cell-type-agnostic GSEA.

We noted the association of ChromBPNet models across diverse contexts with fine-mapped bulk eQTLs (Figure 2A, Supplementary Figure 3), which likely reflects a combination of shared regulatory architectures, conserved TF binding motifs, and similar cell types that are present across tissues. However, when we focused on bulk brain eQTLs, we discovered evidence that they could be attributed to specific cell types: ChromBPNet scores from adult microglia were significantly reduced at fine-mapped brain eQTLs showing a neuron-specific effect^60^ (*P* = 3.5 × 10^-5^) (Figure 2B; see Methods), while adult neuron ChromBPNet scores were not significantly different (*P* > 0.05). Furthermore, we found that fine-mapped eQTLs from bulk brain RNA-seq were more likely to be driven by cell-type-shared variants, such as a two-fold enrichment by variants with effects in both adult microglia and excitatory neurons compared to microglia and neuron-specific variants genome-wide (*P* = 0.037). These findings highlight the need for context-specific models for predicting regulatory variant effects and demonstrate the dilution of context-specific eQTLs using bulk RNA-sequencing.

### Pinpointing disease-relevant variants using cell-type-specific chromatin models

We next utilized our resource to prioritize candidate causal GWAS variants underlying human traits and disease. First, we tested whether GWAS heritability of 8 common cardiovascular and neurological phenotypes were concentrated within open chromatin regions in our single-cell ATAC-seq datasets^61,62^, finding 283 significant cell-type-by-disease combinations (*FDR* < 0.1) (Supplementary Figure 4A, Supplementary Table 6). For example, in Alzheimer’s disease^63^, SNP-heritability was specifically enriched in microglia, with adult microglia showing stronger enrichment than fetal microglia, suggesting that genetic effects continue to manifest later in life (Supplementary Figure 4B). For anorexia nervosa, heritability was primarily enriched in fetal neurons (to a greater extent than in adult) (Supplementary Figure 4C), aligning with a gene-based analysis for anorexia nervosa risk which identified “positive regulation of embryonic development” as the only significant pathway and emphasizing the role of developmental processes in etiology^64^.

Next, we applied ChromBPNet to fine-mapped GWAS data from the UK Biobank^65^, distinguishing disease-associated fine-mapped variants (PIP > 0.2) from non-fine-mapped variants (PIP < 0.01) within the same loci, focusing exclusively on teasing apart putative variant causality in accessible regions (see Methods). We found that ChromBPNet models trained on brain contexts effectively prioritized causal variants linked to body mass index (BMI) and neuroticism risk. In contrast, models derived from heart contexts prioritized variants associated with atrial fibrillation. Notably, brain-context models were not informative for variants related to atrial fibrillation, and heart-context models were not informative for BMI or neuroticism (Figure 2C). Furthermore, we extended our models to meta-analyses of Alzheimer’s disease and coronary artery disease^63,66,67^. Using SuSiE to fine-map the GWAS summary statistics^68^, we found that microglia-specific ChromBPNet models effectively identified fine-mapped variants for Alzheimer’s disease, with a stronger enrichment in adult microglia compared to fetal microglia, consistent with heritability enrichments (Figure 2C). Conversely, heart-derived models demonstrated superior performance in identifying fine-mapped variants associated with coronary artery disease compared to brain contexts (Figure 2C), with correlated cell-type enrichments between fine-mapped variant sets from two distinct published GWAS (Supplementary Figure 5). These findings underscore the context-specific applicability of ChromBPNet models in pinpointing causal regulatory variants relevant to different diseases (Supplementary Table 7).

### Microglia-driven mechanisms of Alzheimer’s disease risk

Using our predictions of non-coding Alzheimer’s disease GWAS loci, we prioritized candidate causal variants with potential effects on chromatin accessibility in adult microglia. For example, statistical fine-mapping identified a credible set of four susceptibility variants (each with PIP > 0.2) in an intron of *RASGEF1C*, which activates small GTPases by catalyzing the exchange of GDP for GTP, which are also associated with decreased *RASGEF1C* expression in GTEx brain tissues^3^. Integrating chromatin accessibility data^44,50^, we found that only two of these variants resided within accessible chromatin peaks in microglia. Of these, our model predicted that only rs113706587 (chr5:180201150:G:A) impacted chromatin accessibility. Using DeepLIFT to interpret the models and derive predictive contribution scores for each base in the local sequence, we found that the alternate allele creates a ZEB/SNAI motif that represses accessibility^69^. Consequently, the ChromBPNet model predicted overall reduced accessibility in the region. In contrast, rs116347724 (chr5:180197032:G:A), while also located within a microglial peak, was not predicted to influence accessibility (Figure 2D). These findings prioritize rs113706587 as the candidate causal variant at this GWAS locus and nominate its regulatory mechanism in microglia. Additionally, evidence converged on multiple regulatory variants at a non-coding GWAS locus near *PICALM*, challenging the conception of a single underlying causal variant^70–75^ (Supplementary Note 3, Supplementary Figure 6a).

To understand the downstream consequences of microglia-acting variants in Alzheimer’s risk, we partitioned pathways specifically acting in microglia by identifying 36 variants at GWAS loci with microglia-specific effects and performing gene-set enrichment analysis (GSEA, see Methods). We compared GSEA results from candidate genes near GWAS variants with predicted effects in microglia to the results from all candidate genes of GWAS variants (ignoring ChromBPNet predictions). While many identified gene sets are not unique to studying cell-type-specific genes (as compared to the cell-type-agnostic GSEA; Supplementary Figure 6), we found a number of pathways with microglia-specific enrichments, such as GTPase binding (FDR = 0.003, involving SNPs linked to *USP6NL, ABCA1, ATG16L1, RASA1, RIN3,* and *PICALM*) or ERK1/2 cascade regulation (FDR = 0.006, involving SNPs linked to *APP, FLT4, PTK2B, ADRA1A, SCIMP,* and *FERMT2*) (Figure 2E, Supplementary Tables 8-9). These microglia-driven mechanisms have been previously linked to microglia function and immune responses in Alzheimer’s pathogenesis^76,77^, but would be challenging to detect using GSEA without cell-type-specific information. Therefore, by integrating (1) statistical fine-mapping, (2) chromatin accessibility profiling in a disease-relevant cell type, and (3) deep learning-enabled functional effect predictions in the same cell type, we provide a framework for dissecting non-coding GWAS loci and further understanding disease mechanisms (Supplementary Table 8-9).

### Ultra-rare variants show larger and more shared regulatory effects than common variants

While GWAS associates common variants to disease, studying rare variants is challenging due to limited statistical power. Since ChromBPNet can be applied to any base pair in the genome, we used our model to investigate the regulatory effects of ultra-rare variants^78^ (MAF < 0.1%). We first compared their effects to common variants (MAF > 5%) to explore the selective pressures shaping their frequencies in human populations. Purifying selection acts to eliminate deleterious variants from the population, keeping them at ultra-rare frequencies. Conversely, genetic drift allows some variants to rise to common frequencies through chance. This interplay between selection and drift helps explain why certain variants remain rare, while others drift to higher frequencies over generations, but it is unclear how regulatory effects in particular cell types influence evolutionary forces (Figure 3A).

**Figure 3:**
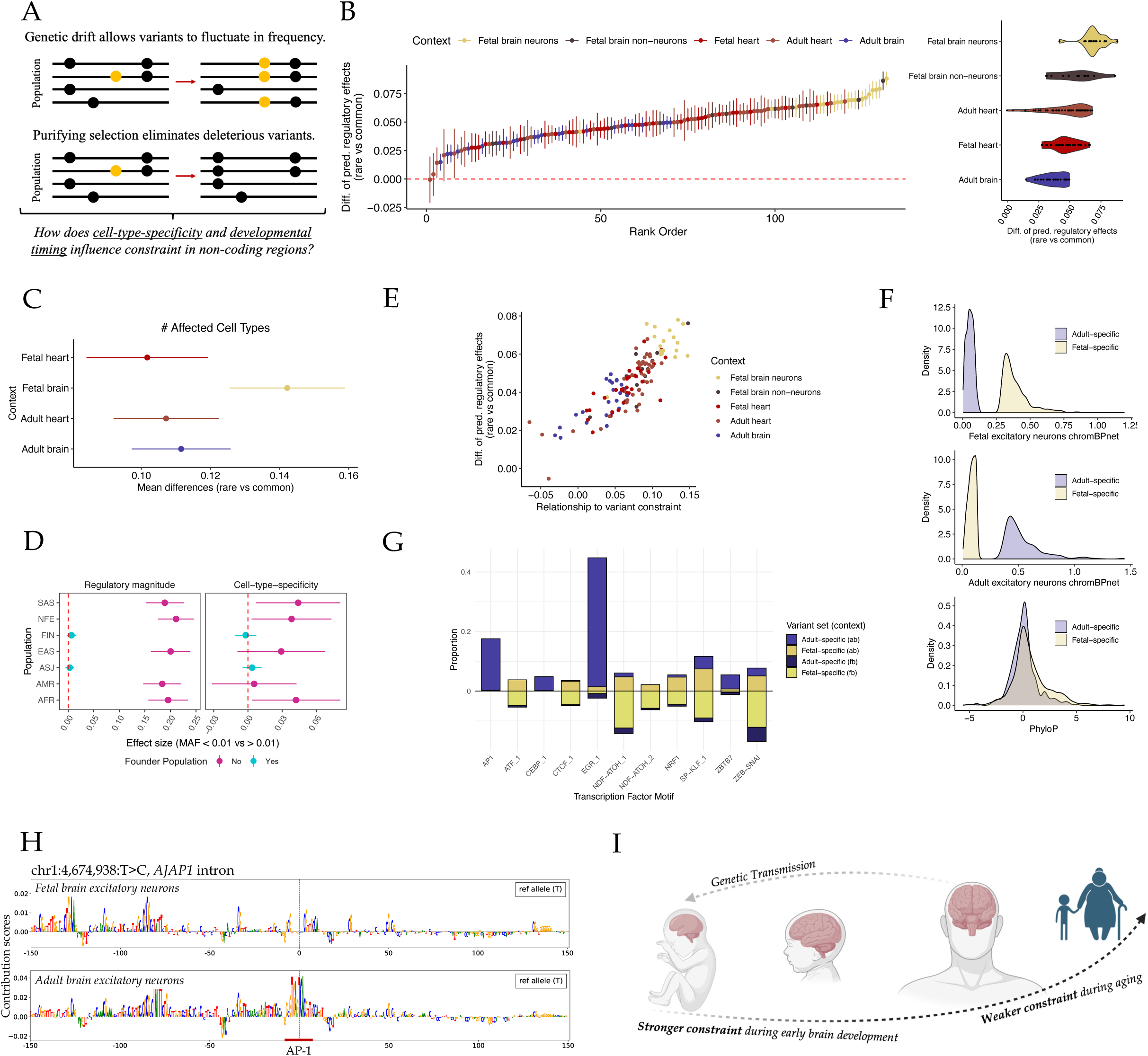
Regulatory effects in fetal neurons shape evolutionary pressures. (A) A schematic illustrating the relationship between constraint and allele frequencies, highlighting the critical challenge of deciphering the role of cell-type-specific regulation in evolutionary pressures. (B) Left, effect sizes (*y*) estimating the mean difference in regulatory effects (ChromBPNet) between common and ultra-rare variant sets, tested for each cell type individually (sorted along the *x* axis, and colored by broad context). Estimates greater than zero indicate larger mean effects in the rare variant set. Error bars show 95% CI. Right, violin plots showing effect sizes stratified by broad context. (C) Estimated differences (*x*) in the number of affected cell types between ultra-rare and common variants, stratified by developmental and organ contexts (*y*). The effect sizes (*x*) reflect standardized mean differences with 95% CI, and are derived from a linear model accounting for nearest gene constraint and TSS distance. Estimates are in standard deviation units, with greater than zero indicating greater cell types affected on average in the ultra-rare set. (D) Left, effect sizes (*x*) with 95% CI estimating the mean difference in regulatory magnitude (maximum ChromBPNet score across all contexts) and right, number of affected cell types between rare variants in 1KG EUR with MAF < 0.01 and those with MAF > 0.01 in 7 gnomAD populations. Points colored by whether it is a founder population or not. “AFR” (African), “AMR” (Latino/Admixed American), “ASJ” (Ashkenazi Jewish), “EAS” (East Asian), “FIN” (Finnish), “NFE” (Non-Finnish European), and “SAS” (South Asian), with “ASJ” and “FIN” reflecting founder populations. (E) Scatterplot showing the relationship between the mean difference in regulatory effects (*y*) between common and ultra-rare variant sets (from the common vs. ultra-rare ChromBPNet analysis) and the ! coefficient (*x*) from a linear model correlating ChromBPNet scores with sequence conservation (PhyloP) across rare variants. Both analysis models account for nearest gene constraint and TSS distance. (F) Density plots displaying fetal excitatory neuron ChromBPNet scores, adult excitatory neuron ChromBPNet scores, and PhyloP for fetal-specific and adult-specific excitatory neuron variants. (G) Proportion of adult-specific or fetal-specific variants that overlap motifs in adult excitatory neurons (ab, top) or fetal excitatory neurons (fb, bottom), as predicted using ChromBPNet models. (H) DeepLIFT contribution scores for the reference allele of an adult-specific variant in adult and fetal excitatory neurons. The scores indicate that AP-1 drives accessibility in adult but not fetal neurons. The alternate T allele is predicted to disrupt the AP-1 binding. (I)Early brain development is under strong purifying selection, reflecting heightened sensitivity to damaging genetic variants particularly in fetal neuronal regulatory elements. Selective pressures decrease with aging, such that variants with large effects later in life (particularly after genetic transmission to future generations) will often escape selection. As a result, the major genetic influences for rare neurodevelopmental disorders are expected to be rare since deleterious variant effects during development will be strongly selected against.

First, for each cell type individually, we compared the regulatory effects (as predicted by ChromBPNet) of accessible common versus ultra-rare variants, controlling for genomic context (nearest gene constraint and TSS distance) and focusing on variants in open chromatin regions. Across nearly all cellular contexts, ultra-rare variants exhibited larger regulatory effects than common variants (Figure 3B). The differences were particularly pronounced in fetal brain neurons, with a 93% greater increase in fetal excitatory neurons than the average cell type (Figure 3B, Supplementary Figure 7) that remained stronger than other cell types when restricting the common vs. ultra-rare analysis to variants accessible in a second cell type for comparison (Supplementary Note 4, Supplementary Figure 8). We similarly observed that ultra-rare variants were more likely than common variants to be accessible in fetal brain neurons relative to other cell types (Supplementary Note 5, Supplementary Figure 7, 9). Not every neuron type in the fetal brain displayed this enrichment, as limbic system neurons had smaller effect differences than other cell types. Furthermore, immune cells exhibited notably smaller differences across developmental organ contexts (*P* = 1.0×10^-8^), suggesting reduced regulatory constraint in immune contexts (Supplementary Figure 7). This may facilitate the emergence and rise in frequency of new regulatory mutations, enabling adaptation to environmental cues and enhancing human survival against pathogens over evolutionary time (Supplementary Table 10).

In addition to larger regulatory effects, we also observed differences in the cell-type-specificity of these effects between ultra-rare and common variants. We evaluated cell-type specificity of accessible common versus ultra-rare variants, focusing on those associated with at least one cell type. Our analysis found, across development and organ contexts, that ultra-rare variants impacted a greater number of cell types on average compared to common variants. This suggests that ultra-rare variants tend to have more cell-type-shared effects, while common variants tend to exhibit more cell-type-specific effects. Further, this trend was more pronounced in the fetal brain than in the adult brain or heart contexts, reinforcing that mutations that affect fetal brain development are subject to stronger constraints and are less likely to drift to common frequencies (Figure 3C).

We replicated these trends by examining ultra-rare variants in European 1KG that reached higher frequencies in global gnomAD populations (due to genetic drift or resampling). These variants with higher frequencies in other global populations exhibited smaller and more cell-type-specific regulatory effects compared to those that remained rare. In contrast, ultra-rare variants that were common in founder populations did not follow this pattern. Deleterious variants are often found at higher population frequencies in founder populations, such as Ashkenazi Jews or Finns, due to historical population bottlenecks and isolation, genetic drift, and limited gene flow, which supports that purifying selection present in global populations has not manifested in the same manner for founder populations (Figure 3D).

### Fetal neurons shape selective constraint in non-coding regions

To gain further support for the observations made in our rare versus common variant analysis, we computed sequence conservation (using PhyloP conservation scores^79^ derived from alignment of 100 vertebrate species) for our set of 8,757,029 ultra-rare SNPs (MAF < 0.1%). We then correlated this measure of selective constraint with regulatory function (using ChromBPNet) at accessible ultra-rare variants, individually for each cell type, accounting for genomic context (see Methods). Stronger correlations indicate selective pressures that act preferentially in certain cellular and developmental contexts.

Overall, we found that ChromBPNet scores significantly correlated positively with constraint across the majority of cell types, and that the strongest correlations were observed at fetal brain neurons (Supplementary Figure 11-12). Indeed, both the rare vs common variant analysis and the constraint vs regulatory effects across rare variants both aligned at similar results, with strong correlations across cell types when comparing the two analyses and prioritizing fetal neurons as the most constrained cellular contexts analyzed (Figure 3E, Supplementary Table 10).

Finally, we curated ultra-rare variants accessible in both adult and fetal excitatory neurons. Variants with effects during fetal development often had an effect in adult excitatory neurons and vice versa (OR = 68.5, *P* < 10^-308^). However, we identified a subset of 381 adult-specific and 1236 fetal-specific excitatory neuron variants within the top and bottom 10% of differences between fetal and adult ChromBPNet scores and with significant effects in one development context but not the other (see Methods). Testing for associations between constraint with fetal- or adult-specific variants revealed that fetal-specific variants have significantly increased constraint on average compared to genome-wide co-accessible variants (∼0.55 SD increase, *P* = 6.1×10^-84^), whereas adult-specific variants did not (*P* > 0.05), with significant constraint differences observed directly between fetal- and adult-specific variants (*P* = 6.3×10^-9^) (Figure 3F). This further supports that genetic effects occurring early in brain development have been tightly regulated throughout human evolution.

To explore transcription factors driving development-specific regulatory differences and the evolutionary forces shaping them, we compared disrupted motifs between adult-specific and fetal-specific variants. First, we used context-specific model interpretations from ChromBPNet (Figure 3G, Supplementary Table 11). We predicted that 43.3% and 17.3% of adult-specific variants disrupted an EGR motif or AP-1 motif in adult neurons respectively (compared to 1.5% and 0.02% of fetal-specific variants). Furthermore, disrupting these motifs was not predicted to have strong effects on fetal neuron accessibility; in fetal neurons, 1.6% and 0% of adult-specific variants were predicted to disrupt EGR and AP-1 binding, illustrating the context-specificity of regulatory variant mechanisms as EGR and AP-1 are less frequent drivers of chromatin accessibility in fetal neurons (Figure 1B, Figure 3H, Supplementary Figure 12, Supplementary Note 6). Fetal-specific variants altered a diverse set of motifs, including the fetal-specific NDF/ATOH (Figure 1B), as well as SP/KLF, ZEB/SNAI, NRF1, and ATF, which are also relevant in the adult context (Figure 3G, Supplementary Figure 13). To further support the findings based on ChromBPNet, we used position weight matrices via motifbreakR^80^ and TF binding predictions via SNP-SELEX^81^. AP-1 and EGR1 motifs were enriched among adult-specific variants, while SP/KLF, NRF1, and ATOH1 motifs were predominantly disrupted in fetal-specific variants, with adult-specific variants predicted to disrupt EGR1 binding and fetal-specific variants predicted to disrupt ATOH1 binding (Supplementary Note 7, Supplementary Figure 14, Supplementary Tables 12-13). Overall, ChromBPNet highlighted development-specific, predictive motif instances that enable interpreting non-coding variant effects across development. Furthermore, our results indicate that in the adult brain, activity- and stress-responsive TFs (AP-1, EGR) become more critical, whereas during fetal development, neuronal differentiation factors (e.g. ATOH) along with key regulatory proteins (e.g. SP/KLF) shape the regulatory landscape.

In summary, our comparative analysis of rare and common variants reveals that ultra-rare variants tend to have larger regulatory effects and more cell-type-shared functions, particularly in the fetal brain, and this is driven by impacts on specific TFs. Therefore, ultra-rare variants are more likely to impact early developmental processes, while common variants impact more cell-type-specific regulatory functions; ultimately, this supports the evidence that the genetic contribution to rare neurodevelopmental disorders would be expected to be overwhelmingly driven by rare variants^82^ (Figure 3I).

### FLARE: a functional genomic model of constraint

While our results indicate that variants disrupting early brain development are more likely to impact fitness, many new mutations that arise will not influence traits or disease risk. Overall, it remains a major challenge to implicate the specific rare non-coding variants that are disease-causing and fitness-reducing among the ones that have a predicted impact on molecular functions. To address this, we developed **FLARE** (Functional Lasso Analysis of Regulatory Evolution), a lasso regression model that integrates deep learning predictions with evolutionary conservation. Evolutionary conservation is a key predictor of disease risk, influenced by diverse variant mechanisms. FLARE aims to predict PhyloP conservation scores using TSS distance, nearest gene constraint, peak overlap, ChromBPNet scores, and ChromBPNet scores conditional on the variant residing within a peak. Since PhyloP scores are, by definition, not context-specific, we expect FLARE to model the relationship between genomic context, predicted regulatory effects, and evolutionary conservation specifically in cell contexts where regulation is highly relevant to conservation. Thus, FLARE (i) disentangles the contributions of context-specific accessibility and regulatory effects to conservation, (ii) provides an intuitive framework for integrating multiple functional genomic features into a unified model, and (iii) captures variants’ regulatory potential across multiple cell types through its predictions. A powerful advantage of FLARE is that the model can be trained on any context of interest. We therefore trained FLARE models in several different tissues and developmental contexts across 8,757,029 rare variants. Regularization hyperparameters were tuned with 4-fold cross-validation. For each chromosome, FLARE scores are predicted from a model trained on the remaining chromosomes (Figure 4A, Supplementary Figure 15).

**Figure 4:**
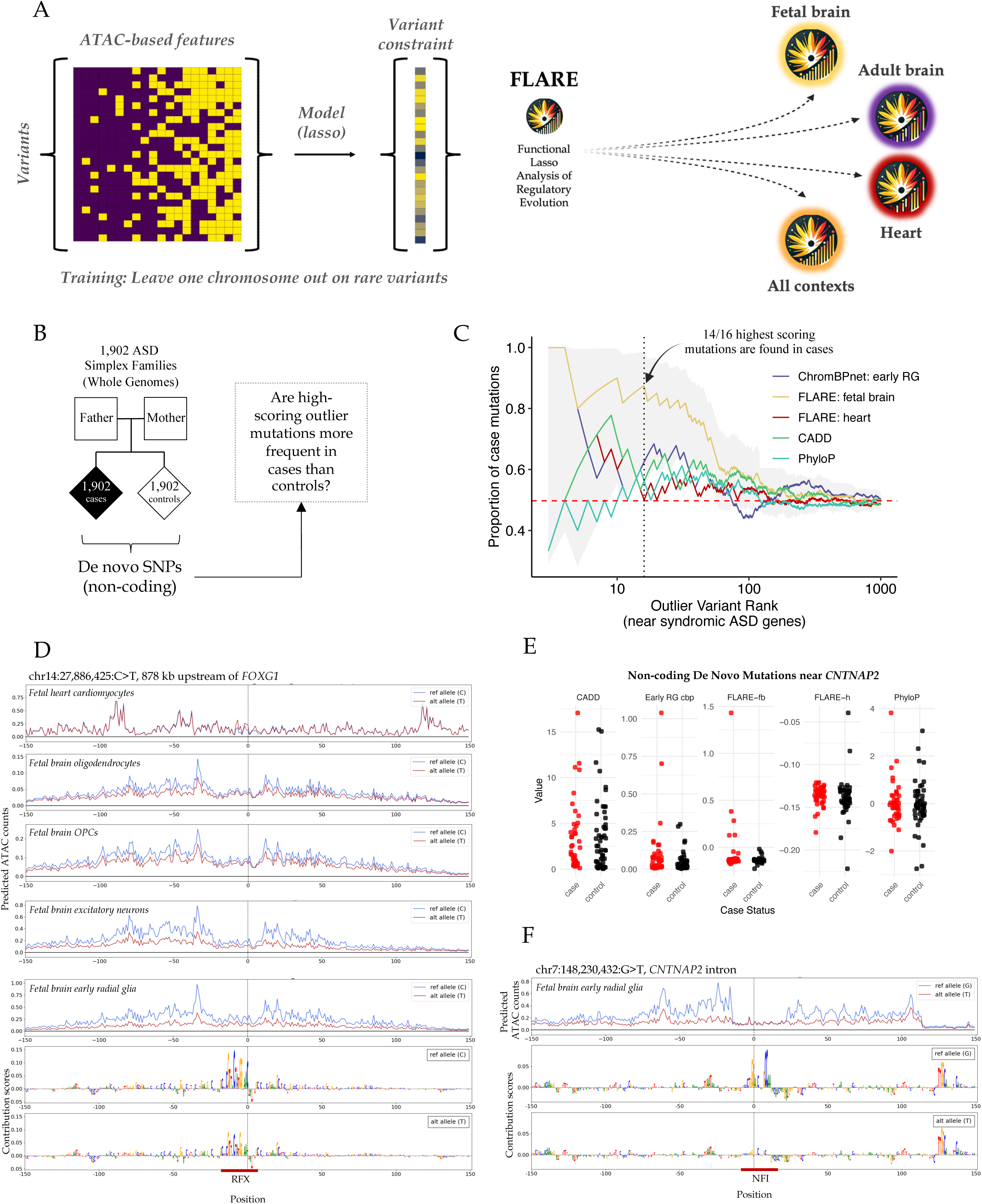
FLARE prioritizes de novo non-coding mutations in autism families. (A) Overview of the FLARE framework, which predicts sequence conservation (PhyloP scores) using regulatory and genomic features for any context of interest. (B) Schematic of de novo mutation prioritization in simplex families containing children with ASD and their unaffected siblings. Outlier mutations represent those with the highest scores. (C) Prioritization of non-coding de novo mutations near syndromic ASD genes using multiple metrics. The *x*-axis displays the rank value for each metric, where 10 would indicate the 10th highest scoring variant according to that metric. The *y*-axis displays the proportion of mutations found in cases in the set of variants with higher scores than the rank (*x*). “ChromBPNet: early RG” refers to the ChromBPNet score for early radial glia in the fetal brain. (D) ChromBPNet predictions for a high-scoring de novo mutation (chr14:27,886,425:C>T) near *FOXG1*. Predictions are shown for a 300 bp window surrounding the variant, with predicted profiles for the reference and alternate alleles shown in blue and red respectively. The variant disrupts an RFX3 motif, reducing chromatin accessibility in many cell types from the fetal brain, but not in the heart (top panel shows the predictions for cardiomyocytes). Motif contribution scores are related to fetal brain radial glia. (E) Values of de novo mutations near *CNTNAP2* for five scoring metrics, stratified by case and control status. Each subplot has a unique axis to display the distribution of each metric. “Early RG cbp” refers to the ChromBPNet score for early radial glia in the fetal brain. (F) ChromBPNet predictions for a high-scoring de novo mutation (chr7:148230432:G>T) in a *CNTNAP2* intron, with the DeepLIFT contribution scores shown in the bottom two panels along with highlighting the putative causal motif.

In the rare variant set, FLARE-fb (fetal brain) outperformed the adult brain (FLARE-ab) and heart (FLARE-h) models by over two-fold in terms of *R*^2^ between predicted and observed PhyloP scores (Supplementary Figure 15). FLARE-fb also showed an improvement of 32.5% over a peak-only model (FLARE-fb-peaks), demonstrating the added value of including sequence-based predictions. Models trained on (i) adult and fetal brain contexts or (ii) all contexts together performed similarly to FLARE-fb (0.2% and 1% improvements respectively), further supporting the fetal brain as a key context shaped by evolutionary constraint. We replicated these trends using additional variant sets, finding that FLARE models informed context-specific regulatory evolution (Supplementary Figure 15, Supplementary Note 8).

### FLARE prioritizes de novo non-coding mutations in autism

ASD-associated genes are predominantly expressed in the brain^83,84^ and show higher levels of evolutionary constraint^85^. Previous studies have established that *de novo* mutations in coding^84^ and promoter^86^ regions contribute to ASD (for the latter, specifically those in conserved sequences with known TF motifs^86^), but linking *de novo* mutations in distal non-coding regions to ASD remains a challenge^85–88^. We hypothesized that FLARE could prioritize rare non-coding variants under increased constraint, specifically those affecting regulatory elements in the developing brain, by predicting conservation of rare variants using each cell context-specific FLARE model. Consequently, FLARE should be able to capture each variant’s regulatory potential across multiple cell types and their relevance to conservation.

Using FLARE-fb predictions of de novo non-coding mutations from ASD families, we focused on 1,855 variants near genes linked to syndromic forms of autism, which occur as part of a broader genetic syndrome. These syndromes often involve a range of developmental issues, including intellectual disabilities, physical abnormalities, and other neurological features alongside autism. As these genes are already implicated in well-defined genetic syndromes linked to autism, they are likely to harbor regulatory mutations that could play outsized roles in broad autism-related manifestations. At syndromic ASD genes, we aimed to assess whether high-scoring “outlier” variants, which represent variants with extreme regulatory effects, are more frequent in cases than controls. We ranked the 1,855 mutations by their predicted FLARE-fb scores, and evaluated the proportion of cases according to the ranked scores (Figure 5B).

We found that FLARE-fb successfully prioritized *de novo* non-coding outlier mutations that were preferentially enriched in ASD cases (Supplementary Table 14). For example, 14 of the 16 highest-scoring mutations by FLARE-fb were observed in ASD cases. In contrast, outlier mutations prioritized using other metrics, such as ChromBPNet scores for individual cell types (e.g. fetal early radial glia), FLARE-heart, CADD^89^, or PhyloP^79^, showed minimal to null enrichment for being from ASD cases versus controls under the same analysis framework (Figure 5C). Overall, across the 1,855 mutations, FLARE-fb had significant differences in mean scores (*t*-test *P* = 6.8×10^-3^) but no significant differences in median scores (Mann-Whitney U test *P* > 0.05), indicating that the discriminatory signal was found in the outlier mutations. In contrast, no other metric had significant differences in median or mean scores (nominal *P* > 0.05) (Supplementary Figure 16). These findings emphasize the context-specific strength of FLARE in prioritizing rare non-coding variants contributing to autism risk, which would be missed by using more generalized approaches.

One interesting variant prioritized by the FLARE-fb model was a *de novo* mutation chr14:27,886,425:C>T located in an ENCODE cCRE^5^ in a highly conserved region across multiple vertebrate species. The closest gene was *FOXG1*, located 878 kb downstream, where loss-of-function mutations cause a clinical syndrome characterized by intellectual disability, developmental delays, and microcephaly^90^. This non-coding mutation decreased predicted accessibility across a variety of cell types in the fetal brain, including neurons and glial cells. Using the DeepLIFT method^46^, we interpreted the models and derived predictive contribution scores for each base in the sequence. Our model recognized an RFX motif that becomes weaker at the alternate T allele, and it has been shown that disruption of RFX family transcription factors causes autism and other neurological disorders^91^. Thus, FLARE can pinpoint a non-coding mutation that perturbs binding of an important TF in a constrained region distally located to a known clinical gene (Figure 5D).

Our model also detected three intronic and two upstream de novo mutations near *CNTNAP2*, which would be missed by alternative approaches (Figure 5E). *CNTNAP2* is the largest gene in the human genome, and mutations in *CNTNAP2* have been linked to ASD, intellectual disability, ADHD, schizophrenia, and epilepsy^92^. Using DeepLIFT, the highest-scoring mutation (chr7:148230432:G>T) was predicted to almost completely disrupt NFI binding in a *CNTNAP2* intron (Figure 5F). *NFI* family transcription factors are critical in fetal development^93^ due to their roles in regulating the differentiation of neural progenitor cells^94^, and haploinsufficiency of TFs belonging to this family, such as *NFIB,* is associated with intellectual disability and brain malformations^95^. Overall, FLARE predictions suggest that *de novo* regulatory mutations of *CNTNAP2* are potentially rare causes of ASD in families.

## Discussion

Our study presents a large-scale resource of predicted chromatin accessibility disruptions across adult and fetal brain and heart tissues. By leveraging ChromBPNet, we generated two billion effect predictions for 15 million common and rare variants in 132 cellular contexts, enabling a systematic exploration of how non-coding variation influences gene regulation. These data reveal putative causal variants underlying brain- and heart-related diseases and highlight evolutionary forces that have shaped the cellular landscapes observed in humans. Given their significance, we have made our two billion predictions from 132 ChromBPNet models publicly available for researchers to utilize in their own research.

Traditionally, population-based QTL analyses have linked genetic variants to regulatory effects, but they are often limited to studying common, strong-effect variants^26^. Single-cell QTL studies offer valuable population-level insights yet remain constrained by sample size, power, linkage disequilibrium, and the range of observed cell types^96–98^. In contrast, deep learning models like ChromBPNet, trained on only a handful of high-quality samples, enable genome-wide predictions of variant effects. These approaches inform one another: QTL analyses can guide the interpretation of computational predictions, while models such as ChromBPNet help refine QTL hypotheses. By integrating ChromBPNet with GWAS signals, we identified genetic-driven mechanisms missed by population-level analyses. In Alzheimer’s disease, for instance, we uncovered putative causal variants at GWAS loci and convergent microglia-specific effects genome-wide, demonstrating the power of leveraging context-specific predictions for understanding regulatory mechanisms and disease.

Beyond population insights, our approach also illuminates evolutionary constraints that shape gene regulation. By contrasting ultra-rare and common variants, we found that disruptions in fetal neuronal regulatory elements appear to be under particularly strong purifying selection (concordant with other findings^92–95^). Thus, regulatory networks in fetal neurons exhibit heightened sensitivity to damaging variants^99^, potentially driven by the non-renewing nature of neurons amplifying the lifetime consequences of regulatory disruptions^100,101^. Our findings align with Medawar’s theory of aging, whereby selective pressures wane post-reproduction, potentially allowing greater tolerance of later-acting variants^102,103^ (Figure 3I). Importantly, this indicates that genetic contributors to neurodevelopmental disorders are predominantly rare variants, underscoring the need to map and prioritize damaging rare or *de novo* mutations to better understand their etiology and risk. Common variants, by contrast, are expected to have small individual effects that evade selection, resulting in a modest overall contribution. This is consistent with findings from population cohorts, which attribute only 7–10% of the variance in rare neurodevelopmental disorder risk to common variation^82,104^. A key unanswered question is whether rare developmental disorders in other organ systems, such as the heart, exhibit a greater contribution from common variants, given the relatively weaker selective pressures compared to the fetal brain.

We further introduce FLARE to prioritize impactful rare variants. FLARE can be trained on any context of interest, integrating deep learning predictions with evolutionary constraint. Using a FLARE model of the fetal brain, we enriched for a set of de novo mutations almost exclusively found in autism cases relative to their sibling controls. Distal regulatory mutations have been historically understudied in autism^85–87,105^, and FLARE prioritized mutations with extreme regulatory consequences near autism-associated genes such as *CNTNAP2.* We expect that extending these models to somatic genomes from cancer could help distinguish mutations directly altering accessibility from those arising secondary to chromatin changes^106,107^. FLARE highlights the importance of cellular and developmental context for interpreting rare non-coding variants and establishes a ubiquitous approach for prioritizing and integrating rare variants in risk prediction^86,108–110^.

Our findings underscore the utility of deep learning sequence models for human genetics research in the non-coding genome. Training and deploying these models on single-cell analyses across environments or human development (e.g. dGTEx or HCA)^17,18,107,111^, would illuminate the temporal dynamics of regulatory variants, but also raises the importance of computational efficiency with increasingly large datasets. Subsequently validating these predictions through cost-intensive perturbation experiments, such as being undertaken by the IGVF Consortium^112^, will be essential for confirming causal effects. Furthermore, although we concentrated on variants in accessible chromatin, future efforts could explore those predicted to induce accessibility in closed regions, potentially uncovering pioneer factor binding^113^ where integrating multi-omic data (e.g., expression, splicing, methylation) and including additional epigenomic features may improve variant interpretation efforts^29,35,42,114–118^. Overall, by systematically mining variants and examining their regulatory consequences, we highlight critical roles in evolution, gene regulation, and health.

## Methods

### Curation of single-cell ATAC-seq datasets

We used scATAC-seq data from 5 studies. Our groups have previously published scATAC-seq data on the fetal brain^44^, fetal heart^45^, adult brain^50^, and adult heart, which were incorporated into this study. The adult heart data and cell-type annotations are currently unpublished as part of the latest ENCODE release (see: https://www.encodeproject.org/search/?searchTerm=human+left+ventricle+snATAC&type=Experiment&assay_title=snATAC-seq&limit=100&lab.title=Michael+Snyder%2C+Stanford). The adult heart fragments were generated from ENCODE snATAC-seq from left ventricles in 64 adult human hearts. From each of these samples, fragments were separated by cell-type and then re-grouped into cell-type pseudobulk files. In the suffix of the cell-type annotations, “H”, “I”, “NI”, and “In” refer to samples from the left ventricle from healthy donors or those with ischemic, non-ischemic, or inflammatory cardiomyopathy, respectively.

We additionally downloaded fetal brain and fetal heart data from a chromatin accessibility atlas that were published within the same study^15^ (Domcke et al.), which also mitigates publication-specific batch effects that would exist if contrasting cell types from the brain and heart in downstream analyses. This fetal brain and fetal heart data was downloaded from the following link: <u>descartes.brotmanbaty.org</u>. Cell-type annotations were taken from the published labels in the original papers.

### Processing of single-cell ATAC-seq data

For each single cell cluster from each dataset, we first created pseudobulk fragment files by pooling all of the fragments that belonged to each cluster based on barcode to cell type maps. Then, we converted each fragment file to a tagalign file by splitting each fragment in half to obtain one read per strand and used these as input to the ENCODE ATAC-seq pipeline v1.10.0 (available at https://github.com/ENCODE-DCC/atac-seq-pipeline), which generated peak calls using MACS2^119^ (available at https://github.com/macs3-project/MACS). For downstream analyses, we used the overlap peak set, which only includes peaks called from the original sample that overlap with peak calls from both of the pseudo-replicates created by the pipeline by randomly allocating the reads from the original sample into each synthetic replicate.

### Training ChromBPNet models

To model the cell type–specific chromatin accessibility in each adult and fetal brain and heart cell type, we trained ChromBPNet models^43^ on cell type–resolved pseudobulk ATAC-seq profiles (ChromBPNet is available at https://github.com/kundajelab/ChromBPNet). For each sample, we used as input the peak and tagalign files generated by the ENCODE ATAC-seq pipeline, along with the pre-trained K562 bias model provided in the ChromBPNet repository (https://storage.googleapis.com/ChromBPNet_data/input_files/bias_models/ATAC/ENCSR868F <u>GK_bias_fold_0.h5</u>).

We used a 5-fold cross-validation scheme to train the ChromBPNet models, ensuring that each chromosome appeared in the test set of at least one cross-validation fold. The table below outlines the chromosomes included in the train, validation, and test splits for each fold.

| Fold | Train Chromosomes | Validation Chromosomes | Test Chromosomes |
| --- | --- | --- | --- |
| Fold 0 | chr2, chr4, chr5, chr7, chr9, chr10, chr11, chr12, chr13, chr14, chr15, chr16, chr17, chr18, chr19, chr21, chr22, chrX, chrY | chr8, chr20 | chr1, chr3, chr6 |
| Fold 1 | chr1, chr3, chr4, chr5, chr6, chr7, chr10, chr11, chr13, chr14, chr15, chr18, chr19, chr20, chr21, chr22, chrX, chrY | chr12, chr17 | chr2, chr8, chr9, chr16 |
| Fold 2 | chr1, chr2, chr3, chr5, chr6, chr8, chr9, chr10, chr13, chr14, chr16, chr17, chr18, chr19, chr20, chr21, chrX | chr7, chr22 | chr4, chr11, chr12, chr15, chrY |
| Fold 3 | chr1, chr2, chr3, chr4, chr7, chr8, chr9, chr11, chr12, chr13, chr15, chr16, chr17, chr19, chrX, chrY | chr6, chr21 | chr5, chr10, chr14, chr18, chr20, chr22 |
| Fold 4 | chr1, chr2, chr3, chr4, chr5, chr6, chr8, chr9, chr11, chr12, chr14, chr15, chr16, chr20, chr22, chrY | chr10, chr18 | chr7, chr13, chr17, chr19, chr21, chrX |

One of the benefits of using ChromBPNet to model chromatin accessibility is that its bias-correction procedure can automatically regress out the contributions to the base-pair level predictions coming from the sequence preferences of certain enzymes used in assays, such as tn5 for ATAC-seq in the samples used in this project. Upon completion of training, any model that continued to respond to the tn5 motif’s consensus sequences after unplugging the bias model, indicating an unsuccessful bias-correction procedure, was retrained until the model showed a limited response to the tn5 motif’s consensus sequences embedded in random genomic backgrounds. Specifically, this retraining was performed if the maximum prediction from the profile head for any base-pair from the examples containing the tn5 motif exceeded 0.002.

### Removal of cell types with poor ChromBPNet model performance

After training, we removed outlier models where predicted read counts in peaks deviated from observed read counts in peaks to a much greater extent than other models. We set an outlier threshold of greater than 5 standard deviations from the mean model performance (calculated using spearman’s correlation coefficient) across all models. We removed two contexts with models exceeding the outlier threshold, resulting in 132 models representing different cellular contexts across 4 developmental organ contexts (fetal brain, fetal heart, adult heart, adult brain) from 5 distinct studies.

### Curation of variants from the 1000 Genomes Project

The New York Genome Center, funded by NHGRI, sequenced 3,202 individuals from the 1000 Genomes Project at 30× coverage, with variant data aligned to the GRCh38 reference genome^49^. We downloaded the data from http://ftp.1000genomes.ebi.ac.uk/vol1/ftp/data_collections/1000G_2504_high_coverage/working/20201028_3202_raw_GT_with_annot/20201028_CCDG_14151_B01_GRM_WGS_2020-08-05_chr${chrNum}.recalibrated_variants.vcf.gz and limited our analysis to autosomal chromosomes 1 to 22. Variants with missing or unspecified alleles (“*”) were excluded, and VCF genotypes were converted to PLINK format for downstream analysis^120^. We further created PLINK files for a European ancestry subset of 633 individuals and retained only variants with a minor allele count of at least one. Minor allele frequencies (MAF) were computed for the full cohort of 3,202 individuals, encompassing diverse global populations, and separately for the European subset. Indels were excluded from further analysis, resulting in a curated dataset of 6,349,771 common SNPs (MAF > 5% in both global and European populations) and 8,757,029 rare SNPs (MAF < 1%), totaling 15,106,800 curated common and rare SNPs genome-wide.

### Annotation of non-coding variants

We annotated selective constraint for variants using PhyloP scores^79^ by downloading the “hg38.phyloP100way.bedGraph” file from the UCSC Genome Browser^121^. Variants were annotated using the “bedtools intersect” command, excluding any variants not present in the UCSC PhyloP file from PhyloP-based analyses. Additional annotations, including VEP consequences^122^, CADD-related annotations^89^, and gnomAD minor allele frequencies^123^, were performed using a Snakemake pipeline available at https://github.com/e-271/watershed_dataprep/tree/main. VEP annotations were generated using VEP version 110, while gnomAD minor allele frequencies were annotated using gnomAD v4.0.0. Variants were annotated with the gene constraint values for the nearest TSS, leveraging *s*_het_ calculations based on prior work^124^. We focused on non-coding variants and excluded variants with VEP consequences indicative of coding or protein-altering changes, including terms such as “coding_sequence_variant,” “missense_variant,” “synonymous_variant,” “stop_gained,” “stop_lost,” “frameshift_variant,” “inframe_insertion,” “inframe_deletion,” “splice_donor_variant,” “splice_acceptor_variant,” “start_lost,” “non_coding_transcript_exon_variant,” “mature_miRNA_variant,” “start_retained_variant,” “stop_retained_variant,” “incomplete_terminal_codon_variant,” and “protein_altering_variant.” We additionally filtered all splicing-associated variants from later analyses.

TSS coordinates were obtained using the start position of protein coding genes on the positive strand and the end position of protein coding genes on the negative strand from the GENCODE v47 basic annotation GTF file for hg38, available at: https://ftp.ebi.ac.uk/pub/databases/gencode/Gencode_human/release_47/gencode.v47.basic.annot <u>ation.gtf.gz</u>

### Predicting variant effects using ChromBPNet models

To score each variant in this study, we used the ChromBPNet models for each cell type to predict the base-resolution scATAC-seq pseudobulk coverage profiles for the 1 kb genomic sequence centered at each variant and containing the reference and alternate allele. We then estimated the variant’s effect size using two measures: (1) the log2 fold change in total predicted coverage (total counts) within each 1 kb window for the alternate versus reference allele, and (2) the Jensen–Shannon distance (JSD) between the base-resolution predicted probability profiles for the reference and alternate allele (capturing changes in profile shape).

We assessed statistical significance for these scores using empirical null distributions constructed by shuffling the 2114 bp sequence around each variant multiple times while preserving dinucleotide frequency. Next, each shuffled sequence was duplicated, and the variant’s reference or alternate allele was inserted at the center, resulting in a total of one million null variants for each set of observed variants scored. Each null variant was scored with the same procedure as the observed variants. For each observed variant, we then computed the proportion of null variants with an equally high or higher (more extreme) score to derive empirical *P*-values for both the log2 fold change and JSD scores. For all analyses in this study, we use the log2 fold change score, which we refer to as the ChromBPNet score. The code repository for scoring variants is at https://github.com/kundajelab/variant-scorer.

### Inferring predictive motif instances using ChromBPNet

In order to identify the cell-type-specific motif instances that are predictive of accessibility, we first ran the DeepLIFT^46^ algorithm on the counts prediction for all peaks in each brain cell type using the ChromBPNet model for each cross-validation fold to generate contribution scores for each base-pair. The DeepLIFT contribution scores were generated using the DeepSHAP^47^ implementation of the DeepLIFT algorithm (available at https://github.com/kundajelab/shap/tree/master) with 20 dinucleotide shuffled reference sequences for each peak sequence. Since many peaks have multiple instances with identical genomic coordinates but different summit positions called by the peak-caller, we used the peak instance with the highest -log10(q-value) from MACS2 among peaks with identical coordinates. Then, for each cell type, we computed the mean contribution scores across folds and used them as input to the TF-MoDISco^48^ tool (https://github.com/jmschrei/tfmodisco-lite), which identified a unique set of sequence patterns corresponding to both known and novel transcription factor motifs learned by the models trained in that cell type. We ran TF-MoDISco using the contribution scores for the 500 bp window around each peak summit and allowed up to 1 million seqlets per metacluster for each run.

Since many of these motif patterns are shared across cell types, we used the MotifCompendium tool (which will soon be available at https://github.com/kundajelab/MotifCompendium), which uses an efficient, GPU-accelerated method to calculate pairwise similarity and cluster a large number of motif patterns, to create a non-redundant set of clustered patterns discovered across all brain cell types. A similarity score threshold of 0.96 was used for clustering the 2488 individual motif patterns into 999 non-redundant patterns. Next, we used the Fi-NeMo tool (available at https://github.com/austintwang/finemo_gpu) to identify the specific genomic instances of each of these clustered motif patterns within the ATAC-seq peaks of each brain cell type using the DeepLIFT contribution scores generated for those peaks. We ran Fi-NeMo separately using both the unique TF-MoDISco contribution weight matrices (CWMs) originally identified for each pattern in each individual cell type and the mean CWMs representing the motif clusters across all brain cell types. A lambda regularization parameter of 0.8 was used for Fi-NeMo. For downstream analyses, the motif instances attributed to a given motif cluster in a specific cell type were the Fi-NeMo instances mapped to all original TF-MoDISco patterns that belonged to that motif cluster in that cell type. For any motif cluster that did not have any original TF-MoDISco pattern originating from that cell type, the Fi-NeMo instances mapped to the mean CWM for the motif cluster were used. This ensured that the exact motif pattern learned in each cell type was used to determine the motif instances that mapped to that motif cluster in each cell type whenever the original pattern was available, while allowing for matches to motifs that were not originally detected by TF-MoDISco in each cell type. Furthermore, this allowed us to identify any cases of aberrant motif instance-calling, as we filtered out those motif clusters that had more than 5 times the number of instances mapped to the mean CWM of the motif cluster than were mapped to the original patterns that belonged to that motif cluster in any cell type.

Finally, for downstream analyses, we considered the clustered motif patterns that were supported by at least 500 total seqlets from TF-MoDISco, annotated the consensus motif of these clusters based on similarity to known transcription factor motifs from JASPAR^125^, CIS-BP^126^, and HOCOMOCO^127^, obtained from the curated set^128^ at https://resources.altius.org/~jvierstra/projects/motif-clustering-v2.1beta/, and further filtered out those patterns that were either simple CG or AT repeats or could not be reliably matched to individual motifs or composites of known motifs. Clustered patterns that matched to a composite of known motifs were annotated as pairs of motif labels separated by “:”, while multiple clustered patterns that matched to the same known motif or motif composite were distinguished using a “_n” suffix in the annotation, where “n” is the n-th clustered pattern that matched to the same known motif or motif composite. Altogether, these yielded a final set of 141 annotated clustered motif patterns that were used in downstream analyses (Supplementary File 1).

### Analysis of genomic context and cell-type-specificity

To investigate the relationship between genomic context and cell-type specificity, we utilized single-cell ATAC-seq data from Domcke et al. (2021)^15^. This dataset exclusively encompasses fetal developmental samples across a limited number of cell types, avoiding potential confounders such as overlapping cell types across datasets or variability due to developmental stage (adult vs. fetal). By concentrating on a single developmental time point, we ensured that the analysis highlights differences in cell-type specificity among variants. Compared to other datasets, such as Ameen et al.^45^ and Trevino et al.^44^, which focus specifically on the fetal heart (20 cell types) and fetal brain (22 cell types), Domcke et al. offers broader but coarser annotations, spanning 21 cell types across both fetal brain and heart.

For each variant, regulatory magnitude was defined as the highest ChromBPNet score observed across fetal brain and heart cell types. Variant cell-type specificity was quantified as the number of cellular contexts in which the variant was accessible and had a ChromBPNet empirical *P*-value < 0.01 within the same cell type. Analyses were restricted to variants accessible in at least one cell type. Variants were further categorized into four cell-type specificity groups based on their predicted impact: “null” (impacting 0 cell types), “specific” (impacting only 1 cell type), “shared” (impacting >80% of cell types), and “multiple” (impacting more than 1 but fewer than 80% of cell types).

We conducted two statistical analyses to evaluate relationships between variant characteristics and genomic context. First, we examined the association between cell-type specificity group and distance to the nearest transcription start site (TSS) using a linear model, with “null” as the reference category in the factor variable setup. Second, we assessed the relationship between regulatory magnitude and TSS distance. In both analyses, TSS distance was modeled as log⁡_10_(distance to nearest TSS+1), with the constraint value of the nearest gene included as an additional covariate.

In the results presented in the main text, “accessible but non-functional variants” represent a set of variants that were accessible in at least 1 cell type but not predicted to affect accessibility in any cell type. This is equivalent to the “null” cell-type specificity group.

### Analysis of large-effect cell-type-specific versus shared variants

We identified variants with regulatory magnitudes in the top 1% across all analyzed variants and classified them as either shared or specific based on the definitions used in the prior analysis. Using *bedtools intersect*, we overlapped these variants with experimentally verified human promoter regions obtained from the Eukaryotic Promoter Database^59^. The proportion of variants falling within promoter regions was calculated separately for shared and specific variants.

To further annotate the variants, we utilized the motifbreakR tool^80^ with the information content method, applying a threshold of 10^-4^ and using the HOCOMOCOv11-core-A, HOCOMOCOv11-core-B, and HOCOMOCOv11-core-C motif databases with filterp setting equal to TRUE. Variants were not filtered further, allowing a single variant to have multiple entries corresponding to potential disruptions of different transcription factors. We calculated the proportion of variants predicted to disrupt motifs and focused subsequent analyses on motifs disrupted in at least 5% of specific or shared variants. Enrichment was assessed using the absolute value of the log2 fold-change in the proportion of shared versus specific variants disrupting each transcription factor motif, and we filtered by limiting to motifs with values greater than 2. For visualization, the top 10 motifs most relevant to shared and specific variants were selected based on their proportions.

### Acquisition and processing of GTEx QTLs

We obtained publicly available cis-eQTL fine-mapping data from the Genotype-Tissue Expression (GTEx) Project (supported by the NIH Common Fund, NCI, NHGRI, NHLBI, NIDA, NIMH, and NINDS). All data used in this study were downloaded from the GTEx Portal (https://gtexportal.org/home/datasets) on September 18, 2024. Specifically, we retrieved the “Fine-Mapping cis-eQTL Data” results generated using the DAP-G algorithm^129^.

From all GTEx brain tissue datasets, we downloaded all fine-mapped eQTL results and merged the files. For each variant, we calculated the maximum posterior inclusion probability (PIP), and identified variants with high posterior probability of being causal (PIP > 0.9) as well as those with low posterior probability (PIP < 0.01) but that remained within the tested fine-mapped region. The former group served as putatively causal variants. We repeated the same using all heart and artery tissue eQTL datasets in GTEx.

### Analysis of GTEx QTLs

To evaluate whether fine-mapped variants with PIP > 0.9 had greater predicted impact on chromatin accessibility, we integrated cell-type–specific chromatin accessibility predictions generated by ChromBPNet. We assessed each variant’s predicted chromatin accessibility effect in multiple cell types, including various adult and fetal brain cell populations, as well as adult and fetal heart cell populations. We then tested whether the fine-mapped variants (PIP > 0.9) were associated with higher ChromBPNet scores compared to low-PIP variants (PIP < 0.01). We limited the analysis to accessible variants only. Additionally, we examined whether fine-mapped variants influenced a greater number of distinct cell types, evaluating the association between the number of affected cell types and the variant’s putative causality (PIP classification) genome-wide.

For statistical analyses, we applied a linear model with the fine-mapping PIP classification (high vs. low) as a categorical predictor. Continuous covariates included the log10-transformed distance to the nearest transcription start site (TSS) plus one [log_10_(distance to nearest TSS+1)] and gene-level constraint (*s*_het_) for the nearest gene. These models allowed us to control for genomic context and gene-level constraints when assessing the relationship between inferred eQTL causality and accessibility predictions. We visualized the *P*-values for the fine-mapped categorical predictor from these models within the manuscript figures.

### Comparison of GTEx bulk eQTLs to genome-wide regulatory variants

To assess cell-type specificity underlying brain eQTLs relative to other regulatory variants, we considered all genome-wide variants predicted to have effects in adult excitatory neurons or adult microglia. Variants were included if they had a ChromBPNet *P*-value < 0.01 in at least one of these cell types. From this set, we identified variants that showed regulatory effects in both adult neurons and adult microglia.

We then performed a logistic regression to test for an association between membership in this neuron–microglia-shared variant set and membership in the fine-mapped eQTL set (PIP > 0.9). To control for potential confounding factors, we included multiple covariates in the logistic model: ChromBPNet scores in neurons and microglia, non-Finnish European minor allele frequency (MAF) from gnomAD, distance to the nearest TSS, and the constraint of the nearest gene. This approach ensured that any observed association between shared regulatory variants and fine-mapped eQTLs was not driven by differences in allelic frequency, genomic context, or gene constraint.

### Comparison of neuron-specific eQTLs to non-neuron-specific eQTLs

To explore cell-type specificity further, we identified neuron-specific eQTL variants by leveraging interaction eQTL (ieQTL) data from the GTEx portal (brain-cerebellum neuron ieQTL analysis, originally identified using eigenMT)^60,130^. Variants with neuron ieQTL interaction *P*-values below 0.05 were considered neuron-specific eQTLs. We then compared the distribution of ChromBPNet scores for adult microglia and adult excitatory neurons at neuron-specific eQTLs versus non–neuron-specific variants within the fine-mapped bulk brain eQTL set using a t-test. This comparison allowed us to determine whether neuron-specific regulatory variants exhibit distinct chromatin accessibility patterns compared to other regulatory variants.

### Cell-type-relevance of open chromatin regions to GWAS

We used stratified linkage disequilibrium score regression (LDSC)^61^ to assess the enrichment of GWAS heritability in open chromatin regions across 132 cellular contexts. Peak files, initially aligned to the hg38 reference genome, were converted to hg19 using liftOver. These converted peak files were then transformed into LDSC annotation files using the 1000 Genomes Project reference dataset, following the make_annot.py and ldsc.py pipelines. Thin annotations were created with a 1 cM LD window specified using --ld-wind-cm 1, and annotation files were streamlined by including only HapMap SNPs through the --print-snps option to reduce file size. The resulting l2.ldscore.gz files were subsequently used in the LD score regression analyses.

To facilitate these analyses, we constructed ldcts files and generated a joint annotation that represented the union of all peaks across the 132 cell types. GWAS summary statistics were obtained from the GWAS Catalog or directly from authors’ publications, with genomic positions converted to hg38 and alleles standardized to ref/alt format based on the 1000 Genomes Project. The harmonized summary statistics were formatted for LDSC using munge_sumstats.py, with sample sizes provided via --N, or, where relevant, --N-cas and --N-con. We used the HapMap3 SNP list (w_hm3.snplist) with the --merge-alleles option, processed in chunks of 500,000 SNPs (--chunksize 500000), and ensured proper allele alignment using the --signed-sumstats, --a1, and --a2 options.

LDSC analyses were parallelized for each of the five datasets, specifying the ldcts file through --ref-ld-chr-cts in a --h2-cts analysis. Both dataset-specific union peak annotations and baseline LD score annotation files for each cell type were provided via the --ref-ld-chr command. For each GWAS, false discovery rates (FDR) were calculated independently across the 132 cellular contexts, with significance defined as FDR < 0.1.

### UK Biobank GWAS fine-mapping analysis

We analyzed GWAS fine-mapping data from the UK Biobank, downloaded from https://www.finucanelab.org/data. Briefly, this dataset uses association analyses for 94 traits involving 361,194 White British individuals from the UK Biobank, where fine-mapping was performed using an in-sample linkage disequilibrium (LD) reference panel for genome-wide significant variants (P < 5×10^−8^) within a 3 Mb window. Overlapping regions were merged, and analyses were conducted using FINEMAP and SUSIE, with a maximum of 10 causal variants per locus. Variants within the major histocompatibility complex (MHC) region were excluded from the analysis^65^.

Given the smaller number of fine-mapped variants identified for certain GWAS phenotypes compared to GTEx eQTL analyses, we applied a posterior inclusion probability (PIP) threshold of >0.2 to retain more putative causal variants for downstream analyses. Subsequently, for each cell type, we replicated the same analysis framework as was used for eQTLs, applying it to GWAS fine-mapped variants.

### Fine-mapping of GWAS meta-analyses

GWAS summary statistics were obtained from the GWAS Catalog or directly from links provided in the authors’ publications. Genomic positions were converted to the hg38 reference genome, and alleles were standardized to ref/alt format based on the 1000 Genomes Project genotype file. For each GWAS locus, we identified a lead variant, ensuring a minimum distance of 1 Mb between lead variants. Variants within the major histocompatibility complex (MHC) region were excluded from the fine-mapping analyses. Linkage disequilibrium (LD) was calculated using the 1000 Genomes Project European genotype data as the reference panel, employing the PLINK --r square command.

Some issues with missing (NA) values in the LD matrix were resolved systematically. NA values in LD calculations can arise from two scenarios: (1) SNPs with no variation (e.g., all genotypes are the same) result in NA values across all SNPs; (2) SNP pairs where at least one SNP has missing genotype calls, leading to variation at one SNP being confined to the individuals with missing data at the other SNP. To address this, we flagged all SNPs with no variation (case 1), identified problematic SNP-SNP combinations (case 2), and randomly removed one SNP from each flagged pair while retaining lead SNPs in downstream analysis.

Fine-mapping was performed using SuSiE, specifying the sample size of the GWAS and testing two models: L=1 (allowing for a single causal signal) and L=10 (allowing for up to ten causal signals). The L=1 analysis was included to mitigate potential mismatches between the LD reference panel and GWAS meta-analysis summary statistics. Results from both analyses were compiled, and the maximum posterior inclusion probability (PIP) (between the two analyses) for each variant was used in downstream enrichment analyses. The same linear model as the UK Biobank GWAS and GTEx eQTLs was used for evaluating statistical significance.

### GWAS variant analysis and gene set enrichment analysis in Alzheimer’s disease

Microglia-specific GWAS variants were defined as those with a posterior inclusion probability (PIP) > 0.005, a predicted effect in microglia (ChromBPNet empirical *P*-value < 0.05), and no predicted effects in other adult brain cell types (ChromBPNet empirical *P*-value > 0.05 in all other contexts). Variants located on chromosome 6 were excluded to avoid confounding effects from the major histocompatibility complex (MHC) region. For each selected variant, we identified the two closest transcription start sites (TSS) and their corresponding genes, yielding a list of candidate genes. This gene list was used as input for gene set enrichment analysis (GSEA) with Enrichr, focusing on the “GO_Biological_Process_2021” and “GO_Molecular_Function_2021” databases. To perform a baseline, cell-type-agnostic GSEA, we repeated the procedure without considering ChromBPNet predictions. In this case, we included all variants with PIP > 0.005 and their two closest genes based on TSS distance. We restricted both analyses to significant gene sets containing at least two genes, excluding sets with only a single gene. For the *PICALM* analysis, we used the L=10 results from the Alzheimer’s GWAS meta-analysis to define the two credible sets and calculate PIP values for the *PICALM* locus.

### Analysis of common versus rare variants

To compare the regulatory effects of accessible rare and common variants across cell types, we employed a linear modeling approach. Variants were first subsetted to those located in accessible chromatin regions (peaks). The dependent variable in the model was the standardized ChromBPNet score, representing the predicted regulatory effect of each variant. A binary variable indicated whether a variant belonged to the rare or common variant set. Additional covariates included the logarithm of the transcription start site (TSS) distance plus one (log⁡10(TSS distance+1)) and the constraint score of the nearest gene to account for genomic context. A separate linear model was run for each cell type to estimate the effect of variant rarity on predicted regulatory magnitude. Results were visualized by sorting the estimated effect sizes (betas) from these models, with 95% confidence intervals included to assess statistical robustness.

To investigate differences in effect sizes in immune contexts, we categorized cellular contexts as immune or non-immune. Immune cell types included lymphocytes, macrophages, mast cells, microglia, and myeloid cells, specifically: “LV.lymphocyte.H,” “LV.lymphocyte.I,” “LV.lymphocyte.In,” “LV.lymphocyte.NI,” “fetal_heart.Myeloid_cells,” “LV.macrophage.H,” “LV.macrophage.I,” “LV.macrophage.In,” “LV.macrophage.NI,” “LV.mast_cell.H,” “LV.mast_cell.I,” “LV.mast_cell.In,” “LV.mast_cell.NI,” and “Microglia.” Non-immune cell types comprised all other contexts. A factor variable distinguishing immune from non-immune cell types was included in a linear model, along with covariates for context (adult or fetal brain and heart) and mean model performance across folds (Spearman correlation within peak regions). This analysis quantified differences in the cell-type-specific betas describing the per-cell-type variation in ChromBPNet scores between common and rare variants, enabling assessment of effects across immune and non-immune contexts.

To address potential confounding due to the set of variants analyzed across cell types, we conducted a robustness analysis focused on excitatory neurons in the fetal brain, identified as the top-ranked cell type. For each additional cell type, we restricted the analysis to variants that were jointly accessible in both the fetal brain excitatory neurons and the cell type of interest. Using this shared set of variants, we reanalyzed ChromBPNet scores to compare the predicted regulatory effects of common versus rare variants. Effect sizes were calculated as the mean difference in predicted effects between common and rare variants for each cell type. To quantify differences between the two cell types, we calculated the difference in effect sizes from the ChromBPNet analyses, along with 95% confidence intervals. By limiting the analysis to a shared set of peaks, we controlled for potential confounding due to differences in variant accessibility across cell types. This approach ensured that any observed differences reflected intrinsic cell-type-specific effects rather than disparities in the variant set analyzed.

We analyzed the proportion of accessible variants for common versus rare variants across cell types using genome-wide variants in a logistic regression model. This complementary approach allowed us to evaluate variant effects at both the regulatory level (via ChromBPNet, the prior analyses) and the accessibility level (via peaks analysis). To assess the consistency between these methods, we quantified the similarity of their findings by correlating the effect sizes estimated from the ChromBPNet models with those from the peaks-based analysis. This comparison provided insight into the alignment of predictions from the two analyses.

To assess differences in cell-type specificity between rare and common variants, we focused on accessible variants associated with at least one cell type. For each variant, we quantified cell-type specificity as the number of cell types in which the variant was accessible and had a ChromBPNet empirical *P*-value < 0.01 in the same cell type. Variants were then classified as rare or common based on their minor allele frequencies, and the average number of cell types impacted by each category was compared using a linear model accounting for *s*_het_ and log_10_(distance to the nearest transcription start site + 1). Analyses were stratified by developmental and organ contexts, including fetal and adult brain and heart, to evaluate how trends in cell-type specificity varied across these contexts.

### Analysis of rare variants with increased population frequencies

We analyzed rare variants from the 1000 Genomes Project (1KG) that exhibited higher minor allele frequencies (MAF > 1%) in different gnomAD populations. This analysis aimed to complement the rare versus common variant comparisons by investigating variants whose frequencies may have increased due to genetic drift, population resampling, or founder effects. Seven gnomAD populations were included in the analysis: “afr” (African), “amr” (Latino/Admixed American), “asj” (Ashkenazi Jewish), “eas” (East Asian), “fin” (Finnish), “nfe” (Non-Finnish European), and “sas” (South Asian), with “asj” and “fin” reflecting founder populations. We evaluated two aspects of regulatory function for these variants: (i) Cell-Type Specificity: The number of cell types in which the variant was accessible and had a ChromBPNet empirical *P*-value < 0.01, and (ii) Regulatory Magnitude: The maximum ChromBPNet score across all cell types. Both metrics were standardized to have a mean of 0 and standard deviation of 1 within each analysis. For each metric, we applied a linear model with the following covariates: a binarized variable indicating whether the variant frequency exceeded 0.01 in the population of interest, *s*_het_, a measure of selective constraint, and log_10_(distance to the nearest transcription start site + 1). The estimated effect sizes for the binarized frequency variable (indicating common or rare frequency) were reported and visualized for each population. This approach allowed us to compare trends in regulatory function across populations, including founder populations where higher frequencies of deleterious variants are observed due to historical bottlenecks, genetic drift, and limited gene flow. By incorporating these population-specific effects, we aimed to capture regulatory trends that distinguish global purifying selection from founder population dynamics.

### Correlating constraint with regulatory effects at rare variants

We correlated constraint, as measured by PhyloP, with ChromBPNet scores using a linear model. We tested each cell type individually. We limited the analysis to accessible variants, and removed SNPs with missing PhyloP values. ChromBPNet scores were standardized to have a mean of 0 and standard deviation of 1 within each cell type. PhyloP was tested as the dependent variable, accounting for *s*_het_ and log_10_(distance to the nearest transcription start site + 1) as additional covariates. The results were compared to the common versus rare analyses by computing the correlation between effect sizes between the two.

### Identification of development-specific excitatory neuron variants

We obtained a set of ultra-rare variants (MAF < 0.1%) from the 1000 Genomes Project that occur within accessible chromatin in both fetal and adult excitatory neurons. From these variants, we identified those with context-specific regulatory effects by applying several filters. Specifically, we required that each variant: (1) had a significant ChromBPNet score (*P* < 0.01) in one developmental context (fetal or adult) but not in the other (P > 0.1), and (2) exhibited a difference in ChromBPNet scores between fetal and adult contexts in the top or bottom 10% of all co-accessible variants. After applying these criteria, we obtained a set of 1,617 development-specific excitatory neuron variants that are accessible across development but have strong, context-dependent regulatory effects.

### Testing for differences in evolutionary constraint of development-specific variants

To assess the evolutionary constraint of these development-specific variants, we first compared adult-specific and fetal-specific variants separately to a background set of co-accessible variants that lacked developmental specificity. We used PhyloP scores as a measure of evolutionary conservation and fit linear models to test differences in conservation between each development-specific set and the non-specific set, accounting for log_10_(distance to the nearest transcription start site + 1) and nearest gene constraint (as quantified by *s*_het_), and performed the same analysis for fetal-specific variants.

Next, we directly compared the adult-specific variants to the fetal-specific variants using a similar linear modeling approach. We limited the analysis to adult-specific or fetal-specific variants, testing for differences in PhyloP scores using a linear model between the two groups while accounting for log_10_(distance to the nearest TSS + 1) and nearest gene constraint (as quantified by *s*_het_).

### Analysis of predicted motif disruptions of development-specific variants

We used motifbreakR to identify TF motifs disrupted by the development-specific variants. For both adult-specific and fetal-specific variant sets, we calculated the proportion of variants predicted to disrupt each TF motif. We then focused on motifs meeting two criteria: (1) a greater than 4-fold difference in the proportion of disrupted motifs between adult and fetal variants, and (2) at least 5% of variants in the more affected group predicted to disrupt that motif. TFs corresponding to these motifs were reported in the main text.

We additionally used ChromBPNet to identify putative motif instances impacting particular cell types. For all development-specific variants, we generated DeepLIFT contribution scores using the ChromBPNet counts prediction for both allele of each variant using the models for all cross-validation folds, computed the mean contribution scores across folds, and applied the Fi-NeMo tool to these contribution scores and the annotated motif patterns identified across all brain cell types to label the motifs overlapping either allele of these variants.

### Analysis of predicted TF binding disruptions of development-specific variants

We applied deltaSVM^131^ trained on SNP-SELEX^81^ experiments to estimate the predicted transcription factor (TF) binding impact of 1,617 variants specific to fetal or adult excitatory neurons, encompassing 94 TFs in total. Prior to enrichment analyses, we normalized all deltaSVM scores on a per-TF basis, ensuring a mean of zero and a standard deviation of one. Next, we focused on variants annotated by the model as likely to occur in TF binding sites, filtering out those for which TF binding was not predicted. We then identified variants that surpassed a stringent z-score threshold (≥ 2) relative to each TF’s average distribution. Using this set of variants, we tested for enrichment distinguishing fetal- and adult-specific variants, which would imply predicted differences in TF binding. We calculated the odds ratio to measure the enrichment of TF binding effects in fetal- and adult-specific variants using logistic regression, where we compared the presence and absence of predicted binding sites for each group. The odds ratio was log2-transformed for interpretability, with positive values indicating enrichment in fetal-specific variants and negative values indicating enrichment in adult-specific variants. We also applied a slightly less stringent threshold (z-score ≥ 1) to evaluate the choice of threshold. Finally, to contextualize these enrichments, we examined TF expression patterns using the BrainSpan web interface for developmental brain data and the GTEx portal for regional brain expression profiles.

### FLARE method details

FLARE (Functional Lasso for Accessible Regulatory Elements) employs a cutting-edge regularized regression model to predict evolutionary conservation from single-cell chromatin accessibility data. This adaptable approach can be applied to any ATAC-seq dataset, enabling the generation of FLARE models tailored to specific developmental and tissue contexts.

For a desired collection of cell types, we construct a comprehensive feature matrix for a set of variants (here, 8,757,029 rare SNPs). The feature matrix includes:

1. *Genomic context*: Log-transformed distance to the nearest transcription start site (TSS), calculated as log_10_(distance to the nearest TSS + 1)).
2. *Gene constraint*: the *s*_het_ constraint score of the gene closest to the TSS.
3. *Chromatin accessibility*: Accessibility scores for each cell type in the dataset.
4. *Predicted accessibility effects*: ChromBPNet-derived regulatory scores for each cell type.
5. *Conditional predicted accessibility effects*: A ChromBPNet-based measure set to “0” if the variant is not accessible in a given cell type.

FLARE uses a linear regression framework with Gaussian errors and L1 loss (lasso regression) to model PhyloP conservation scores from this feature matrix. To ensure biological interpretability, coefficients for all ATAC-seq-derived features are constrained to be non-negative. Models are trained using a leave-one-chromosome-out approach, resulting in 22 independent models, one for each chromosome. The optimal regularization parameter is determined via 4-fold cross-validation, and model performance is evaluated on held-out chromosomes by comparing predictions to observed PhyloP values using Pearson correlation. Variants with missing PhyloP annotations are excluded from training and testing.

FLARE was applied across five primary tissue contexts: (i) fetal brain, (ii) adult brain, (iii) fetal and adult heart, (iv) combined fetal and adult brain, and (v) combined brain and heart. To benchmark performance, FLARE was compared to two alternative models: a baseline model excluding ATAC-seq features and a peak-only fetal brain model. Additionally, FLARE was evaluated on curated sets of common variants from the 1000 Genomes Project and de novo mutations from the Simons Simplex Collection.

### FLARE application to autism

*De novo* mutations from 1,902 children with Autism Spectrum Disorder (ASD) and their unaffected siblings were obtained from the Simons Simplex Collection, as previously described in An et al. 2018^86^. Variants annotated with coding or splice consequences (based on Gencode v27 annotations) were excluded from the analysis, as were mutations observed in gnomAD^86^, located on non-standard chromosomes, or within low-complexity repeat regions identified by RepeatMasker^132^. Additionally, non-singleton de novo mutations—those found in both affected and unaffected siblings or in multiple SSC families—were removed. These processed and published data were used as the starting point for our analysis. We further removed variants from the published list that had potential coding or splicing consequences within our annotation step.

We focused on *de novo* non-coding mutations located near genes associated with syndromic forms of autism. Syndromic ASD genes were defined as those with at least 10 reports in the SFARI database and annotated as syndromic. Variants that were retained in analysis include those assigned to these genes based on proximity to the variant’s closest transcription start site (TSS) or gene annotations described in Trevino et al^44^.

To identify regulatory mutations with outsized contributions to ASD, we ranked the mutations by their values for a score of interest (e.g. FLARE fetal brain). In this way, high-scoring “outlier” variants were prioritized at the top of the list, representing those with extreme predicted effects. For each rank threshold, we evaluated the proportion of variants from ASD cases using a binomial test, with the overall proportion of case variants in the dataset as the background expectation. Confidence intervals and *P*-values were calculated for each rank threshold to assess enrichment of case variants. We compared performances of several metrics, including fetal early radial glia ChromBPNet scores, FLARE predictions, CADD scores, and PhyloP conservation scores. For each metric, mutations were ranked, and the enrichment of case variants among the top-ranked mutations was assessed using the same binomial testing framework. We additionally tested for differences in FLARE-fb scores between case and control mutations across all 1,855 variants using a *t*-test for mean differences and a Mann-Whitney U test for median differences. Variants with missing values for any metric were excluded from analysis.

## Acknowledgements

This study was supported by NIH 5U01HG012069-03 (NHGRI IGVF – Kundaje Grant) and NIH 5R01MH125244-04 (NIMH). D.P.M. is supported by NSF GRFP (DGE-2146755). The Genotype-Tissue Expression (GTEx) Project was supported by the Common Fund of the Office of the Director of the National Institutes of Health, and by NCI, NHGRI, NHLBI, NIDA, NIMH, and NINDS. The data used for the analyses described in this manuscript were obtained from the GTEx Portal on 9/18/24. BioRender was used to help make figures.

## Author Contributions

A.R.M., S.K., A.K., and S.B.M. designed the study. A.R.M. and S.K. led and oversaw all analyses. S.D. processed fetal heart data, and S.D. and A.W. processed the adult heart data. A.W. developed Fi-NeMo, a method for calling motif instances in the genome using DeepLIFT contribution scores and motif patterns identified by TF-MoDISco. S.D. and C.M.Y. co-developed the method for clustering and automatically annotating motif patterns from TF-MoDISco. S.J. contributed to the manual annotation of motif patterns from TF-MoDISco. E.M.P. provided computational support, including common versus rare analyses, SNP-SELEX analysis, and motifbreakR. D.P.M. conducted the final SNP-SELEX analysis. E.R. and D.N. assisted with VEP, CADD, and gnomAD variant annotations. Y.S. contributed evolutionary annotations of variants. Y.X. helped inform fine-mapping enrichment analyses. A.R.M. and S.K. wrote the original manuscript draft. All authors edited and reviewed the manuscript.

## Data and Code Availability

We have deposited the custom code for this manuscript at https://github.com/kundajelab/neuro-variants. This makes use of the generalized code for:

- Training new models and generating DeepLIFT scores from: https://github.com/kundajelab/ChromBPNet
- Scoring new variants of interest from: https://github.com/kundajelab/variant-scorer
- Running TF-MoDISco from: https://github.com/jmschrei/tfmodisco-lite
- Clustering motif patterns from: https://github.com/kundajelab/MotifCompendium
- Calling motif instances from: https://github.com/austintwang/finemo_gpu
- Training FLARE models for variant scoring: https://github.com/drewmard/FLARE

We have made available our peak calls, ChromBPNet models, ChromBPNet variant scores, and FLARE variant scores at https://www.synapse.org/Synapse:syn64693551/files/. Additional generalized code for GWAS harmonization, variant curation, and fine-mapping can be found at https://github.com/drewmard/VariantPrioritization or https://github.com/drewmard/HarmonizeGWAS.

## References

1. Claussnitzer, M. et al. A brief history of human disease genetics. Nature 577, 179–189 (2020).

2. Maurano, M. T. et al. Systematic localization of common disease-associated variation in regulatory DNA. Science 337, 1190–1195 (2012).

3. GTEx Consortium. The GTEx Consortium atlas of genetic regulatory effects across human tissues. Science 369, 1318–1330 (2020).

4. Roadmap Epigenomics Consortium et al. Integrative analysis of 111 reference human epigenomes. Nature 518, 317–330 (2015).

5. ENCODE Project Consortium et al. Expanded encyclopaedias of DNA elements in the human and mouse genomes. Nature 583, 699–710 (2020).

6. Zheng, Y. et al. Role of conserved non-coding DNA elements in the Foxp3 gene in regulatory T-cell fate. Nature 463, 808–812 (2010).

7. Christmas, M. J. et al. Evolutionary constraint and innovation across hundreds of placental mammals. Science 380, eabn3943 (2023).

8. Sullivan, P. F. et al. Leveraging base-pair mammalian constraint to understand genetic variation and human disease. Science 380, eabn2937 (2023).

9. Bejerano, G. et al. Ultraconserved elements in the human genome. Science 304, 1321– 1325 (2004).

10. Kellis, M. et al. Defining functional DNA elements in the human genome. Proc. Natl. Acad. Sci. U. S. A. 111, 6131–6138 (2014).

11. Cano-Gamez, E. & Trynka, G. From GWAS to Function: Using Functional Genomics to Identify the Mechanisms Underlying Complex Diseases. Front. Genet. 11, 424 (2020).

12. Visscher, P. M. et al. 10 Years of GWAS Discovery: Biology, Function, and Translation. Am. J. Hum. Genet. 101, 5–22 (2017).

13. Farh, K. K.-H. et al. Genetic and epigenetic fine mapping of causal autoimmune disease variants. Nature 518, 337–343 (2015).

14. Zhang, K. et al. A single-cell atlas of chromatin accessibility in the human genome. Cell 184, 5985–6001.e19 (2021).

15. Domcke, S. et al. A human cell atlas of fetal chromatin accessibility. Science 370, eaba7612 (2020).

16. Buenrostro, J. D. et al. Integrated Single-Cell Analysis Maps the Continuous Regulatory Landscape of Human Hematopoietic Differentiation. Cell 173, 1535–1548.e16 (2018).

17. Zhu, K. et al. Multi-omic profiling of the developing human cerebral cortex at the single-cell level. Sci. Adv. 9, eadg3754 (2023).

18. Mannens, C. C. A. et al. Chromatin accessibility during human first-trimester neurodevelopment. Nature (2024) doi:10.1038/s41586-024-07234-1.

19. Jiang, X., Holmes, C. & McVean, G. The impact of age on genetic risk for common diseases. PLoS Genet. 17, e1009723 (2021).

20. van der Wijst, M. et al. The single-cell eQTLGen consortium. eLife 9, e52155 (2020).

21. Ding, R. et al. scQTLbase: an integrated human single-cell eQTL database. Nucleic Acids Res. 52, D1010–D1017 (2024).

22. Tewhey, R. et al. Direct Identification of Hundreds of Expression-Modulating Variants using a Multiplexed Reporter Assay. Cell 165, 1519–1529 (2016).

23. Martin-Rufino, J. D. et al. Massively parallel base editing to map variant effects in human hematopoiesis. Cell 186, 2456–2474.e24 (2023).

24. Mahajan, A. et al. Fine-mapping type 2 diabetes loci to single-variant resolution using high-density imputation and islet-specific epigenome maps. Nat. Genet. 50, 1505–1513 (2018).

25. Connally, N. J. et al. The missing link between genetic association and regulatory function. eLife 11, e74970 (2022).

26. Mostafavi, H., Spence, J. P., Naqvi, S. & Pritchard, J. K. Systematic differences in discovery of genetic effects on gene expression and complex traits. Nat. Genet. 55, 1866–1875 (2023).

27. Hawkes, G. et al. Whole-genome sequencing in 333,100 individuals reveals rare non-coding single variant and aggregate associations with height. Nat. Commun. 15, 8549 (2024).

28. Selvaraj, M. S. et al. Whole genome sequence analysis of blood lipid levels in >66,000 individuals. Nat. Commun. 13, 5995 (2022).

29. Ferraro, N. M. et al. Transcriptomic signatures across human tissues identify functional rare genetic variation. Science 369, eaaz5900 (2020).

30. Tardaguila, M. et al. Variant-to-function dissection of rare non-coding GWAS loci with high impact on blood traits. Preprint at 10.1101/2024.08.05.606572 (2024).

31. Ribeiro, D. M. & Delaneau, O. Non-coding rare variant associations with blood traits on 166 740 UK Biobank genomes. Preprint at 10.1101/2023.12.01.569422 (2023).

32. Kuderna, L. F. K. et al. Identification of constrained sequence elements across 239 primate genomes. Nature 625, 735–742 (2024).

33. Dubovik, T. et al. Interactions between immune cell types facilitate the evolution of immune traits. Nature 632, 350–356 (2024).

34. Quintana-Murci, L. Human Immunology through the Lens of Evolutionary Genetics. Cell 177, 184–199 (2019).

35. Avsec, Ž., et al. Effective gene expression prediction from sequence by integrating long-range interactions. Nat. Methods 18, 1196–1203 (2021).

36. Avsec, Ž., et al. Base-resolution models of transcription-factor binding reveal soft motif syntax. Nat. Genet. 53, 354–366 (2021).

37. Zhou, J. & Troyanskaya, O. G. Predicting effects of noncoding variants with deep learning-based sequence model. Nat. Methods 12, 931–934 (2015).

38. Chen, K. M., Wong, A. K., Troyanskaya, O. G. & Zhou, J. A sequence-based global map of regulatory activity for deciphering human genetics. Nat. Genet. 54, 940–949 (2022).

39. Maslova, A. et al. Deep learning of immune cell differentiation. Proc. Natl. Acad. Sci. U. S. A. 117, 25655–25666 (2020).

40. Minnoye, L. et al. Cross-species analysis of enhancer logic using deep learning. Genome Res. 30, 1815–1834 (2020).

41. Dudnyk, K., Cai, D., Shi, C., Xu, J. & Zhou, J. Sequence basis of transcription initiation in the human genome. Science 384, eadj0116 (2024).

42. Hingerl, J. C. et al. scooby: Modeling multi-modal genomic profiles from DNA sequence at single-cell resolution. BioRxiv Prepr. Serv. Biol. 2024.09.19.613754 (2024) doi:10.1101/2024.09.19.613754.

43. Pampari, A. et al. ChromBPNet: bias factorized, base-resolution deep learning models of chromatin accessibility reveal cis-regulatory sequence syntax, transcription factor footprints and regulatory variants. BioRxiv Prepr. Serv. Biol. 2024.12.25.630221 (2025) doi:10.1101/2024.12.25.630221.

44. Trevino, A. E. et al. Chromatin and gene-regulatory dynamics of the developing human cerebral cortex at single-cell resolution. Cell 184, 5053–5069.e23 (2021).

45. Ameen, M. et al. Integrative single-cell analysis of cardiogenesis identifies developmental trajectories and non-coding mutations in congenital heart disease. Cell 185, 4937–4953.e23 (2022).

46. Shrikumar, A., Greenside, P. & Kundaje, A. Learning Important Features Through Propagating Activation Differences. (2017) doi:10.48550/ARXIV.1704.02685.

47. Lundberg, S. & Lee, S.-I. A Unified Approach to Interpreting Model Predictions. Preprint at 10.48550/ARXIV.1705.07874 (2017).

48. Shrikumar, A. et al. Technical Note on Transcription Factor Motif Discovery from Importance Scores (TF-MoDISco) version 0.5.6.5. Preprint at 10.48550/ARXIV.1811.00416 (2018).

49. Byrska-Bishop, M. et al. High-coverage whole-genome sequencing of the expanded 1000 Genomes Project cohort including 602 trios. Cell 185, 3426–3440.e19 (2022).

50. Corces, M. R. et al. Single-cell epigenomic analyses implicate candidate causal variants at inherited risk loci for Alzheimer’s and Parkinson’s diseases. Nat. Genet. 52, 1158–1168 (2020).

51. Kierdorf, K. et al. Microglia emerge from erythromyeloid precursors via Pu.1- and Irf8-dependent pathways. Nat. Neurosci. 16, 273–280 (2013).

52. Lyons, G. E., Micales, B. K., Schwarz, J., Martin, J. F. & Olson, E. N. Expression of mef2 genes in the mouse central nervous system suggests a role in neuronal maturation. J. Neurosci. Off. J. Soc. Neurosci. 15, 5727–5738 (1995).

53. Bedogni, F. et al. Tbr1 regulates regional and laminar identity of postmitotic neurons in developing neocortex. Proc. Natl. Acad. Sci. U. S. A. 107, 13129–13134 (2010).

54. Pataskar, A. et al. NeuroD1 reprograms chromatin and transcription factor landscapes to induce the neuronal program. EMBO J. 35, 24–45 (2016).

55. Yamada, M. et al. Specification of spatial identities of cerebellar neuron progenitors by ptf1a and atoh1 for proper production of GABAergic and glutamatergic neurons. J. Neurosci. Off. J. Soc. Neurosci. 34, 4786–4800 (2014).

56. Veyrac, A. et al. Zif268/egr1 gene controls the selection, maturation and functional integration of adult hippocampal newborn neurons by learning. Proc. Natl. Acad. Sci. U. S. A. 110, 7062–7067 (2013).

57. Miller, J. A. et al. Transcriptional landscape of the prenatal human brain. Nature 508, 199–206 (2014).

58. Fauman, E. B. & Hyde, C. An optimal variant to gene distance window derived from an empirical definition of cis and trans protein QTLs. BMC Bioinformatics 23, 169 (2022).

59. Meylan, P., Dreos, R., Ambrosini, G., Groux, R. & Bucher, P. EPD in 2020: enhanced data visualization and extension to ncRNA promoters. Nucleic Acids Res. 48, D65–D69 (2020).

60. Kim-Hellmuth, S. et al. Cell type-specific genetic regulation of gene expression across human tissues. Science 369, eaaz8528 (2020).

61. Finucane, H. K. et al. Heritability enrichment of specifically expressed genes identifies disease-relevant tissues and cell types. Nat. Genet. 50, 621–629 (2018).

62. Kim, S. S. et al. Leveraging single-cell ATAC-seq and RNA-seq to identify disease-critical fetal and adult brain cell types. Nat. Commun. 15, 563 (2024).

63. Bellenguez, C. et al. New insights into the genetic etiology of Alzheimer’s disease and related dementias. Nat. Genet. 54, 412–436 (2022).

64. Watson, H. J. et al. Genome-wide association study identifies eight risk loci and implicates metabo-psychiatric origins for anorexia nervosa. Nat. Genet. 51, 1207–1214 (2019).

65. Wang, Q. S. et al. Leveraging supervised learning for functionally informed fine-mapping of cis-eQTLs identifies an additional 20,913 putative causal eQTLs. Nat. Commun. 12, 3394 (2021).

66. Aragam, K. G. et al. Discovery and systematic characterization of risk variants and genes for coronary artery disease in over a million participants. Nat. Genet. 54, 1803–1815 (2022).

67. Tcheandjieu, C. et al. Large-scale genome-wide association study of coronary artery disease in genetically diverse populations. Nat. Med. 28, 1679–1692 (2022).

68. Zou, Y., Carbonetto, P., Wang, G. & Stephens, M. Fine-mapping from summary data with the ‘Sum of Single Effects’ model. PLoS Genet. 18, e1010299 (2022).

69. Cheng, P. et al. ZEB2 Shapes the Epigenetic Landscape of Atherosclerosis. Circulation 145, 469–485 (2022).

70. Abell, N. S. et al. Multiple causal variants underlie genetic associations in humans. Science 375, 1247–1254 (2022).

71. Wu, Z. et al. PICALM rs3851179 Variants Modulate Left Postcentral Cortex Thickness, CSF Amyloid β42, and Phosphorylated Tau in the Elderly. Brain Sci. 12, 1681 (2022).

72. Wu, Z. et al. The Effects of PICALM rs3851179 and Age on Brain Atrophy and Cognition Along the Alzheimer’s Disease Continuum. Mol. Neurobiol. 61, 6984–6996 (2024).

73. Santos-Rebouças, C. B. et al. rs3851179 Polymorphism at 5’ to the PICALM Gene is Associated with Alzheimer and Parkinson Diseases in Brazilian Population. Neuromolecular Med. 19, 293–299 (2017).

74. Sun, D.-M. et al. Effect of PICALM rs3851179 polymorphism on the default mode network function in mild cognitive impairment. Behav. Brain Res. 331, 225–232 (2017).

75. Liu, G. et al. Lack of association between PICALM rs3851179 polymorphism and Alzheimer’s disease in Chinese population and APOEε4-negative subgroup. Neurobiol. Aging 34, 1310.e9–10 (2013).

76. Socodato, R. et al. Microglia Dysfunction Caused by the Loss of Rhoa Disrupts Neuronal Physiology and Leads to Neurodegeneration. Cell Rep. 31, 107796 (2020).

77. Chen, M. J. et al. Extracellular signal-regulated kinase regulates microglial immune responses in Alzheimer’s disease. J. Neurosci. Res. 99, 1704–1721 (2021).

78. Fallerini, C. et al. Common, low-frequency, rare, and ultra-rare coding variants contribute to COVID-19 severity. Hum. Genet. 141, 147–173 (2022).

79. Pollard, K. S., Hubisz, M. J., Rosenbloom, K. R. & Siepel, A. Detection of nonneutral substitution rates on mammalian phylogenies. Genome Res. 20, 110–121 (2010).

80. Coetzee, S. G., Coetzee, G. A. & Hazelett, D. J. motifbreakR: an R/Bioconductor package for predicting variant effects at transcription factor binding sites. Bioinforma. Oxf. Engl. 31, 3847–3849 (2015).

81. Yan, J. et al. Systematic analysis of binding of transcription factors to noncoding variants. Nature 591, 147–151 (2021).

82. Niemi, M. E. K. et al. Common genetic variants contribute to risk of rare severe neurodevelopmental disorders. Nature 562, 268–271 (2018).

83. Wang, P., Zhao, D., Lachman, H. M. & Zheng, D. Enriched expression of genes associated with autism spectrum disorders in human inhibitory neurons. Transl. Psychiatry 8, 13 (2018).

84. Fu, J. M. et al. Rare coding variation provides insight into the genetic architecture and phenotypic context of autism. Nat. Genet. 54, 1320–1331 (2022).

85. Shin, T. et al. Rare variation in non-coding regions with evolutionary signatures contributes to autism spectrum disorder risk. Cell Genomics 4, 100609 (2024).

86. An, J.-Y. et al. Genome-wide de novo risk score implicates promoter variation in autism spectrum disorder. Science 362, eaat6576 (2018).

87. Padhi, E. M. et al. Coding and noncoding variants in EBF3 are involved in HADDS and simplex autism. Hum. Genomics 15, 44 (2021).

88. Short, P. J. et al. De novo mutations in regulatory elements in neurodevelopmental disorders. Nature 555, 611–616 (2018).

89. Rentzsch, P., Witten, D., Cooper, G. M., Shendure, J. & Kircher, M. CADD: predicting the deleteriousness of variants throughout the human genome. Nucleic Acids Res. 47, D886– D894 (2019).

90. Florian, C., Bahi-Buisson, N. & Bienvenu, T. FOXG1-Related Disorders: From Clinical Description to Molecular Genetics. Mol. Syndromol. 2, 153–163 (2012).

91. Harris, H. K. et al. Disruption of RFX family transcription factors causes autism, attention-deficit/hyperactivity disorder, intellectual disability, and dysregulated behavior. Genet. Med. Off. J. Am. Coll. Med. Genet. 23, 1028–1040 (2021).

92. Rodenas-Cuadrado, P., Ho, J. & Vernes, S. C. Shining a light on CNTNAP2: complex functions to complex disorders. Eur. J. Hum. Genet. EJHG 22, 171–178 (2014).

93. Steele-Perkins, G. et al. The transcription factor gene Nfib is essential for both lung maturation and brain development. Mol. Cell. Biol. 25, 685–698 (2005).

94. Betancourt, J., Katzman, S. & Chen, B. Nuclear factor one B regulates neural stem cell differentiation and axonal projection of corticofugal neurons. J. Comp. Neurol. 522, 6–35 (2014).

95. Schanze, I. et al. NFIB Haploinsufficiency Is Associated with Intellectual Disability and Macrocephaly. Am. J. Hum. Genet. 103, 752–768 (2018).

96. Cuomo, A. S. E. et al. Single-cell RNA-sequencing of differentiating iPS cells reveals dynamic genetic effects on gene expression. Nat. Commun. 11, 810 (2020).

97. Yazar, S. et al. Single-cell eQTL mapping identifies cell type-specific genetic control of autoimmune disease. Science 376, eabf3041 (2022).

98. Emani, P. S. et al. Single-cell genomics and regulatory networks for 388 human brains. Science 384, eadi5199 (2024).

99. Libé-Philippot, B. & Vanderhaeghen, P. Cellular and Molecular Mechanisms Linking Human Cortical Development and Evolution. Annu. Rev. Genet. 55, 555–581 (2021).

100. Spalding, K. L. et al. Dynamics of hippocampal neurogenesis in adult humans. Cell 153, 1219–1227 (2013).

101. Koketsu, D., Mikami, A., Miyamoto, Y. & Hisatsune, T. Nonrenewal of neurons in the cerebral neocortex of adult macaque monkeys. J. Neurosci. Off. J. Soc. Neurosci. 23, 937–942 (2003).

102. Yamamoto, R. et al. Tissue-specific impacts of aging and genetics on gene expression patterns in humans. Nat. Commun. 13, 5803 (2022).

103. Turan, Z. G. et al. Molecular footprint of Medawar’s mutation accumulation process in mammalian aging. Aging Cell 18, e12965 (2019).

104. Huang, Q. Q. et al. Examining the role of common variants in rare neurodevelopmental conditions. Nature 636, 404–411 (2024).

105. Turner, T. N. & Eichler, E. E. The Role of De Novo Noncoding Regulatory Mutations in Neurodevelopmental Disorders. Trends Neurosci. 42, 115–127 (2019).

106. Akdemir, K. C. et al. Somatic mutation distributions in cancer genomes vary with three-dimensional chromatin structure. Nat. Genet. 52, 1178–1188 (2020).

107. Marderstein, A. R. et al. Single-cell multi-omics map of human fetal blood in Down syndrome. Nature 634, 104–112 (2024).

108. Wang, Z. et al. The Value of Rare Genetic Variation in the Prediction of Common Obesity in European Ancestry Populations. Front. Endocrinol. 13, 863893 (2022).

109. Kim, Y. et al. CWAS-Plus: estimating category-wide association of rare noncoding variation from whole-genome sequencing data with cell-type-specific functional data. Brief. Bioinform. 25, bbae323 (2024).

110. Smail, C. et al. Integration of rare expression outlier-associated variants improves polygenic risk prediction. Am. J. Hum. Genet. 109, 1055–1064 (2022).

111. Coorens, T. H. H. et al. The human and non-human primate developmental GTEx projects. Nature 637, 557–564 (2025).

112. IGVF Consortium. Deciphering the impact of genomic variation on function. Nature 633, 47–57 (2024).

113. Barral, A. & Zaret, K. S. Pioneer factors: roles and their regulation in development. Trends Genet. TIG 40, 134–148 (2024).

114. Li, T. et al. The functional impact of rare variation across the regulatory cascade. Cell Genomics 3, 100401 (2023).

115. Nasser, J. et al. Genome-wide enhancer maps link risk variants to disease genes. Nature 593, 238–243 (2021).

116. Mitra, S. et al. Single-cell multi-ome regression models identify functional and disease-associated enhancers and enable chromatin potential analysis. Nat. Genet. 56, 627–636 (2024).

117. Sakaue, S. et al. Tissue-specific enhancer-gene maps from multimodal single-cell data identify causal disease alleles. Nat. Genet. 56, 615–626 (2024).

118. Rozowsky, J. et al. The EN-TEx resource of multi-tissue personal epigenomes & variant-impact models. Cell 186, 1493–1511.e40 (2023).

119. Zhang, Y. et al. Model-based analysis of ChIP-Seq (MACS). Genome Biol. 9, R137 (2008).

120. Purcell, S. et al. PLINK: a tool set for whole-genome association and population-based linkage analyses. Am. J. Hum. Genet. 81, 559–575 (2007).

121. Kuhn, R. M., Haussler, D. & Kent, W. J. The UCSC genome browser and associated tools. Brief. Bioinform. 14, 144–161 (2013).

122. McLaren, W. et al. The Ensembl Variant Effect Predictor. Genome Biol. 17, 122 (2016).

123. Karczewski, K. J. et al. The mutational constraint spectrum quantified from variation in 141,456 humans. Nature 581, 434–443 (2020).

124. Zeng, T., Spence, J. P., Mostafavi, H. & Pritchard, J. K. Bayesian estimation of gene constraint from an evolutionary model with gene features. Nat. Genet. 56, 1632–1643 (2024).

125. Rauluseviciute, I. et al. JASPAR 2024: 20th anniversary of the open-access database of transcription factor binding profiles. Nucleic Acids Res. 52, D174–D182 (2024).

126. Weirauch, M. T. et al. Determination and inference of eukaryotic transcription factor sequence specificity. Cell 158, 1431–1443 (2014).

127. Vorontsov, I. E. et al. HOCOMOCO in 2024: a rebuild of the curated collection of binding models for human and mouse transcription factors. Nucleic Acids Res. 52, D154–D163 (2024).

128. Vierstra, J. et al. Global reference mapping of human transcription factor footprints. Nature 583, 729–736 (2020).

129. Lee, Y., Luca, F., Pique-Regi, R. & Wen, X. Bayesian Multi-SNP Genetic Association Analysis: Control of FDR and Use of Summary Statistics. Preprint at 10.1101/316471 (2018).

130. Davis, J. R. et al. An Efficient Multiple-Testing Adjustment for eQTL Studies that Accounts for Linkage Disequilibrium between Variants. Am. J. Hum. Genet. 98, 216–224 (2016).

131. Lee, D. et al. A method to predict the impact of regulatory variants from DNA sequence. Nat. Genet. 47, 955–961 (2015).

132. Tarailo-Graovac, M. & Chen, N. Using RepeatMasker to identify repetitive elements in genomic sequences. Curr. Protoc. Bioinforma. Chapter 4, 4.10.1–4.10.14 (2009).

