## Supplementary Information for "Mapping the regulatory effects of common and rare non-coding variants across cellular and developmental contexts in the brain and heart"

### Supplementary Notes

**Supplementary Note 1: Removal of cell-types with poor ChromBPNet performance.**

To build our variant prediction resource, we harmonized and curated pseudobulk ATAC-seq data from 134 cellular contexts across five studies, including 30 from fetal brain, 24 from adult brain, 33 from fetal heart, and 47 from adult heart, and trained ChromBPNet models for each. We evaluated model performance using Spearman's correlation between predicted and observed ATAC read counts within peak regions, removing two adult heart cell types with outlier model performance (>4 standard deviations). The correlations between observed and predicted Tn5 insertion counts within peaks ranged from 0.44 to 0.83 (median 0.71, mean 0.70) for the final 132 cell types kept for downstream analyses (Supplementary Figure 1).

**Supplementary Note 2: Relationship between cell-type-specificity and magnitude.**

We observed a positive correlation between regulatory magnitude (defined as the maximum predicted effect across cell types) and cell-type-specificity (measured as the predicted number of affected cell types). This correlation may reflect a phenomenon akin to winner's curse, where the largest-effect variants are more likely to exhibit effects in multiple cell types due to shared regulatory syntax. This shared syntax may enable weaker effects in additional cell types to surpass the threshold for classification as an "affected" cell type. To address potential confounders in this analysis, we conducted a series of sensitivity analyses.

1. Restricting to Accessible Variants: We limited our analysis to variants predicted to have effects in at least one cell type. This restriction had minimal impact on the results, as our original analysis already focused exclusively on accessible variants.
2. Using Mean Effects for Regulatory Magnitude: We re-estimated regulatory magnitude using the mean predicted effect across cell types instead of the maximum. The association between regulatory magnitude and cell-type-specificity was even stronger when using mean effects. Variants with consistently strong mean effects across cell types are more likely to impact a higher number of cell types, reinforcing the observed correlation.
3. Focusing on Variants Near Highly Constrained Genes: We restricted our analysis to variants near genes with high constraint, using shet scores. We tested thresholds for constraint by selecting genes in the top 10%, 5%, and 1% of constraint. Even with these restrictions, we observed robust correlations between regulatory magnitude and cell-type-specificity.
4. Varying the Threshold for Defining Affected Cell Types: In the initial analysis, we classified a cell type as affected if the ChromBPNet empirical *P*-value $< 0.01$. To assess sensitivity to this threshold, we repeated the analysis with thresholds of $P < 0.1$ and $P < 0.001$. Across all thresholds, the correlation between regulatory magnitude and cell-type-specificity remained strong.

These sensitivity analyses demonstrate that the observed association between cell-type-specificity and regulatory magnitude is robust to methodological choices and threshold variations. While this association may partly result from winner's curse, it remains consistent across a range of analytic frameworks and assumptions.

**Supplementary Note 3: Putative causal variants at the Alzheimer's *PICALM* locus.**

We found evidence for multiple putative causal variants at the non-coding Alzheimer's disease GWAS locus near *PICALM*, challenging the conception of a single underlying causal variant[1–6]. At this locus, there is extensive linkage disequilibrium spanning ~200 kb region, with 9,794 genome-wide significant variants ($P < 5x10^{-8}$) within 1 mb of the sentinel variant rs3851179. Fine-mapping of this locus identified two credible sets with a 95% posterior probability of containing a causal variant, which included two and nine variants respectively. By intersecting the credible sets with our scATAC-seq data, we discovered two variants (one in each credible set) that were accessible in microglia and for which the alternate allele was predicted to disrupt accessibility: rs3844143 and rs10792832. Neither of the two identified variants were accessible (or functional) in any cell type found in the adult brain besides microglia, but one of these two variants (rs10792832) is accessible and predicted to reduce accessibility in fetal brain microglia and fetal heart myeloid cells as well. For both variants, the predicted effects were small but significant (ChromBPNet *P*-values < 0.05). Interestingly, our models did not predict the sentinel variant rs3851179 (PIP = 0.50) to be accessible or alter accessibility, despite being extensively studied previously across populations and phenotypes[2–6]. Our results suggests that, at this GWAS locus, there are two putative causal variants, one disrupting a FOXN1 motif in myeloid cells across development and organ contexts (PIP = 0.50) with the other specifically active later in aging within adult microglia (PIP = 0.26) (Supplementary Figure 6A).

**Supplementary Note 4: Sensitivity analysis of common versus rare comparison.**

We reasoned that since the ChromBPNet analyses only test variants accessible within open chromatin regions, the set of variants analyzed could differ between cell types. If peaks in one cell type are more functionally relevant and thus under stronger evolutionary constraint, ChromBPNet scores in that cell type would be expected to correlate more strongly with measures of constraint, such as minor allele frequencies. A potential confounder in downstream analyses is that a small regulatory effect in a peak within a critical cell type (e.g., fetal brain neurons) may have evolutionary relevance, while the same regulatory effect size in a non-critical peak in another cell type may not.

To address this, we tested the robustness of our results by focusing on our top-ranked cell type, excitatory neurons in the fetal brain[7]. For each additional cell type, we performed the following steps:

1. *Identified a shared set of accessible variants*: We restricted the analysis to variants that were jointly accessible in both the top-ranked cell type (fetal brain excitatory neurons) and the cell type of interest.
2. *Reanalyzed ChromBPNet scores*: We repeated the common vs. rare ChromBPNet score analysis (see Methods) for both cell types using this limited set of jointly accessible variants, calculating effect sizes that reflect the mean difference in predicted effects between common and rare variants.
3. *Quantified differences in effect sizes*: We calculated the difference in effect sizes between the two cell-type analyses, along with 95% confidence intervals for the differences.

By limiting the ChromBPNet analysis to a shared set of peaks between the top-ranked and the other cell type, we ensured that differences in the peak set did not confound the results.

Across all comparisons, fetal brain excitatory neurons consistently showed a stronger correlation between ChromBPNet scores and constraint compared to the other cell type, as the effect size difference was always greater than zero, even when using the same set of peaks for both cell types (Supplementary Figure 8). For example, when restricting the variant set to 275,654 variants accessible in both isocortical excitatory neurons in the adult brain (Corces et al. 2020[8]) and excitatory neurons in the fetal brain (Domcke et al. 2020[7]), we found that the difference in ChromBPNet scores between common and rare variants was significantly larger in the fetal brain compared to the adult brain ($P = 2.4 \times 10^{-6}$). Notably, beyond glutamatergic neuron 3 in fetal brain (Trevino et al. 2021[9]), this was the greatest number of jointly accessible variants (275,654 variants) compared to any other cell context (median = 111,668 variants), suggesting this is the most similar non-fetal brain context in our single-cell ATAC-seq dataset. We observed similar findings using 255,024 jointly accessible variants from adult brain hippocampal excitatory neuron 2 (larger ChromBPNet differences in fetal brain excitatory neurons, $P = 2.6 \times 10^{-7}$).

These results further support the robustness of our observation that fetal brain neurons play a greater role in evolutionary constraint than other cell types, highlighting their critical functional relevance.

**Supplementary Note 5: Constraint on overall accessibility.**

We assessed differences between common and rare variants for their predicted ChromBPNet effects in each cell type, focusing on accessible variants and using a linear model. Separately, we analyzed the proportion of accessible variants for common versus rare variants across cell types, using genome-wide variants and a logistic regression model. Notably, the ChromBPNet analysis revealed a striking enrichment in fetal brain neurons, although the peaks-based analysis showed a somewhat noisier enrichment (Supplementary Figure 7, 9).

To evaluate the consistency between these two approaches, we quantified the similarity of their findings by correlating the effect sizes estimated from the ChromBPNet models with those from the peaks analysis. The two analyses were moderately correlated, with a Pearson correlation coefficient of $r = 0.40$ ($P = 2.5 \times 10^{-6}$) (Supplementary Figure 9). Fetal brain neurons consistently showed the strongest differences between rare and common variants in both analyses, while adult brain cells exhibited relatively minor differences. Fetal and adult heart cell types showed greater heterogeneity, with large differences observed at the peak level but occasionally smaller differences in the ChromBPNet analysis. Overall, these results suggest that the two analyses yield broadly consistent conclusions while highlighting subtle differences in specific cell types.

**Supplementary Note 6: AP-1 and EGR motifs and their role in fetal neuron accessibility.**

Using our ChromBPNet models trained on adult and fetal neuron ATAC-seq data, we observed notable cell type- and developmental stage-specific differences in transcription factor motif usage (see Methods). We generated TF-MoDISco motif reports for each dataset and cell type (adult excitatory neurons from Corces et al. 2020; fetal excitatory neurons from Trevino et al. 2021 and Domcke et al. 2020). These reports confirmed that AP-1 and EGR motifs were among the top learned patterns in adult neurons, whereas in fetal excitatory neurons, these motifs did not appear at all in the top motif clusters. For example, the TGAgTCA motif for AP-1 is one of the top detected patterns in adult excitatory neurons but was virtually absent in fetal excitatory neurons, suggesting that it is more functionally relevant in driving open chromatin regions of adult neurons compared to fetal neurons. Both AP-1 and EGR are known to be involved in activity-dependent gene regulation, neuronal plasticity, and stress responses—processes that become more critical in the mature nervous system. EGR1, for instance, has established roles in learning and memory[10,11], further highlighting its potential importance in adult rather than embryonic neural contexts. The absence of AP-1 and EGR motifs in fetal neurons supports the notion that different TF networks dominate early developmental programs versus adult neuron function.

**Supplementary Note 7: TF binding predictions using SNP-SELEX.**

We employed deltaSVM[12] to score all 1,617 fetal- and adult-specific excitatory neuron variants for their predicted effect on transcription factor (TF) binding across 94 TFs. Prior to enrichment analysis, all deltaSVM scores were standardized to have mean equal to 0 and standard deviation equal to 1 for each TF. We focused on variants that both overlapped predicted TF binding sites (as predicted by deltaSVM using the seq_binding model prediction) and met a stringent threshold (z-score ≥ 2) relative to each TF's average deltaSVM distribution. As illustrated in Supplementary Figure 12B, we observed clear enrichment of TF patterns between fetal- and adult-specific variants at this higher threshold, indicating strong predicted differences in TF binding for these two groups of variants. Additionally, when relaxing this threshold (z-score ≥ 1), we observed similar results. We then qualitatively analyzed our TF enrichments along with their expression patterns in the developing brain using the BrainSpan[13] web interface and different regions of the brain using the GTEx portal[14] (Supplementary Table 13, Supplementary Figure 12B).

Among the top TFs with consistent enrichment of fetal-specific variant effects (z-score ≥ 2), we identified NFE2, which is expressed during early and late fetal stages in the developing human brain, primarily in the cerebellar hemisphere and cerebellum. Other top enriched TFs for fetal-specific variants included ATOH1—expressed in the fetal cerebellum and enriched in the motif-centered analysis—and OLIG3.

In the adult-specific variant set, TFs such as NFE2, EGR1, EGR3, and JDP2 were enriched for adult-specific variant effects. NFE2, noted for its cerebellar expression and role as a negative regulator of proliferation, has also been found at composite binding sites with AP-1 TFs[15], which was enriched at the motif-centered analysis. Additionally, the observation of EGR1 supported the motifbreakR results and exhibits higher postnatal and adult brain expression in BrainSpan, while JDP2 expression is similarly observed during postnatal and adult stages.

**Supplementary Note 8: FLARE in additional variant sets.**

We further evaluated FLARE using (i) the curated common variant dataset from 1KG, and (ii) a dataset of 84,980 de novo non-coding variants from whole-genome sequencing of 1,902 autism spectrum disorder (ASD) simplex families[9,16] (42,931 variants from ASD cases and 42,049 from controls), comparing FLARE predictions to PhyloP. We found that FLARE was a weaker predictor for common variants and a stronger predictor for de novo mutations, despite the genomic context (FLARE-baseline) explaining reduced PhyloP variance in de novo variants (Supplementary Figure 15A). Further, the differences of FLARE-fb scores between variant sets (common vs. rare vs. ASD) was larger than for FLARE-ab or -heart (Supplementary Figure 15B). Finally, we observed a 66.5% improvement of FLARE-fb over FLARE-fb-peaks for ASD de novo mutations compared to only 11.6% improvement for common variants (Supplementary Figure 15A), suggesting that ChromBPNet was particularly useful for interpreting increasingly rare polymorphisms. The de novo nature of these variants, which are more likely to be damaging and have not been subjected to generational selective pressures, likely explains the stronger association between FLARE scores and PhyloP in the ASD set compared to common and rare inherited variants observed in 1KG. Overall, FLARE models informed context-specific regulatory evolution.

### Supplementary Figure Legends

**Supplementary Figure 1: Dataset characteristics.**
(A) Example comparison of the measured and predicted log (natural log, base e) of total counts in peak regions. Each point represents a different peak in cluster c0 from Trevino et al. (a fetal excitatory neuron context), with predictions made using the ChromBPNet fold 0 model.
(B) Spearman correlation of measured and predicted log (natural log, base e) in peak regions. Models were trained and tested over 5 folds. Each point represents a different model fold. The barplots reflect the mean correlation values across folds.

**Supplementary Figure 2: Analysis of cell-type-specificity.**
(A) Number of rare variants in the different cell-type-specificity sets of "null", "specific", "multiple", or "shared".
(B) Histogram of number of affected cell types for rare variants.
(C) Histogram of regulatory magnitude (maximum ChromBPNet score across cell types) stratified by cell-type-specificity variant sets.

**Supplementary Figure 3: Fine-mapped eQTLs are associated with broad effects.** Fine-mapped eQTLs in GTEx brain (bottom) and heart (top) samples are associated with variants that are associated with multiple cell types (according to ChromBPNet empirical *P*-value < 0.01). A linear model was performed for each organ context separately, and the *P*-values describing the enrichment for fine-mapped eQTLs relative to non-fine-mapped eQTLs is shown.

**Supplementary Figure 4: Cell-type-specific GWAS heritability enrichments.**
(A) Heritability enrichments for 8 cardiovascular and neurological disorders are shown across cell types and developmental contexts (encompassing 9 studies, as coronary artery disease (CAD) is shown for two large GWAS meta-analyses). Cell types are separated by organ and developmental context and the original scATAC published study. Each tile represents the significance, which is either "Not Sig" (nominal P > 0.05), "P < 0.05" (nominal), "FDR < 0.1", "FDR < 0.05"< "FDR < 0.01" (referring to significance values after correcting for multiple tests).
(B-C) Highlighted results include adult microglia enrichment for Alzheimer's disease and fetal neurons for anorexia nervosa. Red line refers to Bonferroni significance.

**Supplementary Figure 5: ChromBPNet enrichments in two CAD GWAS.**
Correlation of fine-mapped variant enrichments for ChromBPNet between two distinct coronary artery disease (CAD) GWAS datasets. A linear model was performed for each cell type separately, associating a binarized covariate for fine-mapped eQTLs relative to non-fine-mapped eQTLs with ChromBPNet scores across all accessible variants. ChromBPNet scores were scaled to have mean equal to 0 and standard deviation equal to 1 prior to analysis. The effect size of the binarized covariate is shown along the axes.

**Supplementary Figure 6: Microglia effects at Alzheimer's disease GWAS loci.**
(A) Variant effect predictions for three fine-mapped variants at a non-coding Alzheimer's disease GWAS locus near the *PICALM* gene. The top two and bottom two plots show chromatin

profile predictions for the three variants in a 300-bp window surrounding each variant with the predicted ATAC counts along the *y*-axis. Each colored line represents the base-pair resolution prediction for an input sequence containing the reference or alternate alleles, with the top plot made for fetal brain microglia and the other three plots made for adult brain microglia. The middle two plots show the output from DeepLIFT for the first variant (rs10792832), which reflect contribution scores for each base in the sequence, along the *y*-axis.
(B) Gene set enrichment analysis (GSEA) of candidate genes for GWAS variants (x) compared to a GSEA of candidate genes for GWAS variants with a predicted effect in microglia (y). Each point is a distinct gene set. The axes show $-\log_{10}$ FDR values, and Pearson correlation was calculated between the significance values for the two analyses.

**Supplementary Figure 7: Common versus rare variant analyses.**
The *x*-axis shows effect sizes estimating the mean differences in ChromBPNet scores (left) and peak accessibility (right) for common versus rare variants in the respective analyses.

**Supplementary Figure 8: Restricting common versus rare analyses to identical variant sets.**
To address potential confounding from differences in the variants analyzed across cell types, we performed a robustness analysis focused on fetal brain excitatory neurons, the top-ranked cell type. For each additional cell type, we restricted the analysis to variants jointly accessible in both fetal brain excitatory neurons and the cell type of interest. Using this shared set of variants, we compared ChromBPNet-predicted regulatory effects between common and rare variants. Effect sizes were calculated as the mean difference in predicted effects for each cell type, and differences between cell types were quantified as the difference in effect sizes with 95% confidence intervals. This approach, using a shared set of peaks, controlled for confounding from accessibility differences, ensuring observed effects reflected true cell-type-specific regulatory differences. The figure shows the differences in effect sizes with 95% confidence intervals, sorted from low to high.

**Supplementary Figure 9: Proportions of accessible rare and common variants.**
(A) Analysis of the distribution of rare versus common variants in accessible chromatin across cell types and contexts. Effect sizes and 95% CI were calculated using a logistic regression model.
(B) Comparison of effect sizes from the genome-wide peaks-based analysis and the accessible-wide ChromBPNet-based analysis.

**Supplementary Figure 10: ChromBPNet is correlated with constraint in rare variants.**
Correlation between sequence conservation (PhyloP) and ChromBPNet regulatory predictions across rare variants. Effect sizes and 95% CI are shown from a linear model estimating this correlation.

**Supplementary Figure 11: Fetal neurons shape genomic constraint.**
(A) Effect sizes and 95% CI from an analysis of regulatory effects (ChromBPNet scores) and evolutionary constraint (PhyloP), with each point representing a different cellular context colored by their broad context.
(B) Violin plots showing effect sizes stratified by broad context.

**Supplementary Figure 12: Development-specific predictions of an adult-specific variant.** This shows the full chromatin profile predictions and base-pair contribution scores for the adult-specific variant in Figure 3H in fetal excitatory neurons (top three panels) and adult excitatory neurons (bottom three panels). The top plots show chromatin profile predictions in a 300-bp window surrounding the variant with the predicted ATAC counts along the *y*-axis. Each colored line represents the base-pair resolution prediction for an input sequence containing the reference or alternate alleles. The bottom two plots show the output from DeepLIFT, which reflect contribution scores for each base in the sequence, along the *y*-axis.

**Supplementary Figure 13: Development-specific predictions of fetal-specific variants.** Figure panels (A) and (B) both show fetal-specific variants that are predicted to disrupt a "NEUROD1-ATOH_1" motif. In both panels, the top plots show chromatin profile predictions in a 300-bp window surrounding the variant with the predicted ATAC counts along the *y*-axis. Each colored line represents the base-pair resolution prediction for an input sequence containing the reference or alternate alleles. The bottom two plots show the output from DeepLIFT, which reflect contribution scores for each base in the sequence, along the *y*-axis.

**Supplementary Figure 14: SNP-SELEX-based TF binding disruption predictions.**
(A) Proportion of adult-specific or fetal-specific variants that disrupt different motifs according to motifbreakR predictions.
(B) Variants were predicted to disrupt binding of a TF if the deltaSVM score was greater than 1 (left, moderate variant effects) or greater than 2 (right, strong effect variants). Logistic regression was used to test for the association between fetal-specific versus adult-specific variants and TF disruption according to the specified thresholds. Log2 Enrichment values (*x*-axis) represent the $\log_2$ of the estimate (log-odds ratio) from this model.

**Supplementary Figure 15: FLARE model performance in ASD predictions.**
(A) Comparison of FLARE predictions across common variants, rare variants, and de novo variants. Each color represents a different FLARE model, and the *y*-axis displays model performance. "ASD" de novo, 1KG "rare", and 1KG "common" variant sets. $R^2$ reflects the prediction accuracy of comparing predicted PhyloP versus observed PhyloP scores.
(B) Mean FLARE predictions (*y*-axis) for 4 different FLARE-based models (baseline, heart, adult brain, fetal brain) in the 3 distinct variant sets. Error bars display ±1.96 standard errors for 95% CI.

**Supplementary Figure 16: FLARE prioritization of ASD mutations.**
(E) Values of de novo mutations near syndromic autism-associated genes for five scoring metrics, stratified by case and control status. "Early RG cbp" refers to the ChromBPNet score for early radial glia in the fetal brain.

### Supplementary Tables

**Supplementary Table 1:** Overview of the single-cell ATAC-seq dataset within this study. Includes information on cell types present, as well as number of common and rare variants that are accessible in each cell type and have a significant ChromBPNet prediction (cbp; empirical *P*-value < 0.01) in each cell type.

**Supplementary Table 2:** Information related to ChromBPNet model performance, shown for each of the 5 folds and the mean performance across the 5 folds.

**Supplementary Table 3:** Number of motif instances per peak based on 118 manually-annotated, ChromBPNet-derived motif instances for each adult and fetal brain cell type.

**Supplementary Table 4:** MotifbreakR results for cell-type-specific versus shared variants. Additional columns are the default output from motifbreakR.

**Supplementary Table 5:** Significance values for the ChromBPNet enrichment of fine-mapped eQTL variants.

**Supplementary Table 6:** Stratified linkage disequilibrium score regression analysis of the 9 tested GWAS (8 phenotypes). FDR values refer to nominal P values that were corrected individually for each phenotype.

**Supplementary Table 7:** Significance values for the ChromBPNet enrichment of fine-mapped GWAS variants. FDR calculated for each phenotype separately.

**Supplementary Table 8:** Input related to the Alzheimer's disease GSEA analysis. List of GWAS variants, two closest genes according to TSS, and whether they have evidence for microglia-specific effects.

**Supplementary Table 9:** Results from Alzheimer's disease GSEA analysis. "db"=gene set database, "Term"=gene set, "FDR_*"=FDR-corrected significance values for that term, "Genes_*"=list of genes, "GeneCount_*"=number of genes. Analysis performed for microglia-specific GWAS variants and their genes ("_microglia") and a cell-type-agnostic analysis ("_bulk").

**Supplementary Table 10:** Analyses related to cell-type-specific constraint. "phylop_vs_cbp" refers to correlating ChromBPNet scores with PhyloP constraint across rare variants. "rare_vs_common" refers to comparing the distribution of ChromBPNet scores between rare and common variants. "rare_vs_common.peaks" refers to comparing the proportion of accessible rare versus common variants. "rare_vs_common.diff.fetal_brain.Excitatory_neurons" refers to a sensitivity analysis to ensure that differences in ChromBPNet scores between rare and common variants were not confounded by cell-type-specific accessibility. This analysis restricted to variants accessible in both fetal brain excitatory neurons (our top-ranked cell type) and another cell type (where "numpeak" specifies the number of co-accessible variants tested). "est" and "se" refers to the estimated effect sizes from the linear model that were used to generate all relevant figures.

**Supplementary Table 11:** ChromBPNet-derived motif results for adult-specific versus fetal-specific variants for the adult and fetal excitatory neuron models.

**Supplementary Table 12:** MotifbreakR results for adult-specific versus fetal-specific variants. Additional columns are the default output from motifbreakR.

**Supplementary Table 13:** DeltaSVM TF binding predictions based on snp-selex for adult-specific versus fetal-specific variants.

**Supplementary Table 14:** Scores for ASD mutations near well-reported syndromic ASD genes for 5 metrics (left to right): FLARE-fb, FLARE-h, PhyloP, CADD, and fetal brain early radial glia.

### Supplementary Files

**Supplementary File 1:** Table with embedded TF motif logos for all ChromBPNet-derived clustered motif patterns used in this paper, along with their annotations and closest matches to known motifs in existing motif collections.

A

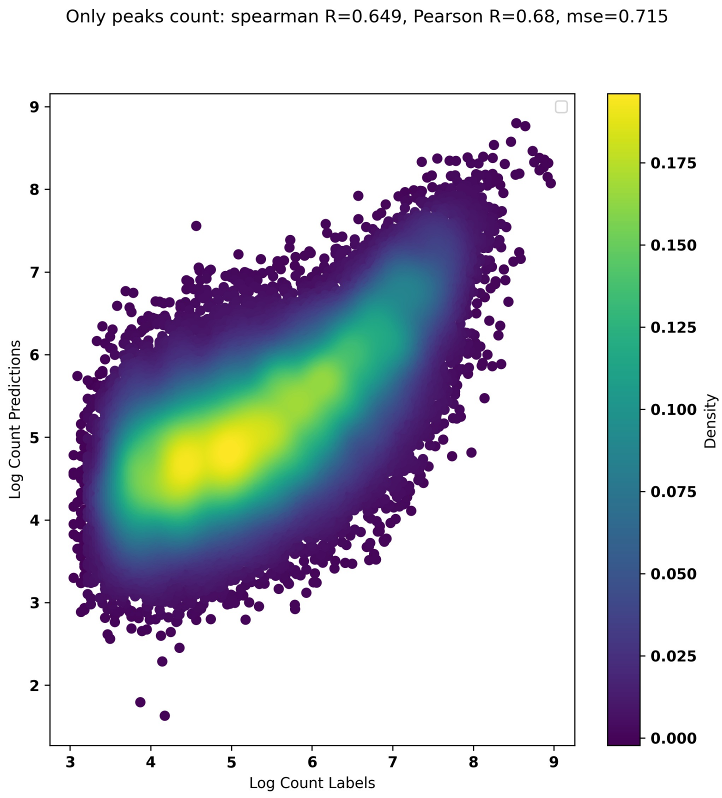


B

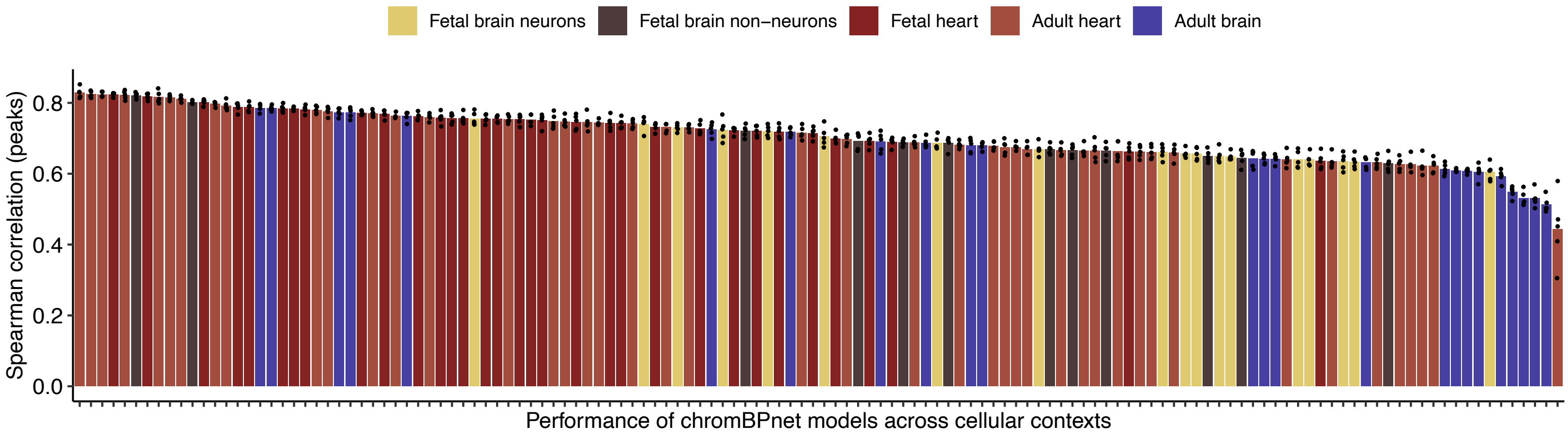


**Supplementary Figure 1: Dataset characteristics.**

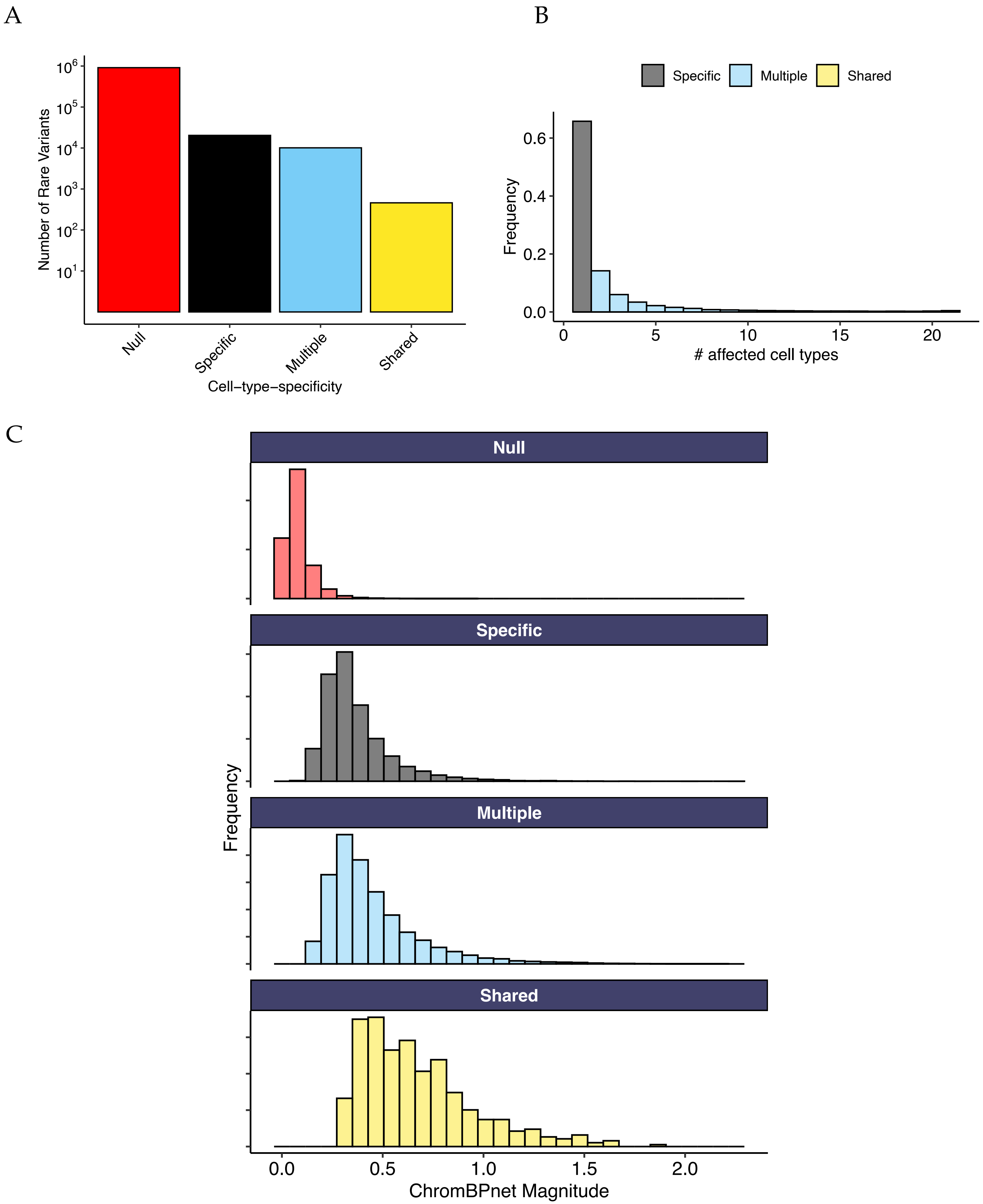


**Supplementary Figure 2: Analysis of cell-type-specificity**.

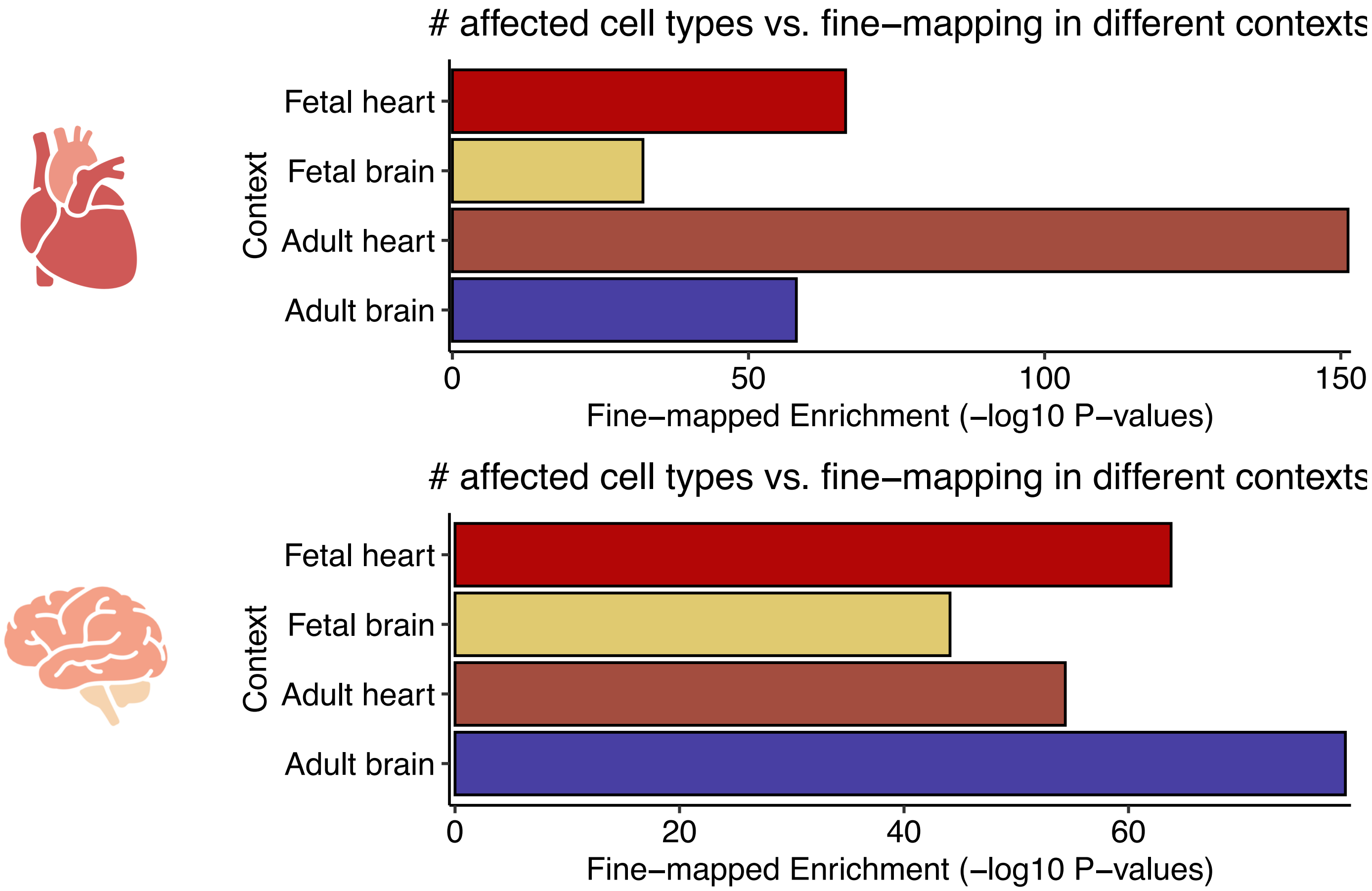


**Supplementary Figure 3: Fine-mapped eQTLs are associated with broad effects**.

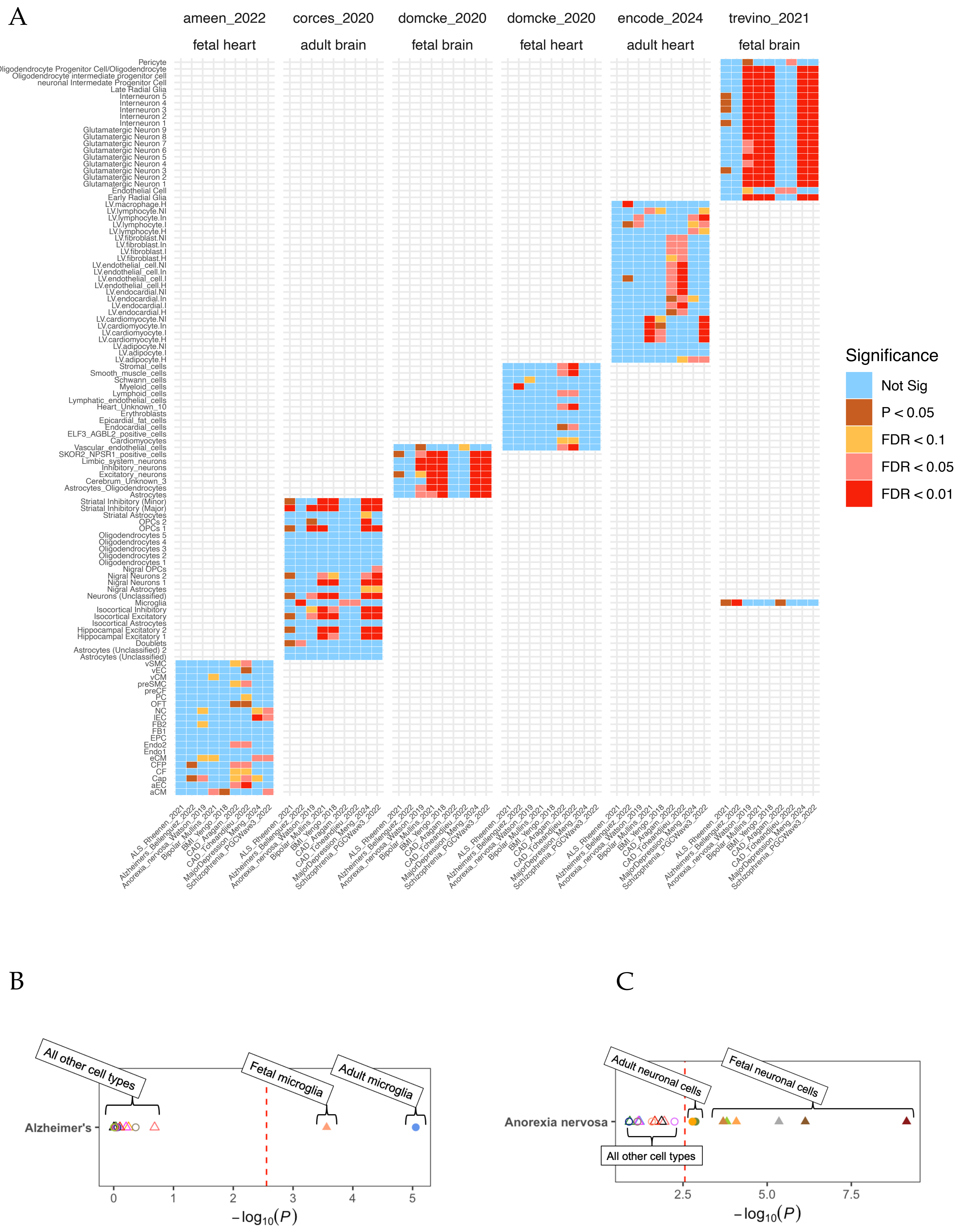


**Supplementary Figure 4: Cell-type-specific GWAS heritability enrichments.**

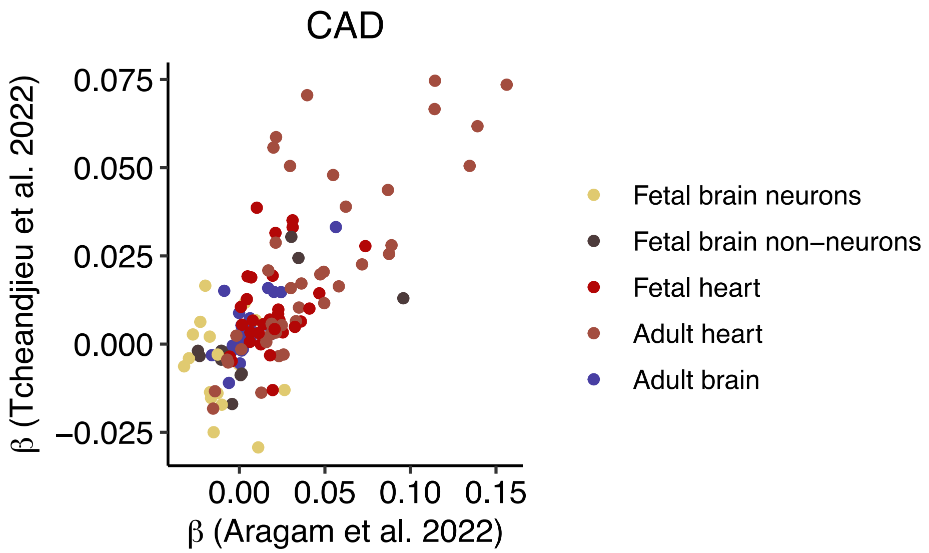


**Supplementary Figure 5: ChromBPnet enrichments in two CAD GWAS.**

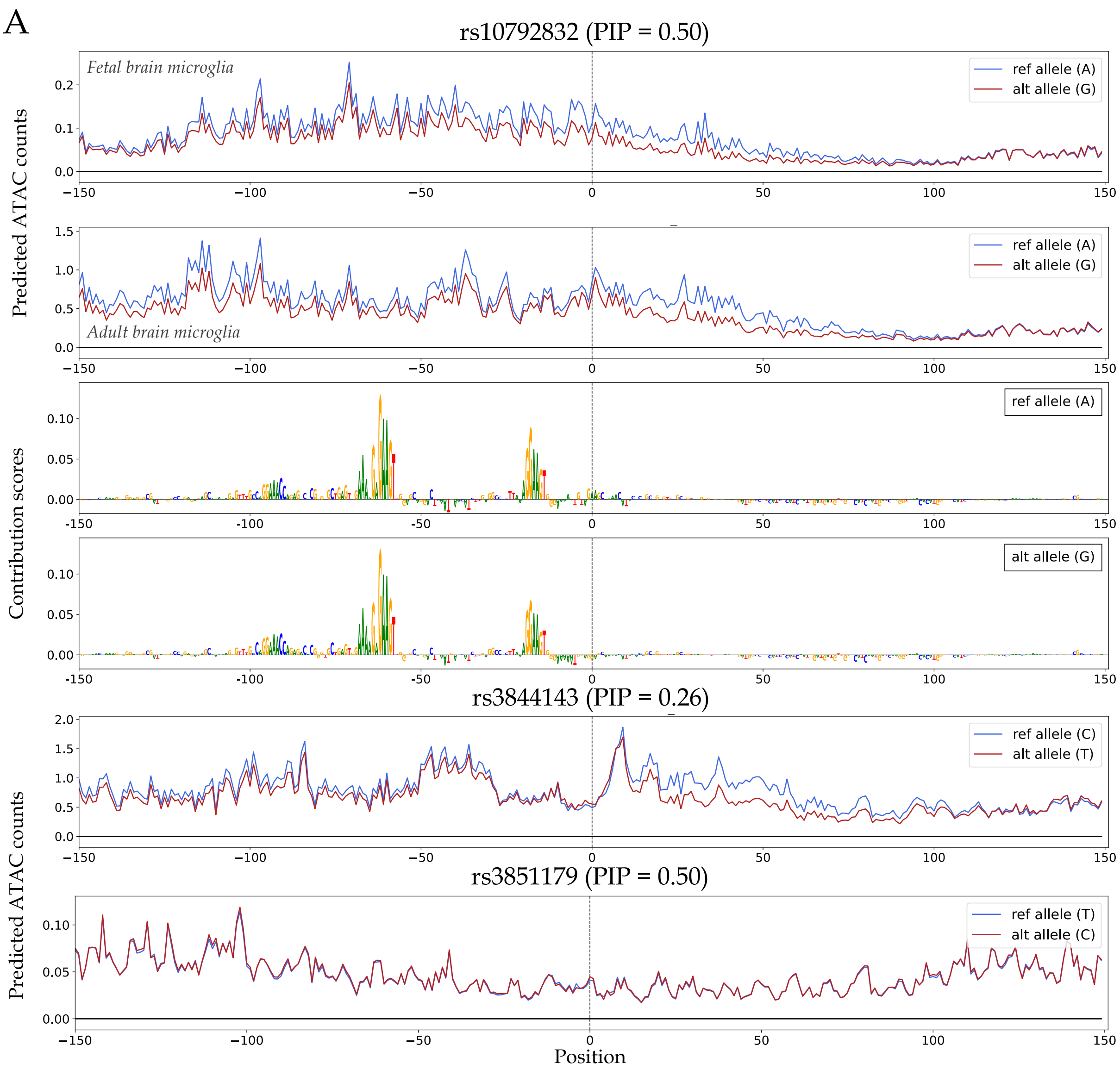


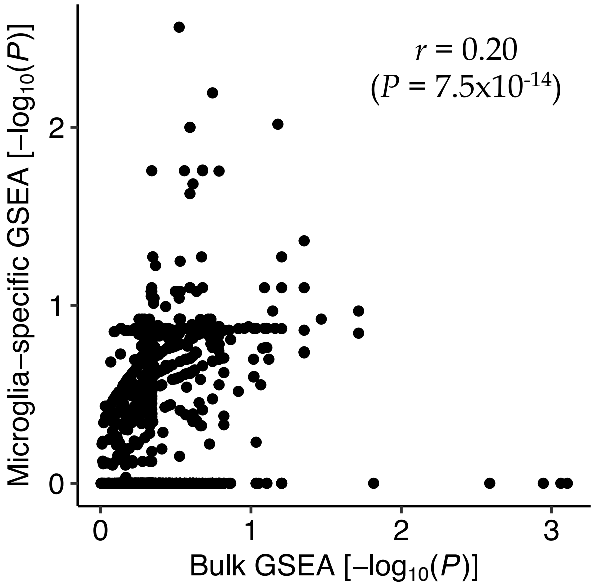


**Supplementary Figure 6: Microglia effects at Alzheimer's disease GWAS loci.**

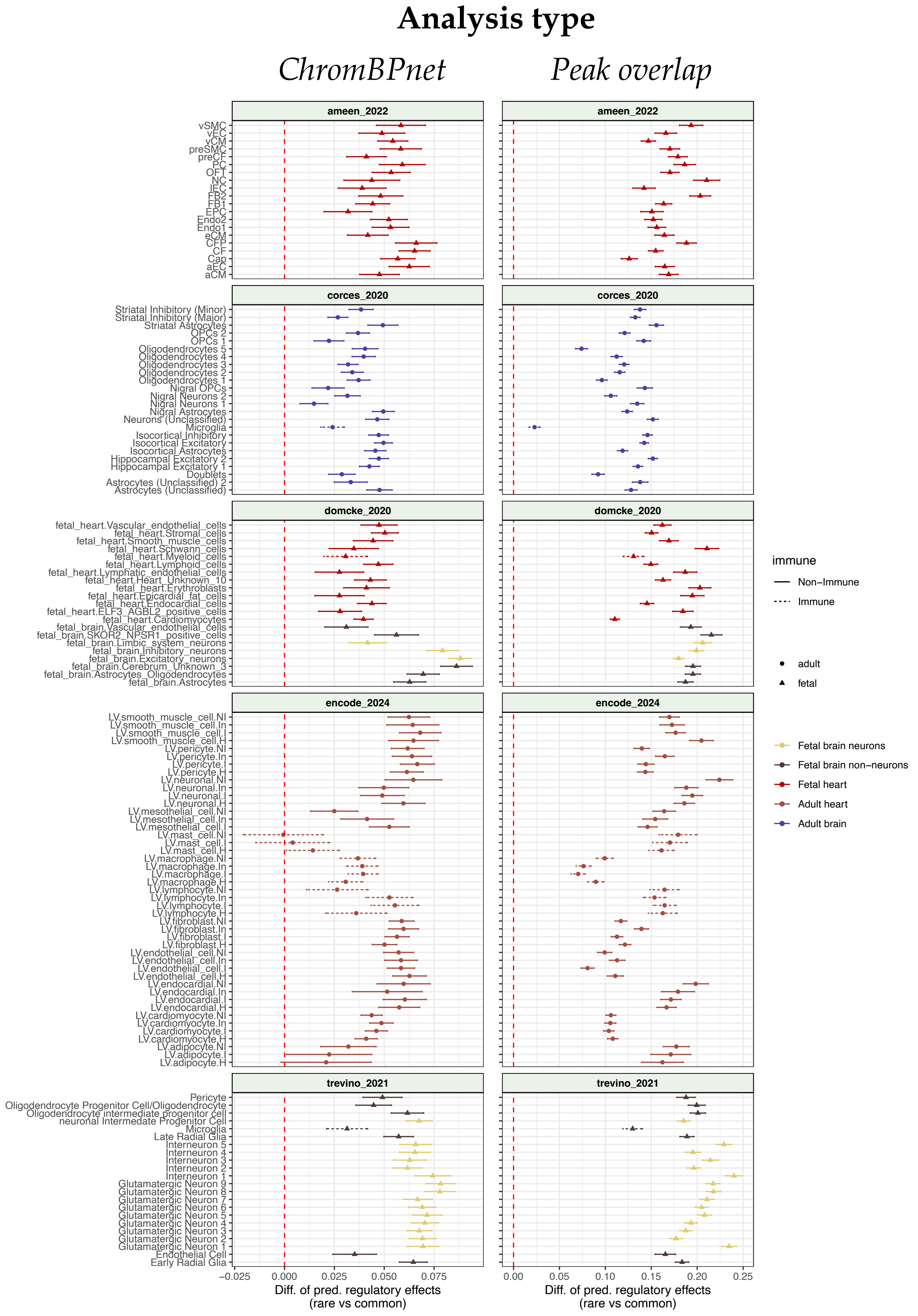


**Supplementary Figure 7: Common versus rare variant analyses.**

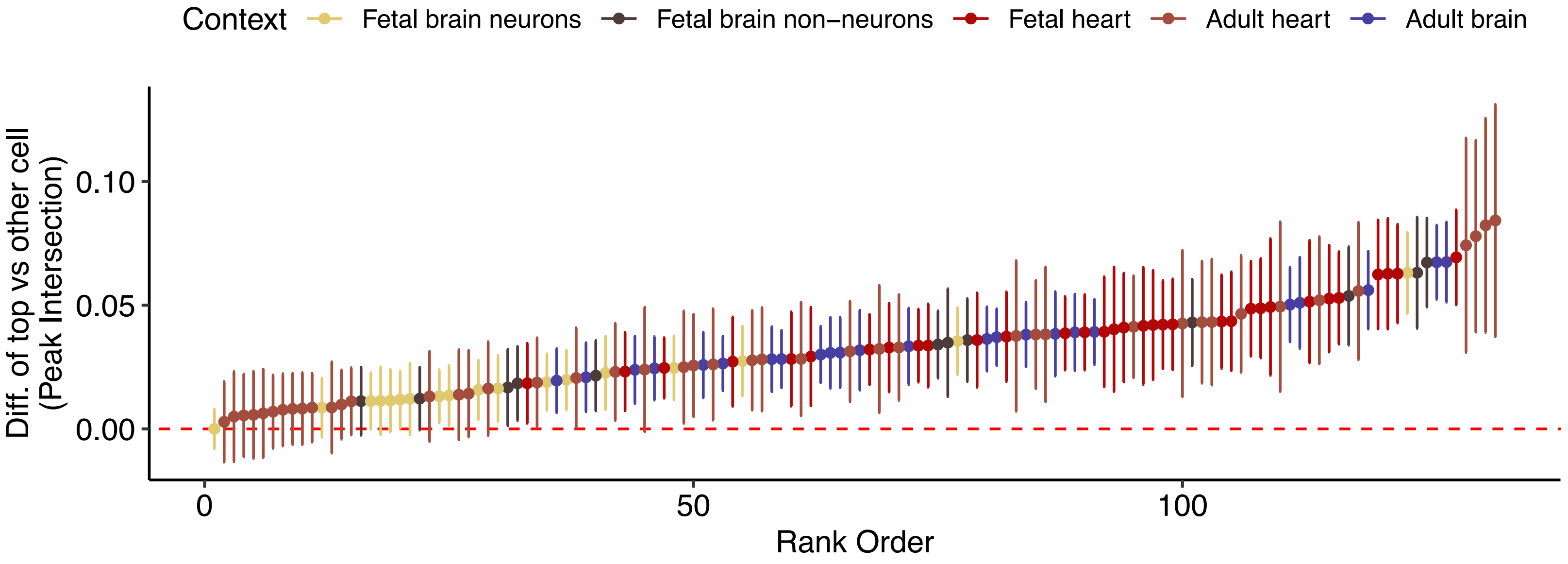


**Supplementary Figure 8: Restricting common versus rare analyses to identical variant sets.**

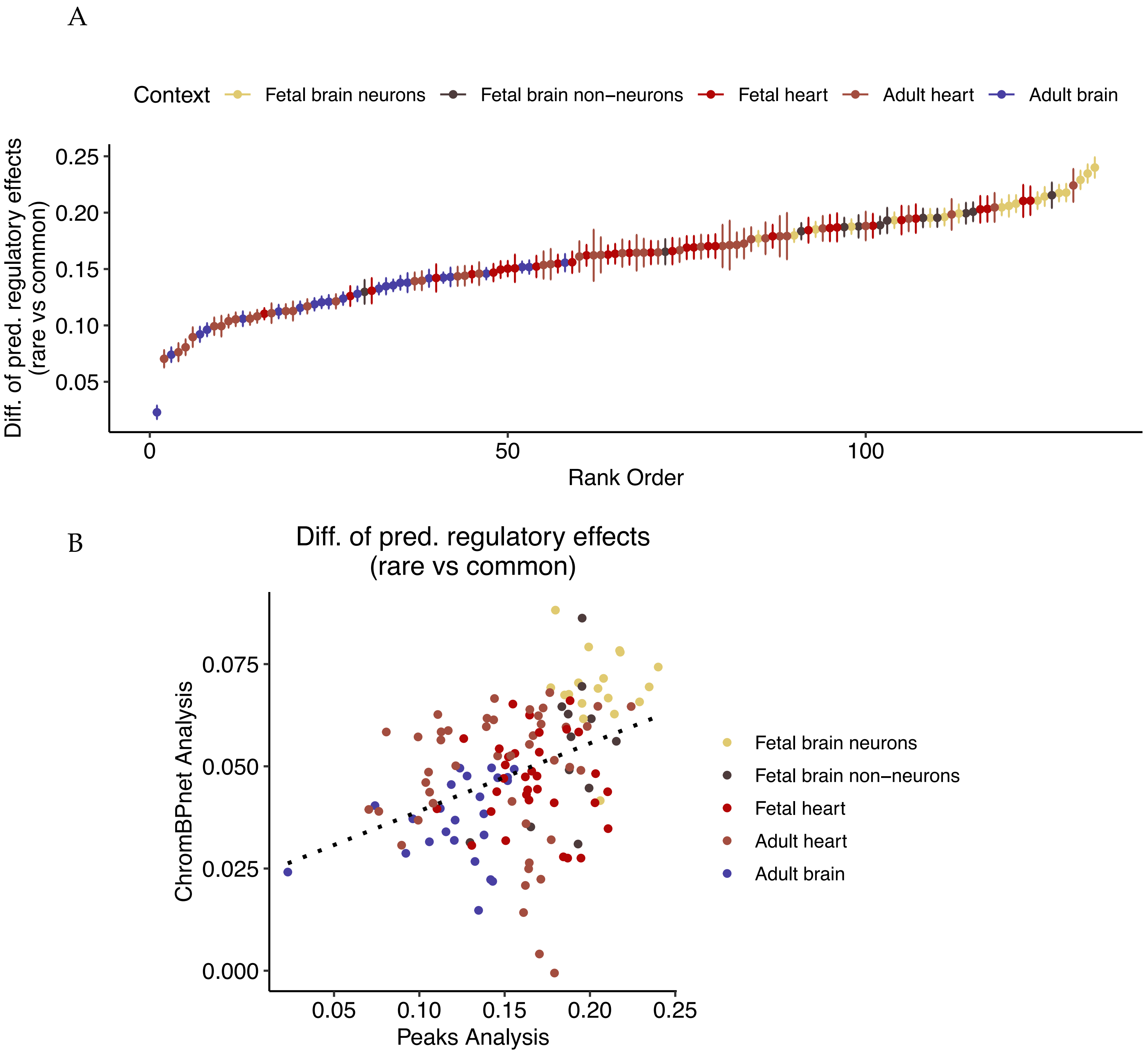


**Supplementary Figure 9: Proportions of accessible rare and common variants.**

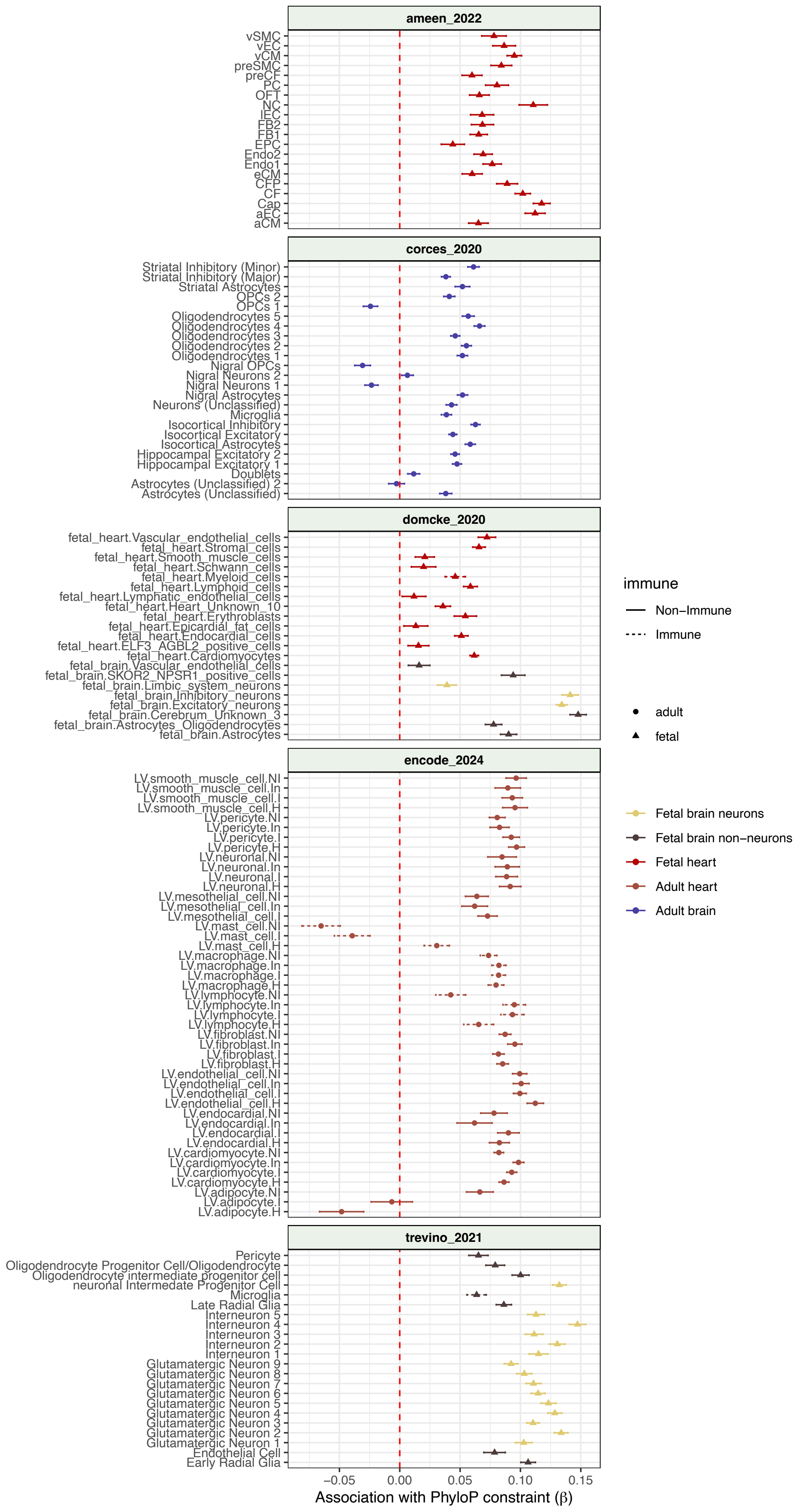


**Supplementary Figure 10: ChromBPnet is correlated with constraint in rare variants.**

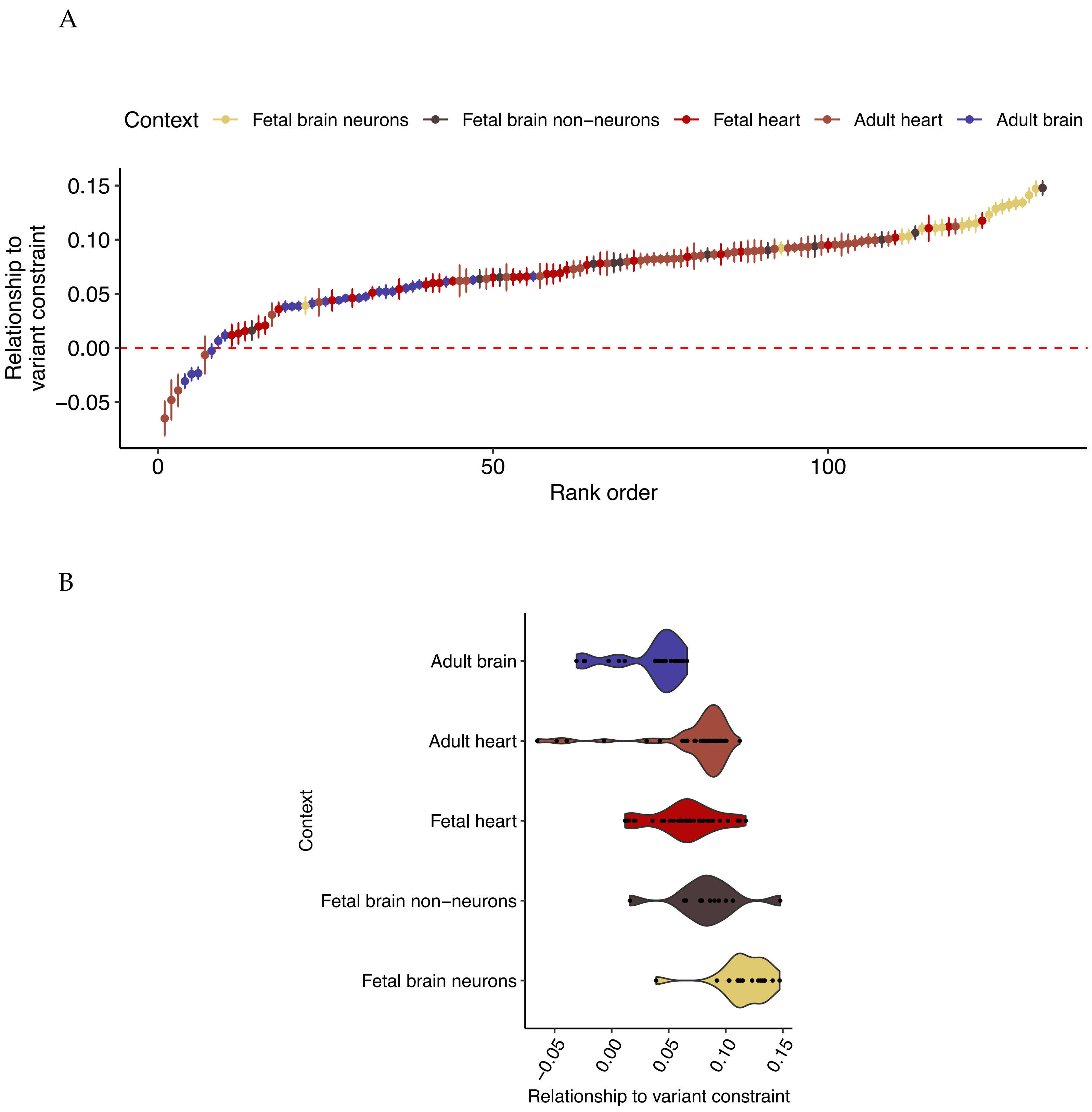


**Supplementary Figure 11: Fetal neurons shape genomic constraint.**

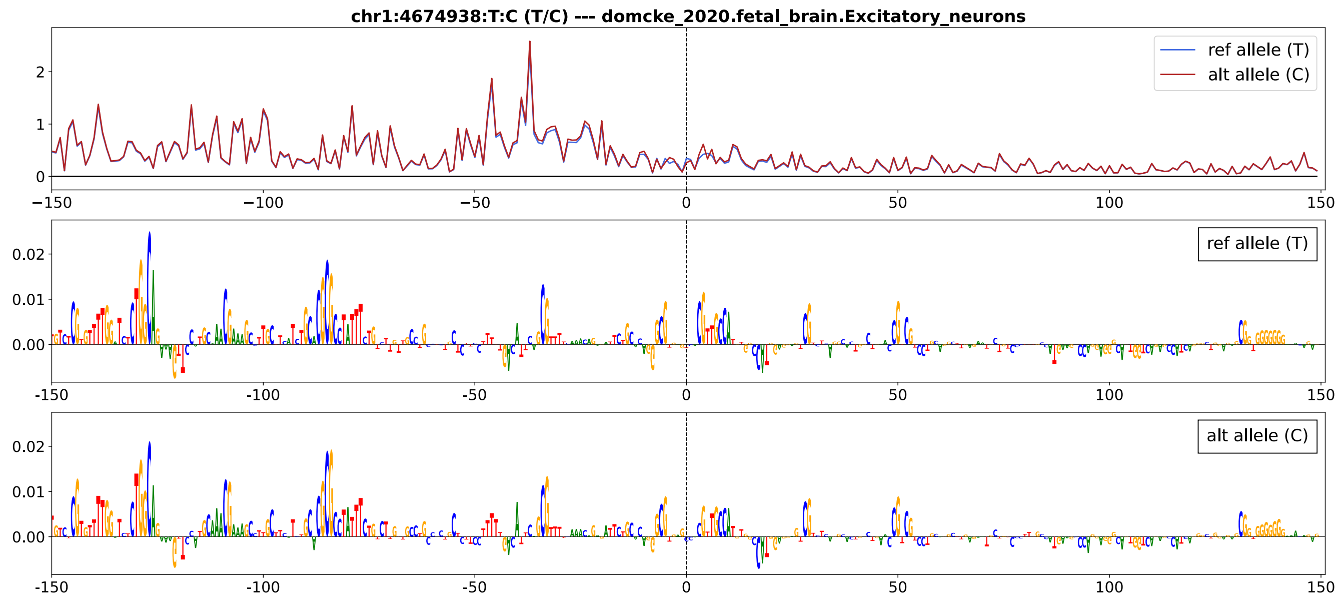


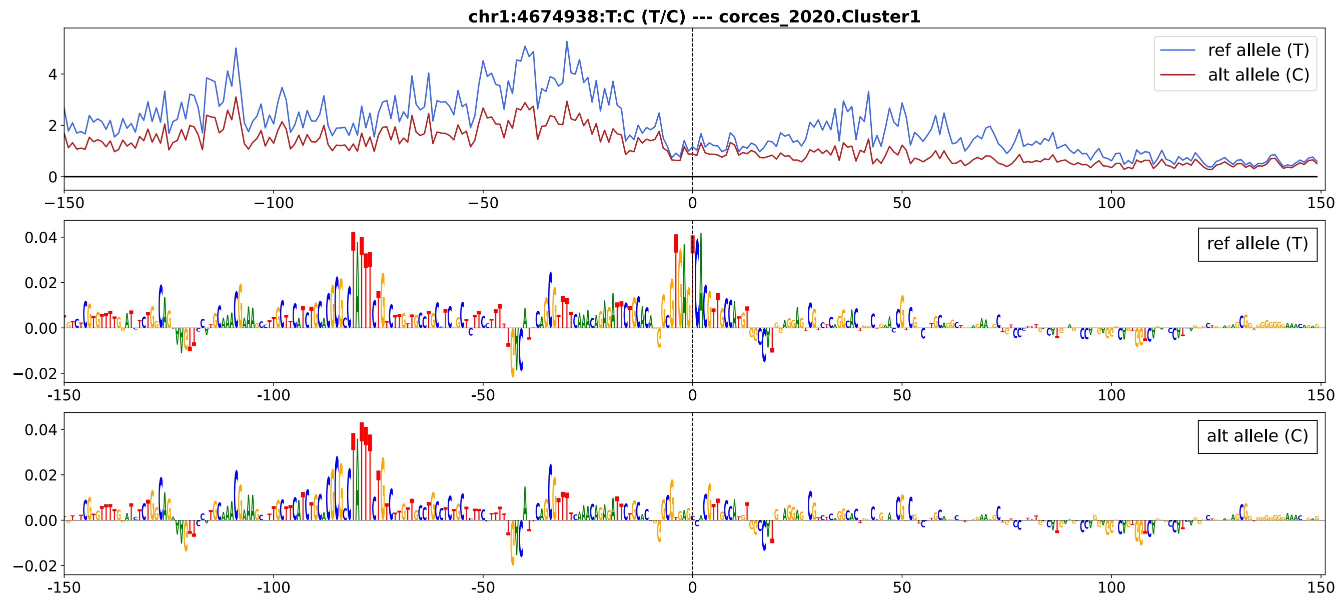


**Supplementary Figure 12: Development-specific predictions of an adult-specific variant.**

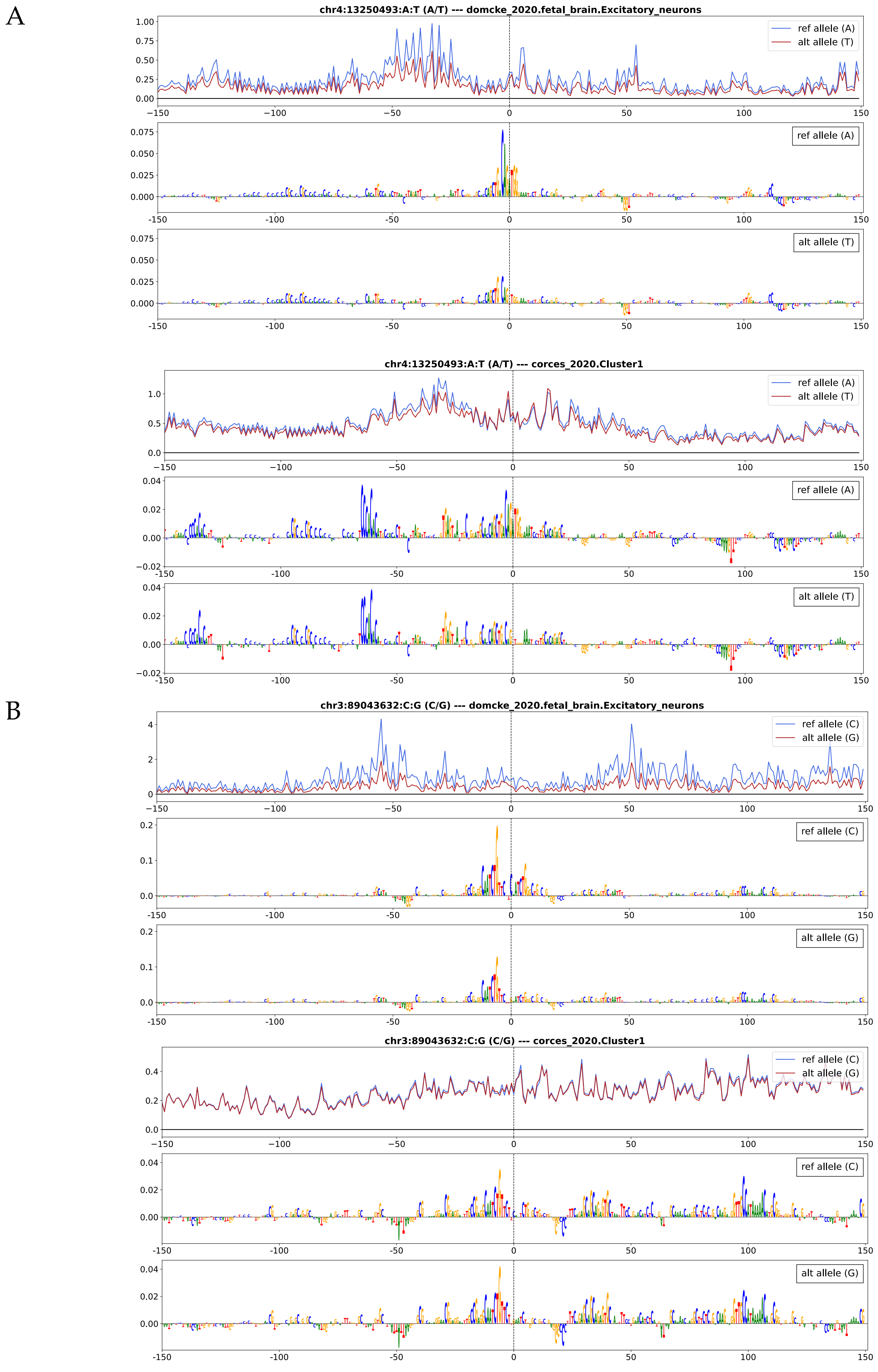


**Supplementary Figure 13: Development-specific predictions of fetal-specific variants.**

A

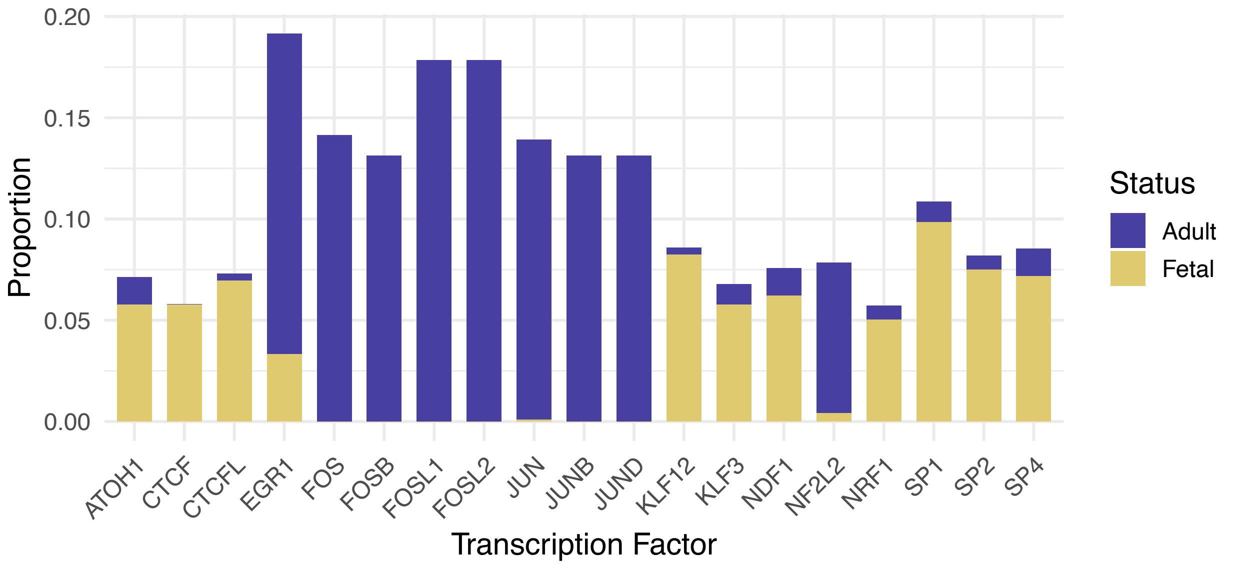


B

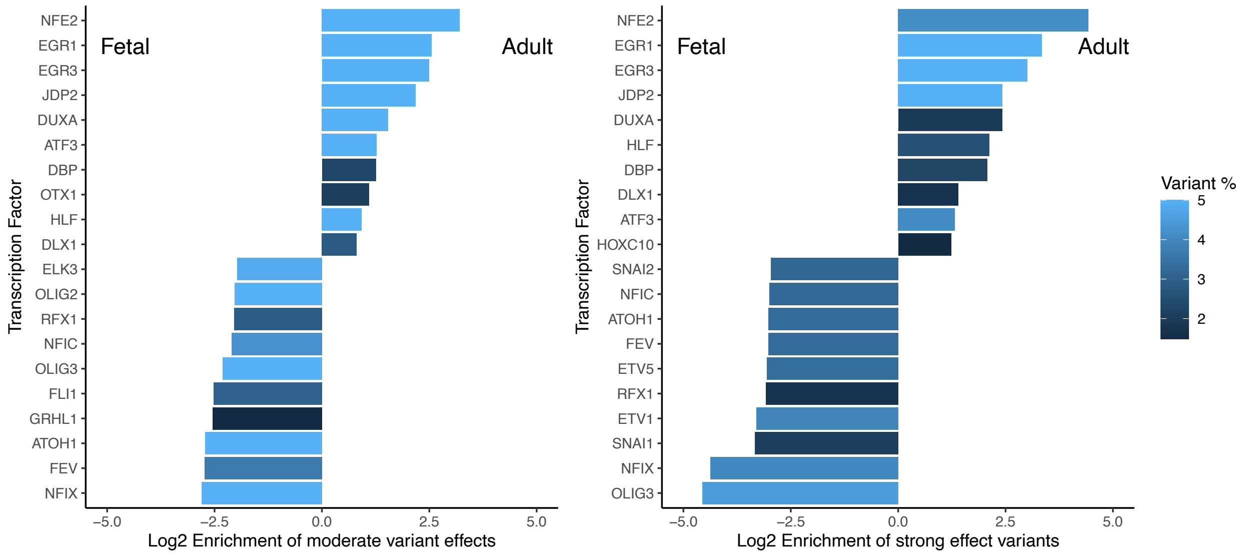


**Supplementary Figure 14: SNP-SELEX-based TF binding disruption predictions.**

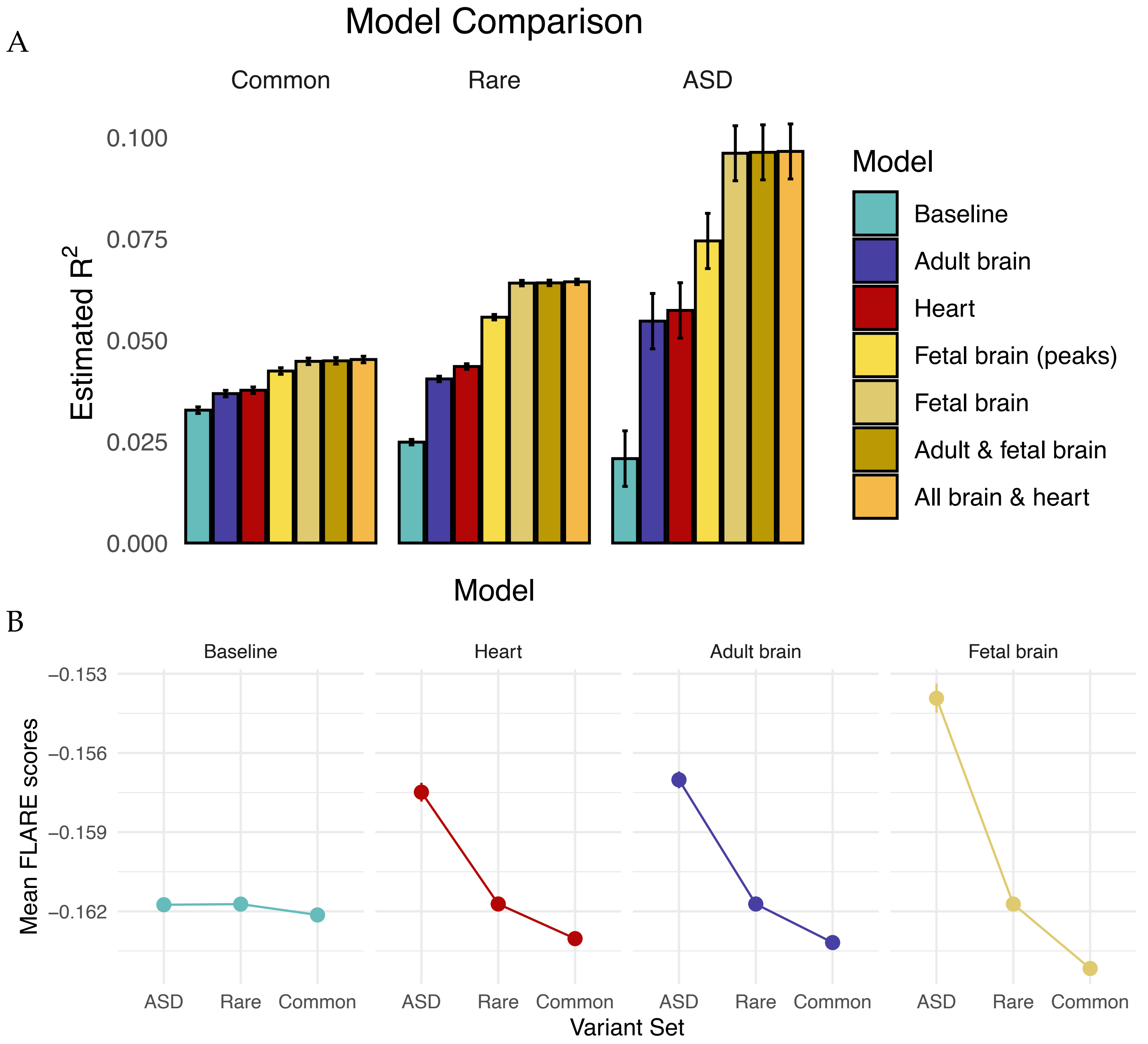


**Supplementary Figure 15: FLARE model performance in ASD predictions.**

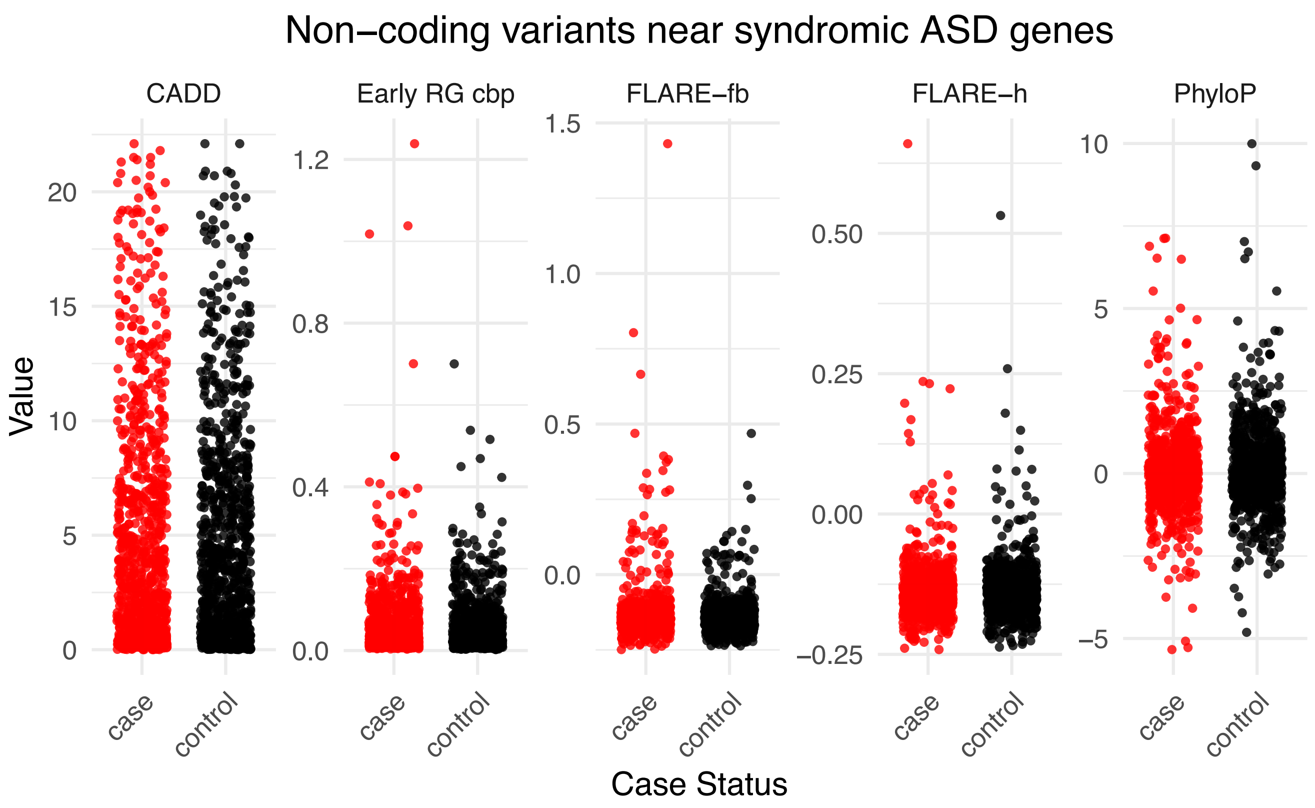


**Supplementary Figure 16: FLARE prioritization of ASD mutations.**

### References


1. Abell, N. S. *et al.* Multiple causal variants underlie genetic associations in humans. *Science* **375**, 1247–1254 (2022).
2. Wu, Z. *et al.* PICALM rs3851179 Variants Modulate Left Postcentral Cortex Thickness, CSF Amyloid β42, and Phosphorylated Tau in the Elderly. *Brain Sci.* **12**, 1681 (2022).
3. Wu, Z. *et al.* The Effects of PICALM rs3851179 and Age on Brain Atrophy and Cognition Along the Alzheimer's Disease Continuum. *Mol. Neurobiol.* **61**, 6984–6996 (2024).
4. Santos-Rebouças, C. B. *et al.* rs3851179 Polymorphism at 5' to the PICALM Gene is Associated with Alzheimer and Parkinson Diseases in Brazilian Population. *Neuromolecular Med.* **19**, 293–299 (2017).
5. Sun, D.-M. *et al.* Effect of PICALM rs3851179 polymorphism on the default mode network function in mild cognitive impairment. *Behav. Brain Res.* **331**, 225–232 (2017).
6. Liu, G. *et al.* Lack of association between PICALM rs3851179 polymorphism and Alzheimer's disease in Chinese population and APOEε4-negative subgroup. *Neurobiol. Aging* **34**, 1310.e9–10 (2013).
7. Domcke, S. *et al.* A human cell atlas of fetal chromatin accessibility. *Science* **370**, eaba7612 (2020).
8. Corces, M. R. *et al.* Single-cell epigenomic analyses implicate candidate causal variants at inherited risk loci for Alzheimer's and Parkinson's diseases. *Nat. Genet.* **52**, 1158–1168 (2020).
9. Trevino, A. E. *et al.* Chromatin and gene-regulatory dynamics of the developing human cerebral cortex at single-cell resolution. *Cell* **184**, 5053-5069.e23 (2021).
10. Duclot, F. & Kabbaj, M. The Role of Early Growth Response 1 (EGR1) in Brain Plasticity and Neuropsychiatric Disorders. *Front. Behav. Neurosci.* **11**, 35 (2017).
11. Maddox, S. A., Monsey, M. S. & Schafe, G. E. Early growth response gene 1 (Egr-1) is required for new and reactivated fear memories in the lateral amygdala. *Learn. Mem. Cold Spring Harb. N* **18**, 24–38 (2011).
12. Lee, D. *et al.* A method to predict the impact of regulatory variants from DNA sequence. *Nat. Genet.* **47**, 955–961 (2015).
13. Li, M. *et al.* Integrative functional genomic analysis of human brain development and neuropsychiatric risks. *Science* **362**, eaat7615 (2018).
14. GTEx Consortium. The GTEx Consortium atlas of genetic regulatory effects across human tissues. *Science* **369**, 1318–1330 (2020).
15. Loyd, M. R., Okamoto, Y., Randall, M. S. & Ney, P. A. Role of AP1/NFE2 binding sites in endogenous alpha-globin gene transcription. *Blood* **102**, 4223–4228 (2003).
16. An, J.-Y. *et al.* Genome-wide de novo risk score implicates promoter variation in autism spectrum disorder. *Science* **362**, eaat6576 (2018).
