## Supplementary File 1 for "Mapping the regulatory effects of common and rare non-coding variants across cellular and developmental contexts in the brain and heart"

| motif_name | forward_logo | reverse_logo | vierstra_similarity | vierstra_match | vierstra_logo | hocomoco_similarity | hocomoco_match | hocomoco_logo | num_patterns | num_seqlets | num_samples | num_datasets | datasets |
|---|---|---|---|---|---|---|---|---|---|---|---|---|---|
| CTCF_1 | | | 0.9834836704 | CTCFL_M09507_2.00 | | 0.9733986486 | CTCF.H12CORE.0.P.B | | 54 | 947567 | 54 | 3 | corces_2020,domcke_2020,trevino_2021 |
| NFI_1 | | | 0.9643396574 | NFIC_M09636_2.00 | | 0.9043977752 | NFIX.H12CORE.0.SM.B | | 47 | 537160 | 47 | 3 | corces_2020,domcke_2020,trevino_2021 |
| SOX::SOX_1 | | | 0.963069327 | SOX10_M09387_2.00 | | 0.9800090371 | SOX8.H12CORE.0.PSM.A | | 36 | 450524 | 23 | 3 | corces_2020,domcke_2020,trevino_2021 |
| NDF-ATOH_1 | | | 0.9550309388 | MSGN1_MA1524.1 | | 0.9598438967 | NGN2.H12CORE.0.P.B | | 16 | 444501 | 16 | 3 | corces_2020,domcke_2020,trevino_2021 |
| SP-KLF_1 | | | 0.9446619468 | SP4_M08296_2.00 | | 0.9699253433 | SP4.H12CORE.0.P.C | | 58 | 372948 | 54 | 3 | corces_2020,domcke_2020,trevino_2021 |
| NFY | | | 0.9848404927 | NFYA_MA0060.1 | | 0.9787944488 | NFYA.H12CORE.0.P.B | | 54 | 317250 | 54 | 3 | corces_2020,domcke_2020,trevino_2021 |
| BHLH_1 | | | 0.9776998739 | MYOG_MA0500.2 | | 0.9554164442 | MYF5.H12CORE.0.P.B | | 21 | 256922 | 20 | 3 | corces_2020,domcke_2020,trevino_2021 |
| LHX_1 | | | 0.9900286651 | VSX1_M03087_2.00 | | 0.9571050543 | RX.H12CORE.0.SM.B | | 23 | 241042 | 23 | 3 | corces_2020,domcke_2020,trevino_2021 |
| RFX_1 | | | 0.9718724672 | RFX3_M09627_2.00 | | 0.9791723332 | RFX1.H12CORE.1.PSM.A | | 50 | 201919 | 50 | 3 | corces_2020,domcke_2020,trevino_2021 |
| NRF1 | | | 0.9512382496 | NRF1_M09443_2.00 | | 0.9144143628 | NRF1.H12CORE.0.PS.A | | 74 | 194827 | 53 | 3 | corces_2020,domcke_2020,trevino_2021 |
| ETS_1 | | | 0.9867246044 | ELF5_M02989_2.00 | | 0.982897238 | ETS2.H12CORE.1.P.B | | 7 | 191381 | 7 | 3 | corces_2020,domcke_2020,trevino_2021 |
| AP1 | | | 0.9937943585 | BNC2_MA1928.1 | | 0.9816644445 | JUNB.H12CORE.0.PM.A | | 26 | 184212 | 24 | 2 | corces_2020,trevino_2021 |
| EGR_1 | | | 0.9832522527 | EGR1_M04438_2.00 | | 0.9099562236 | WT1.H12CORE.1.P.B | | 6 | 182388 | 6 | 1 | corces_2020 |
| ETS_2 | | | 0.9561441054 | ELK3_M04730_2.00 | | 0.9541554717 | ELK4.H12CORE.0.PSM.A | | 53 | 163362 | 52 | 3 | corces_2020,domcke_2020,trevino_2021 |
| ATF_1 | | | 0.9737564435 | CREB1_M04258_2.00 | | 0.9371748248 | GMEB2.H12CORE.2.SM.B | | 55 | 137965 | 53 | 3 | corces_2020,domcke_2020,trevino_2021 |
| MEF2_1 | | | 0.981156337 | MEF2A_M10936_2.00 | | 0.979775856 | MEF2A.H12CORE.0.P.B | | 13 | 116631 | 13 | 3 | corces_2020,domcke_2020,trevino_2021 |
| ZNF143_1 | | | 0.9715007649 | ETS1_M07950_2.00 | | 0.9812102779 | ZN143.H12CORE.0.P.B | | 54 | 92698 | 54 | 3 | corces_2020,domcke_2020,trevino_2021 |
| POU_1 | | | 0.9658141536 | POU3F2_M05476_2.00 | | 0.9271605755 | VENTX.H12CORE.1.S.C | | 23 | 78266 | 23 | 2 | domcke_2020,trevino_2021 |
| NDF-ATOH_2 | | | 0.8557861087 | TBX21_M09425_2.00 | | 0.8875742787 | LYL1.H12CORE.0.P.C | | 8 | 67776 | 8 | 2 | domcke_2020,trevino_2021 |
| ZEB-SNAI | | | 0.9734562349 | ZEB1_M10093_2.00 | | 0.9590286457 | ZEB2.H12CORE.0.P.B | | 45 | 64734 | 42 | 3 | corces_2020,domcke_2020,trevino_2021 |
| RFX_2 | | | 0.9513063933 | RFX4_M03457_2.00 | | 0.9125727529 | RFX6.H12CORE.0.P.C | | 41 | 50986 | 41 | 3 | corces_2020,domcke_2020,trevino_2021 |
| SOX::SOX_2 | | | 0.8604444692 | STAT4_M11361_2.00 | | 0.8519817234 | SOX1.H12CORE.1.S.B | | 16 | 33916 | 10 | 2 | corces_2020,trevino_2021 |
| YY1 | | | 0.9385812511 | YY1_M07859_2.00 | | 0.9281498783 | TYY1.H12CORE.0.PSM.A | | 43 | 33144 | 43 | 3 | corces_2020,domcke_2020,trevino_2021 |

| | | | | | | | | | |
|---|---|---|---|---|---|---|---|---|---|
| NR3C | 0.9571662316 | NR3C1_M07987_2.00 | 0.9746475904 | ANDR.H12CORE.0.P.B | 9 | 33137 | 9 | 1 | corces_2020 |
| ARNT-USF | 0.9843822077 | BHLHE41_M02783_2.00 | 0.9781481898 | BHE41.H12CORE.0.PSM.A | 29 | 32724 | 28 | 3 | corces_2020,domcke_2020,trevino_2021 |
| TEF | 0.9842370534 | TEF_M02850_2.00 | 0.9727042024 | HLF.H12CORE.0.P.B | 8 | 30575 | 8 | 1 | corces_2020 |
| POU2F_1 | 0.9615130258 | POU2F1_M03309_2.00 | 0.9528447686 | PO3F2.H12CORE.0.P.B | 19 | 25296 | 19 | 2 | domcke_2020,trevino_2021 |
| MEIS | 0.9692218315 | MEIS1_M10709_2.00 | 0.9749670787 | PBX1.H12CORE.2.P.B | 2 | 23861 | 2 | 2 | corces_2020,trevino_2021 |
| ETS::ATF_1 | 0.8837521355 | ELK1::SREBF2_MA1933.1 | 0.7846750919 | FEV.H12CORE.0.S.B | 48 | 23172 | 47 | 3 | corces_2020,domcke_2020,trevino_2021 |
| POU_2 | 0.8505824037 | POU5F1_M07976_2.00 | 0.8389827425 | PO5F1.H12CORE.0.P.B | 10 | 20528 | 10 | 3 | corces_2020,domcke_2020,trevino_2021 |
| POU_3 | 0.9465921188 | POU1F1_M03300_2.00 | 0.8951716214 | PO4F1.H12CORE.0.S.B | 12 | 19367 | 12 | 2 | domcke_2020,trevino_2021 |
| ATF_2 | 0.987964615 | JDP2_M04288_2.00 | 0.9782677744 | ATF2.H12CORE.0.PSM.A | 23 | 18792 | 23 | 3 | corces_2020,domcke_2020,trevino_2021 |
| CEBP_1 | 0.9908555597 | CEBPB_M09993_2.00 | 0.9847266396 | CEBPE.H12CORE.0.P.B | 2 | 14708 | 2 | 2 | corces_2020,trevino_2021 |
| IRF_1 | 0.9661781502 | IRF4_M09235_2.00 | 0.895133356 | IRF4.H12CORE.0.P.B | 2 | 14350 | 2 | 2 | corces_2020,trevino_2021 |
| SOX::SOX_3 | 0.8962454688 | SOX10_M09387_2.00 | 0.907772278 | SOX8.H12CORE.0.PSM.A | 3 | 13988 | 3 | 2 | corces_2020,trevino_2021 |
| TBR1_1 | 0.9537955363 | TBR1_M03560_2.00 | 0.965285132 | TBXT.H12CORE.1.PSM.A | 5 | 10212 | 5 | 2 | corces_2020,trevino_2021 |
| ZNF143_2 | 0.9104704746 | ZNF76_M08278_2.00 | 0.8457986004 | ZNF76.H12CORE.0.P.B | 37 | 7583 | 37 | 3 | corces_2020,domcke_2020,trevino_2021 |
| IRF_2 | 0.9340694962 | STAT1_M08230_2.00 | 0.951945464 | IRF3.H12CORE.0.PS.A | 2 | 6758 | 2 | 2 | corces_2020,trevino_2021 |
| NFI-half_2 | 0.8604641614 | TFAP2A_M09755_2.00 | 0.8418682028 | ZBT43.H12CORE.0.S.C | 1 | 6538 | 1 | 1 | corces_2020 |
| FOX_1 | 0.957547766 | FOXF1_M10530_2.00 | 0.9634924287 | FOXL2.H12CORE.0.PSM.A | 2 | 5612 | 2 | 1 | trevino_2021 |
| POU::POU | 0.8769838109 | POU3F2_M05474_2.00 | 0.8096528871 | PIT1.H12CORE.1.S.B | 5 | 5262 | 5 | 2 | domcke_2020,trevino_2021 |
| BHLH_2 | 0.88469737 | TFAP4_M09803_2.00 | 0.8211317597 | MYF5.H12CORE.0.P.B | 2 | 5014 | 2 | 1 | corces_2020 |
| NFI_2 | 0.8276952953 | NFIX_M03477_2.00 | 0.8088602341 | NFIA.H12CORE.0.P.B | 2 | 4423 | 2 | 1 | trevino_2021 |
| ETS::ETS_1 | 0.8807326624 | ETV6_M02990_2.00 | 0.8097556034 | ELK4.H12CORE.0.PSM.A | 15 | 4089 | 15 | 3 | corces_2020,domcke_2020,trevino_2021 |
| MEF2_2 | 0.9264550022 | MEF2A_M08212_2.00 | 0.8663752804 | MEF2D.H12CORE.0.PS.A | 1 | 4017 | 1 | 1 | corces_2020 |
| POU3F_1 | 0.9423534615 | POU3F4_M03318_2.00 | 0.8848224304 | PO3F3.H12CORE.1.P.C | 4 | 3960 | 4 | 1 | trevino_2021 |

| | | | | | | | | | | | | | |
|---|---|---|---|---|---|---|---|---|---|---|---|---|---|
| HLF | | | 0.972250562 | TEF_M04325_2.00 | | 0.9339382789 | CEBPB.H12CORE.1.SM.B | | 6 | 3844 | 6 | 2 | corces_2020,trevino_2021 |
| LHX::LHX_1 | | | 0.9236206335 | LHX2_M03107_2.00 | | 0.8999929688 | DLX2.H12CORE.1.S.B | | 4 | 3805 | 4 | 3 | corces_2020,domcke_2020,trevino_2021 |
| RUNX | | | 0.98231525 | RUNX3_M09369_2.00 | | 0.9690819572 | RUNX2.H12CORE.0.P.B | | 1 | 3789 | 1 | 1 | corces_2020 |
| HESX1 | | | 0.9023478027 | VSX2_M05039_2.00 | | 0.9159162018 | PO2F3.H12CORE.0.PS.A | | 5 | 3543 | 5 | 1 | trevino_2021 |
| SOX::SOX_4 | | | 0.8580829734 | SOX4_M05765_2.00 | | 0.8692345563 | SOX6.H12CORE.0.P.B | | 5 | 3406 | 5 | 1 | corces_2020 |
| SOX::SOX_5 | | | 0.8726257507 | SOX4_M05765_2.00 | | 0.8783968399 | SOX6.H12CORE.0.P.B | | 5 | 3378 | 5 | 1 | corces_2020 |
| ETV_1 | | | 0.9485618896 | ETV4_M04798_2.00 | | 0.8934272019 | ETV6.H12CORE.1.P.B | | 1 | 3293 | 1 | 1 | corces_2020 |
| BHLH::NFI_1 | | | 0.8535547905 | NFIX_M03476_2.00 | | 0.8617343833 | NFIB.H12CORE.1.PS.A | | 4 | 3195 | 4 | 2 | corces_2020,trevino_2021 |
| ROR | | | 0.9497406931 | RORC_M05880_2.00 | | 0.9765492024 | RORG.H12CORE.1.PS.A | | 3 | 2951 | 3 | 1 | corces_2020 |
| FOX_2 | | | 0.9497571422 | FOXC2_M03037_2.00 | | 0.9337876507 | FOXL1.H12CORE.0.S.B | | 2 | 2925 | 2 | 1 | trevino_2021 |
| SOX::SOX_6 | | | 0.7196055937 | SOX8_M05751_2.00 | | 0.6449560493 | SOX8.H12CORE.0.PSM.A | | 1 | 2827 | 1 | 1 | corces_2020 |
| EBF | | | 0.9631674552 | EBF1_M02963_2.00 | | 0.9823024431 | COE2.H12CORE.0.P.B | | 1 | 2690 | 1 | 1 | trevino_2021 |
| BHLH::NFI_2 | | | 0.8880794307 | MYOD1_M04121_2.00 | | 0.8706359207 | NFIA.H12CORE.0.P.B | | 2 | 2638 | 2 | 2 | corces_2020,trevino_2021 |
| SMAD | | | 0.901943819 | SMAD4_M09377_2.00 | | 0.9062370025 | SMAD3.H12CORE.2.P.C | | 2 | 2587 | 2 | 2 | domcke_2020,trevino_2021 |
| SOX::POU_1 | | | 0.8115249825 | POU1F1_M03300_2.00 | | 0.7999800074 | PO3F3.H12CORE.1.P.C | | 3 | 2466 | 3 | 1 | trevino_2021 |
| CEBP_2 | | | 0.9577783965 | CEBPA_M08813_2.00 | | 0.9472966035 | CEBPD.H12CORE.0.P.B | | 1 | 2386 | 1 | 1 | corces_2020 |
| BHLH::NFI_3 | | | 0.8914635279 | NFIX_M03477_2.00 | | 0.888559199 | NFIB.H12CORE.1.PS.A | | 2 | 2362 | 2 | 1 | trevino_2021 |
| SOX::SOX_7 | | | 0.6996468133 | SOX10_M09387_2.00 | | 0.7059262382 | BARH2.H12CORE.1.S.B | | 1 | 2293 | 1 | 1 | domcke_2020 |
| IRF_3 | | | 0.9151502457 | IRF2_M05546_2.00 | | 0.9072446794 | IRF3.H12CORE.0.PS.A | | 2 | 2180 | 2 | 2 | corces_2020,trevino_2021 |
| SOX::SOX_8 | | | 0.8117139023 | SOX10_M09387_2.00 | | 0.7564979998 | SOX8.H12CORE.0.PSM.A | | 1 | 2160 | 1 | 1 | corces_2020 |
| LHX::LHX_2 | | | 0.9091072478 | POU3F1_M05483_2.00 | | 0.8676300561 | MSX2.H12CORE.1.SM.B | | 5 | 2123 | 5 | 2 | domcke_2020,trevino_2021 |
| EGR::NDF-ATOH | | | 0.8776130602 | EGR1_M04438_2.00 | | 0.7986884797 | WT1.H12CORE.1.P.B | | 1 | 2104 | 1 | 1 | corces_2020 |
| NFY-repressive | | | 0.9702508209 | NFYA_MA0060.1 | | 0.9648587583 | NFYC.H12CORE.0.P.B | | 13 | 2060 | 13 | 2 | corces_2020,trevino_2021 |

| | | | | | | | | | | | | | |
|---|---|---|---|---|---|---|---|---|---|---|---|---|---|
| BHLH::BHLH | | | 0.7812442095 | MYOG_MA0500.2 | | 0.7454009905 | MYF5.H12CORE.0.P.B | | 1 | 1962 | 1 | 1 | corces_2020 |
| SOX::SOX_9 | | | 0.7947688002 | SOX10_M09387_2.00 | | 0.731986914 | SOX8.H12CORE.0.PSM.A | | 1 | 1735 | 1 | 1 | corces_2020 |
| RFX_3 | | | 0.8451895933 | RFX5_M09364_2.00 | | 0.8496568676 | RFX3.H12CORE.0.PSM.A | | 3 | 1551 | 3 | 3 | corces_2020,domcke_2020,trevino_2021 |
| BHLH::NFI_4 | | | 0.8964488041 | NFIX_M03476_2.00 | | 0.907458147 | NFIB.H12CORE.1.PS.A | | 4 | 1505 | 4 | 2 | corces_2020,trevino_2021 |
| ETV_2 | | | 0.9568373226 | ETV5_M04801_2.00 | | 0.8269094657 | FLI1.H12CORE.1.P.B | | 1 | 1487 | 1 | 1 | trevino_2021 |
| TCF7L-LEF_1 | | | 0.9746010484 | TCF7L1_M09395_2.00 | | 0.9714856184 | TF7L1.H12CORE.0.PM.A | | 1 | 1475 | 1 | 1 | domcke_2020 |
| NFI::NDF-ATOH | | | 0.7448160767 | CREB3L1_M04320_2.00 | | 0.704323574 | NFIA.H12CORE.0.P.B | | 2 | 1436 | 2 | 1 | trevino_2021 |
| MAF | | | 0.9291268684 | MAFA_M04356_2.00 | | 0.9429743186 | MAFG.H12CORE.1.PSM.A | | 1 | 1431 | 1 | 1 | corces_2020 |
| SOX::SOX_10 | | | 0.8443739927 | KMT2A_M01912_2.00 | | 0.7882904766 | E2F4.H12CORE.0.P.B | | 1 | 1400 | 1 | 1 | corces_2020 |
| NFI_3 | | | 0.8703958667 | NFIC_M09635_2.00 | | 0.8528648918 | PBX1.H12CORE.2.P.B | | 1 | 1391 | 1 | 1 | trevino_2021 |
| TBR1_2 | | | 0.8502843932 | TBX21_M09425_2.00 | | 0.824269399 | MEIS2.H12CORE.2.SM.B | | 3 | 1367 | 3 | 2 | corces_2020,trevino_2021 |
| ETS::SP-KLF | | | 0.8274532372 | ELK3_M04730_2.00 | | 0.8333267403 | ETS2.H12CORE.0.S.C | | 18 | 1357 | 18 | 2 | corces_2020,trevino_2021 |
| RFX_4 | | | 0.9120219855 | RFX2_MA0600.1 | | 0.9036312343 | RFX3.H12CORE.0.PSM.A | | 1 | 1351 | 1 | 1 | corces_2020 |
| POU_5 | | | 0.9414325593 | FOSB::JUNB_MA1135.1 | | 0.8959423494 | MEIS1.H12CORE.0.P.B | | 1 | 1280 | 1 | 1 | trevino_2021 |
| SOX | | | 0.9772998055 | SRY_M11332_2.00 | | 0.983855993 | SOX12.H12CORE.0.SM.B | | 1 | 1238 | 1 | 1 | trevino_2021 |
| SOX::POU_2 | | | 0.8809061853 | POU2F3_MA0627.2 | | 0.8536723493 | VENTX.H12CORE.1.S.C | | 1 | 1237 | 1 | 1 | trevino_2021 |
| LHX_2 | | | 0.8838247337 | VSX2_M05039_2.00 | | 0.9752483203 | PO2F3.H12CORE.0.PS.A | | 1 | 1230 | 1 | 1 | corces_2020 |
| POU4F_1 | | | 0.8541906169 | LMX1B_M05154_2.00 | | 0.8347024281 | HME2.H12CORE.0.SM.B | | 1 | 1165 | 1 | 1 | corces_2020 |
| NFI_4 | | | 0.7872445478 | ZNF223_M07654_2.00 | | 0.7273687914 | PBX1.H12CORE.2.P.B | | 1 | 1141 | 1 | 1 | trevino_2021 |
| SOX::SOX_11 | | | 0.9100338226 | SOX10_M09387_2.00 | | 0.9267555661 | SOX8.H12CORE.0.PSM.A | | 1 | 1136 | 1 | 1 | corces_2020 |
| POU4F_2 | | | 0.9019242236 | NKX2-5_M10743_2.00 | | 0.9005910072 | PO3F3.H12CORE.2.S.B | | 2 | 1069 | 2 | 2 | domcke_2020,trevino_2021 |
| POU6F_1 | | | 0.8947642123 | DLX3_M01503_2.00 | | 0.8791556602 | PAX4.H12CORE.0.SM.B | | 1 | 1066 | 1 | 1 | corces_2020 |
| POU6F_2 | | | 0.9802253178 | POU6F2_M03303_2.00 | | 0.97310362 | PO6F1.H12CORE.0.SM.B | | 1 | 1061 | 1 | 1 | trevino_2021 |

| | | | | | | | | | | | | | |
|---|---|---|---|---|---|---|---|---|---|---|---|---|---|
| NFI_5 | | | 0.8838961179 | NFIX_M03476_2.00 | | 0.8976747109 | NFIX.H12CORE.1.S.B | | 2 | 1002 | 2 | 1 | corces_2020 |
| BHLH::NFI_6 | | | 0.8923743345 | NFIX_MA0671.1 | | 0.888346204 | NFIB.H12CORE.1.PS.A | | 1 | 1000 | 1 | 1 | trevino_2021 |
| POU_6 | | | 0.8357788512 | POU2F1_M10835_2.00 | | 0.7994785585 | PO3F2.H12CORE.2.SM.B | | 2 | 972 | 2 | 1 | trevino_2021 |
| POU4F_3 | | | 0.9414141901 | POU2F2_M05450_2.00 | | 0.837168963 | PO2F2.H12CORE.2.S.B | | 2 | 897 | 2 | 1 | trevino_2021 |
| POU3F_2 | | | 0.9581494538 | POU1F1_M05454_2.00 | | 0.78982937 | PIT1.H12CORE.1.S.B | | 1 | 875 | 1 | 1 | trevino_2021 |
| MEF2::NDF-ATOH | | | 0.8886247697 | MSC_M04188_2.00 | | 0.8612588038 | TCF21.H12CORE.1.SM.B | | 1 | 862 | 1 | 1 | trevino_2021 |
| NFI_6 | | | 0.9084192064 | NFIX_M03476_2.00 | | 0.9089754639 | NFIX.H12CORE.1.S.B | | 1 | 838 | 1 | 1 | trevino_2021 |
| POU_7 | | | 0.8390627845 | POU3F1_M09221_2.00 | | 0.8492438962 | PO3F1.H12CORE.0.P.B | | 1 | 808 | 1 | 1 | domcke_2020 |
| SOX::SOX_14 | | | 0.9343573269 | SOX9_M09389_2.00 | | 0.9177788114 | SOX8.H12CORE.0.PSM.A | | 1 | 794 | 1 | 1 | corces_2020 |
| POU_8 | | | 0.8907273219 | POU5F1_M09222_2.00 | | 0.9204138929 | PO3F3.H12CORE.1.P.C | | 1 | 780 | 1 | 1 | trevino_2021 |
| TFAP2 | | | 0.9670608686 | TFAP2A_M04054_2.00 | | 0.9592383995 | AP2A.H12CORE.0.PSM.A | | 1 | 775 | 1 | 1 | trevino_2021 |
| BHLH_4 | | | 0.8243040042 | ZNF93_MA1721.1 | | 0.7942063689 | MYF5.H12CORE.0.P.B | | 1 | 772 | 1 | 1 | corces_2020 |
| NFI_7 | | | 0.9140291182 | NFIC_M09636_2.00 | | 0.8298864711 | NFIX.H12CORE.0.SM.B | | 8 | 757 | 7 | 2 | corces_2020,trevino_2021 |
| E2F_1 | | | 0.9073394085 | KMT2A_M01912_2.00 | | 0.7974470424 | E2F1.H12CORE.1.S.B | | 1 | 747 | 1 | 1 | domcke_2020 |
| POU_9 | | | 0.9410462862 | POU2F2_M07974_2.00 | | 0.9270544939 | PO3F3.H12CORE.1.P.C | | 1 | 744 | 1 | 1 | trevino_2021 |
| BHLH_5 | | | 0.7810120307 | ZNF93_MA1721.1 | | 0.7759561802 | KMT2B.H12CORE.0.P.B | | 1 | 737 | 1 | 1 | corces_2020 |
| NFI_8 | | | 0.8705604075 | NFIX_M03476_2.00 | | 0.8793421689 | NFIA.H12CORE.0.P.B | | 3 | 728 | 3 | 2 | corces_2020,trevino_2021 |
| FOX::MEIS | | | 0.9228327903 | MEIS1_M10709_2.00 | | 0.9319645648 | PBX1.H12CORE.2.P.B | | 1 | 728 | 1 | 1 | trevino_2021 |
| AP1::NFAT | | | 0.8133831031 | ZKSCAN5_MA1652.1 | | 0.7688842041 | BATF3.H12CORE.0.P.B | | 1 | 724 | 1 | 1 | trevino_2021 |
| BHLH_6 | | | 0.9369285929 | TFAP4_M09803_2.00 | | 0.8727487316 | TBX4.H12CORE.0.PS.A | | 1 | 712 | 1 | 1 | trevino_2021 |
| NFI_9 | | | 0.8496570026 | NFIX_M03476_2.00 | | 0.8495288205 | NFIA.H12CORE.0.P.B | | 1 | 711 | 1 | 1 | corces_2020 |
| RFX_5 | | | 0.9058601292 | RFX3_MA0798.3 | | 0.8572657701 | RFX1.H12CORE.1.PSM.A | | 1 | 711 | 1 | 1 | corces_2020 |
| ETS::ETS_2 | | | 0.9406452946 | ETV6_M02990_2.00 | | 0.8079614115 | IRF4.H12CORE.0.P.B | | 1 | 703 | 1 | 1 | trevino_2021 |

| | | | | | | | | | | | | | |
|---|---|---|---|---|---|---|---|---|---|---|---|---|---|
| NDF-ATOH_3 | | | 0.8524609393 | POU2F2_M05447_2.00 | | 0.870286158 | NDF1.H12CORE.1.S.C | | 1 | 686 | 1 | 1 | trevino_2021 |
| E2F_3 | | | 0.7393840446 | KMT2A_M01912_2.00 | | 0.7396896909 | E2F4.H12CORE.0.P.B | | 1 | 670 | 1 | 1 | domcke_2020 |
| MEF2_3 | | | 0.9176041883 | MEF2A_M08212_2.00 | | 0.8057186025 | MEF2A.H12CORE.0.P.B | | 1 | 668 | 1 | 1 | domcke_2020 |
| POU2F_2 | | | 0.9083068896 | POU1F1_M03300_2.00 | | 0.9331512397 | PO3F2.H12CORE.0.P.B | | 2 | 656 | 2 | 1 | trevino_2021 |
| SP-KLF_3 | | | 0.8919406946 | KLF1_M00224_2.00 | | 0.8808109637 | SP4.H12CORE.0.P.C | | 4 | 652 | 4 | 3 | corces_2020,domcke_2020,trevino_2021 |
| TCF7L-LEF_2 | | | 0.9779431922 | TCF4_M09461_2.00 | | 0.9588014902 | TF7L2.H12CORE.0.P.B | | 1 | 652 | 1 | 1 | trevino_2021 |
| ETS::ATF_2 | | | 0.8609625014 | ELK1::SREBF2_MA1933.1 | | 0.7723147536 | ETV3.H12CORE.0.SM.B | | 1 | 641 | 1 | 1 | trevino_2021 |
| PAX | | | 0.8351488496 | PAX6_M03291_2.00 | | 0.8197710718 | PAX6.H12CORE.0.PSM.A | | 1 | 636 | 1 | 1 | trevino_2021 |
| LHX::LHX_3 | | | 0.9107052972 | EN1_M03181_2.00 | | 0.8743044587 | SP9.H12CORE.0.P.B | | 4 | 631 | 4 | 2 | domcke_2020,trevino_2021 |
| ETS_3 | | | 0.8196216364 | GMEB2_M05736_2.00 | | 0.8103345384 | ELK4.H12CORE.0.PSM.A | | 5 | 611 | 5 | 2 | corces_2020,domcke_2020 |
| MEF2_4 | | | 0.8296548644 | MEF2A_M08212_2.00 | | 0.7735453212 | MEF2C.H12CORE.1.SM.B | | 1 | 597 | 1 | 1 | corces_2020 |
| CTCF_2 | | | 0.9011357283 | ZNF2_M07780_2.00 | | 0.8595958287 | E2F1.H12CORE.1.S.B | | 1 | 594 | 1 | 1 | corces_2020 |
| SPI | | | 0.907374744 | SPIB_M09537_2.00 | | 0.8727621627 | SPI1.H12CORE.0.P.B | | 2 | 584 | 2 | 1 | corces_2020 |
| HSF1 | | | 0.8639571188 | HSF1_M07977_2.00 | | 0.8694334215 | HSF1.H12CORE.2.P.B | | 5 | 581 | 5 | 2 | corces_2020,domcke_2020 |
| NFI_10 | | | 0.8339891483 | ESR1_MA0112.2 | | 0.8208888938 | PBX1.H12CORE.2.P.B | | 1 | 568 | 1 | 1 | corces_2020 |
| LHX::NFI | | | 0.8781062717 | NOTO_MA0710.1 | | 0.8751494212 | HME2.H12CORE.0.SM.B | | 4 | 554 | 4 | 2 | domcke_2020,trevino_2021 |
| SP-KLF_4 | | | 0.8052082077 | KLF1_M00224_2.00 | | 0.8533882366 | SP4.H12CORE.0.P.C | | 1 | 543 | 1 | 1 | corces_2020 |
| BHLH_7 | | | 0.9267243909 | ZNF93_MA1721.1 | | 0.8206603154 | HEN2.H12CORE.0.SM.B | | 1 | 537 | 1 | 1 | corces_2020 |
| LHX::LHX_4 | | | 0.9322112293 | LHX2_M03107_2.00 | | 0.8257632021 | LHX2.H12CORE.1.S.C | | 1 | 528 | 1 | 1 | trevino_2021 |
| LHX::LHX_5 | | | 0.9223196195 | LMX1B_M05154_2.00 | | 0.8890585868 | DLX2.H12CORE.1.S.B | | 1 | 526 | 1 | 1 | trevino_2021 |
| EGR_2 | | | 0.8764224514 | EGR1_M04438_2.00 | | 0.8253946653 | EGR3.H12CORE.0.PSM.A | | 1 | 526 | 1 | 1 | corces_2020 |
| NFI_11 | | | 0.9078871355 | NFIX_M03476_2.00 | | 0.9041565811 | NFIB.H12CORE.1.PS.A | | 1 | 520 | 1 | 1 | trevino_2021 |
| MEF2_5 | | | 0.8089137754 | MEF2A_MA0052.2 | | 0.8513505037 | MEF2C.H12CORE.0.P.B | | 1 | 519 | 1 | 1 | corces_2020 |

| | | | | | | | | | | | | | |
|---|---|---|---|---|---|---|---|---|---|---|---|---|---|
| ZEB-SNAI::ZEB-SNAI | 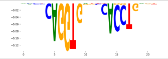 | 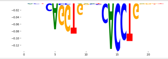 | 0.8362471474 | ZEB1_M10093_2.00 | 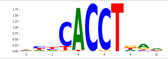 | 0.8345477633 | ZEB2.H12CORE.0.P.B | 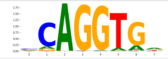 | 8 | 514 | 8 | 3 | corces_2020,domcke_2020,trevino_2021 |
| ETS_4 | 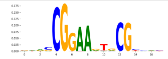 | 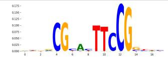 | 0.8162552603 | GMEB1_M05738_2.00 | 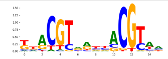 | 0.8064474896 | ELK4.H12CORE.0.PSM.A | 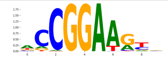 | 2 | 511 | 2 | 2 | corces_2020,domcke_2020 |
| ZBTB7 | 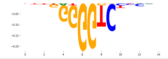 | 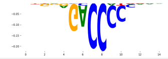 | 0.9598456518 | ZBTB7A_M07937_2.00 | 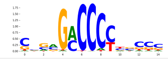 | 0.9623714712 | ZBT7A.H12CORE.0.P.B | 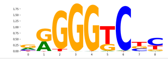 | 4 | 501 | 4 | 1 | corces_2020 |
